# Projecting climate-driven shifts in demersal fish habitat in Iceland’s waters

**DOI:** 10.1101/2021.03.04.433927

**Authors:** Julia G. Mason, Pamela J. Woods, Magnús Thorlacius, Kristinn Guðnason, Vincent S. Saba, Patrick J. Sullivan, Kristin M. Kleisner

## Abstract

As climate change shifts marine species distribution and abundance worldwide, projecting local changes over decadal scales may be a valuable adaptive strategy for managers and industry. In Iceland, one of the top fish-producing nations in the world, long-term monitoring enables model simulations of groundfish species habitat distribution. We used generalized additive models to characterize suitable thermal habitat for 47 fish species in Iceland’s waters. We then projected changes in thermal habitat by midcentury with an ensemble of five general circulation models from the Coupled Model Intercomparison Program 6 (CMIP6) and NOAA (CM2.6) and two scenarios (SSP 5-8.5 and SSP 2-4.5). We find a general northward shift in centroids of habitat distribution, with variable regional dynamics among species. Species thermal affinity was the most significant predictor of future habitat change, with warmer-water species more likely to see projected increases in suitable habitat. We present spatially explicit habitat change projections for commercially and culturally important species. These projections might serve as guideposts to inform long-term management decisions about regional and species-specific suitability for Iceland’s fisheries, infrastructure investment, and risk evaluation under climate change.

## Introduction

Climate change is shifting marine species distribution and abundance worldwide (Parmesan, 2006; Poloczanska et al., 2013). These shifts are projected to continue as climate change intensifies (Molinos et al., 2016), with complex and unevenly distributed social-ecological consequences (Golden et al., 2016; Lam et al., 2016; Pinsky et al., 2018; Sumaila et al., 2011). Alongside the urgent need for bold action to reduce greenhouse gas emissions is the need for adaptive management approaches to maintain desired fishery outcomes under changing and novel conditions. Researchers project that implementing management that accounts for changes in fish species productivity and distribution can improve outcomes for fishery catches and profits under most climate scenarios (Free et al., 2020; Gaines et al., 2018). For fisheries with adequate scientific and technical capacity, conducting forecasts and incorporating future climate scenarios into management decisions is a key aspect of climate-adaptive management (Free et al., 2020; Holsman et al., 2019; Karp et al., 2019; Pinsky & Mantua, 2014). While seasonal and interannual forecasts may be of most immediate use for management and industry, climate-scale projections allow managers and practitioners to evaluate risk, plan for future losses or gains in suitability, and inform longer term decision-making processes such as national legislation or international negotiations (Hobday et al., 2018; Holsman et al., 2019).

In Iceland, the economic and cultural importance of fisheries and the sophistication of its management and scientific systems mean that projecting future change in fish stocks could be a key priority for achieving climate-adaptive fisheries (Kleisner et al. in press). Consistently ranking among the top twenty marine fish capture producing countries worldwide (FAO, 2020), Iceland is located in a highly productive transition zone between warm Atlantic and cold Polar currents. The interactions of these currents create high spatial and temporal oceanographic variability within Iceland’s waters, and the ecosystem is highly sensitive to their dynamics (Astthorsson et al., 2007). Environmental conditions have fluctuated over the past century in relation to multidecadal oscillations and local atmospheric dynamics with profound ecological and economic consequences: Relatively warm periods were associated with fishery booms that drove the development of Iceland’s commercial fisheries and economy, and cooler-water periods (in combination with overfishing) associated with devastating fishery crashes (Astthorsson et al., 2007; Ogilvie & Jónsdóttir, 2000; Valtýsson & Jónsson, 2018).

Recently, a warm anomaly from the mid-1990s to late 2010s drove substantial changes in fish abundance and distribution around Iceland, including documentation of new species and increases of warmer-water species (Astthorsson & Palsson, 2006; Valdimarsson et al., 2012; Valtýsson & Jónsson, 2018). Particularly notorious was the abrupt expansion of Atlantic mackerel (*Scomber scombrus*) into Iceland’s waters (Olafur S. Astthorsson et al., 2012; Olafsdottir et al., 2019) that quickly became an economic boon for Iceland, but led to international political conflict over the fishing of the stock (Spijkers & Boonstra, 2017). Researchers also noted significant shifts in groundfish distribution and community assemblage, with heterogeneous regional trends associated with oceanographic conditions and the influence of coastal fjord systems on predator-prey dynamics (Jónsdóttir et al., 2019; Stefánsdóttir, 2008; Stefánsdóttir, 2019).

Given these past temperature-related changes, a logical next step might be to project how future change might affect fish species abundance and distribution. Global studies have projected that Iceland, like other high-latitude countries, could be a climate “winner,” potentially experiencing increased biodiversity and fisheries catch potential as warmer waters move fish poleward (Cheung et al., 2009; Molinos et al., 2016). However, given the highly local and variable dynamics of Iceland climatic conditions and differing ecological, economic, and cultural importance of its fish species, higher-resolution projections indicating specific species and regional dynamics could be more applicable for fisheries managers and industry. Campana et al. (2020) used Iceland bottom trawl survey data to model fish habitat and projected that a uniform 1°C increase in bottom temperature would drive a general northward shift in habitat distribution, with significant variation across species and quadrants of the exclusive economic zone (EEZ). The authors noted a high level of regional and temporal variation in past warming trends and warned that future warning is unlikely to be homogenous nor linear.

Iceland’s fishery managers have collected standardized fisheries independent and fisheries dependent data since the 1980s, allowing for detailed analyses of species abundance and distribution in relation to environmental conditions that can inform future projections. These long-term monitoring data are ideal for a common but data-intensive approach to anticipating regional or global species distribution shifts: pairing statistical models of species suitable habitat with global climate model outputs (Stock et al., 2011). Given the uncertainty inherent to projecting both the dynamics of the global climate and the human actions and policies that influence those dynamics, using an ensemble of climate models across different scenarios is advised (Morley et al., 2018, 2020). Here we use long-term fisheries independent trawl data and an ensemble of the newest generation of global climate models from the Coupled Model Intercomparison Project (CMIP6) to project how future climate change will affect suitable thermal habitat of 47 species in Iceland’s waters at a 0.25° x 0.25° resolution. These spatially explicit climate projections can more directly inform Iceland’s fishing industry and fisheries management’s needs for adapting to climate-driven changes in fish distribution, and illustrate to other nations and regions how these projections might be considered in long-term climate-adaptive management.

## Methods

### Projecting future temperature changes in Iceland’s waters

We used a suite of global climate models to project future ocean surface and bottom temperatures in Iceland’s waters, including a high-resolution global climate model (CM2.6 from the National Oceanic and Atmospheric Administration Geophysical Fluid Dynamics Laboratory, NOAA GFDL) and four coarser global climate models from the Coupled Model Intercomparison Project 6 (CMIP6). Researchers have found that CM2.6 (10-km ocean resolution) resolves the ocean circulation in the Northwest Atlantic more realistically than coarser models (Saba et al., 2016). However, this model has a cold bias in sea surface and bottom temperature in Iceland’s waters in the historical period from 1982-2012, so we selected the highest resolution CMIP6 models that have a range of cold and warm biases to complement the higher resolution CM2.6. These were GFDL CM4 (25-km ocean resolution) from the National Oceanic and Atmospheric Administration Geophysical Fluid Dynamics Laboratory, U.S.A; CNRM-CM6 (25-km ocean resolution) from the Centre National de Recherches Meteorologiques and Centre Europeen de Recherche et de Formation Avancee en Calcul Scientifique, France; HadGEM3-GC31 (100-km ocean resolution) from the Met Office Hadley Centre, U.K.; and IPSL-CM6A-LR (100-km ocean resolution) from L’Institut Pierre Simon Laplace, France. We downloaded CMIP6 data using the xarray package (version 0.15.1) in Python (version 3.7.6) (Hoyer & Hamman, 2017).

The CMIP6 models use future scenario simulations that combine the Representative Concentration Pathways (RCPs) of radiative forcings used in the Intergovernmental Panel on Climate Change’s Fifth Assessment Report (IPCC AR5) with Shared Socioeconomic Pathways (SSPs) of societal development (Eyring et al., 2016; O’Neill et al., 2017). We examined two future scenarios for the years 2015-2100: SSP 2-4.5, a “middle of the road” scenario where countries continue along historical social, economic, and technological development trajectories as they strive toward sustainable development goals; and SSP 5-8.5, a “fossil-fueled development” scenario where accelerated economic growth emphasizes fossil fuels (O’Neill et al., 2017). The CM2.6 projects an idealized transient climate response (1% per year increase in atmospheric CO_2_) simulation over 80 years, where the last 20 years are comparable to years 2061-2080 of SSP 5-8.5. We thus focused on this 2061-2080 period for our future projections.

We standardized the models using the “delta method,” where we calculated the difference between each model’s monthly projections and modeled historical control (mean of 1985-2015), and then added these deltas to a standard climatology (Anandhi et al., 2011; Kleisner et al., 2017; Morley et al., 2018). We used a sea surface temperature (SST) climatology from daily NOAA Optimum Interpolated Sea Surface Temperature (OISST) data from 1982-2012 and a bottom temperature climatology from the NOAA Greenland-Iceland-Norwegian Seas Regional Climatology version 2 (GINS RC v2) from 1985-2012 (Seidov et al. 2013). All projections were interpolated to a standard 0.25° x 0.25°grid to match the OISST resolution. We ran a monthly SST and bottom temperature hindcast for the years 2000-2018 using the Mercator Ocean Global Reanalysis (GLORYS) dataset from the Copernicus Marine Environment Monitoring Service and a projection for the years 2061-2080 for each the five models and two scenarios on a projection grid of the 2,312 0.25° cells in Iceland’s EEZ.

### Modeling species thermal habitat

The Icelandic Marine and Freshwater Research Institute (MFRI) conducts annual standardized bottom trawl surveys in the spring and autumn to inform groundfish stock assessments. The spring survey has been conducted since 1985 at about 590 fixed stations covering Iceland’s continental shelf to 500m depth (Solmundsson et al., 2020). The autumn survey, initiated in 1996 and expanded in 2000, covers about 400 fixed stations that additionally include deeper waters along the continental slope (400-1500m; Campana et al. 2020). Fish lengths and standard weight conversions are used to calculate biomass (kg) per nautical mile. Surface temperature and bottom temperature are also measured in situ. We used surveys through March 2020, excluding the autumn survey before it was expanded in 2000 and 2011 when a labor strike interrupted the survey, for n = 27,524 total survey tows (Figure 1). We combined the spring and autumn surveys to better account for the full thermal envelope each species experiences throughout the year (Kleisner et al., 2017).

**Figure 1:**
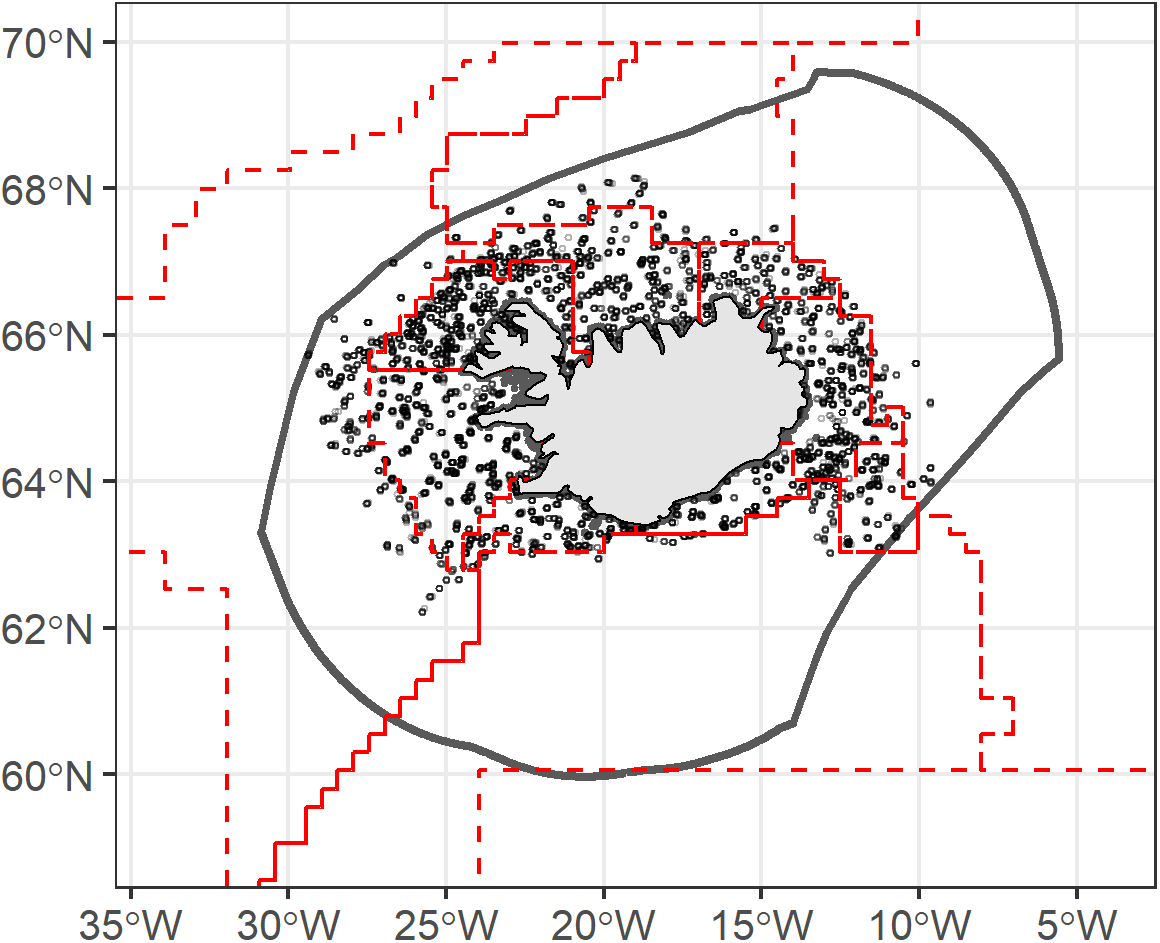
Study region indicating Iceland’s Exclusive Economic Zone (EEZ, solid line), Bormicon regions (red dashed lines), and bottom trawl survey points.

We modeled individual species distributions with two-stage generalized additive models (GAMs), using the mgcv package (version 1.8.33) in R (version 4.0.2) (Wood, 2011). The first stage was a presence-absence model with a binomial error distribution, and the second stage was a biomass given presence model using a log-link transform on non-zero observations with a gaussian error distribution. For both stages, we included a penalized likelihood to reduce model complexity (Morley et al., 2018; Wood, 2006). We calculated suitable thermal habitat as the product of the presence-absence model predictions, the back-transformed log-biomass model predictions, and a smearing estimate (the mean of the log-biomass model residuals) to correct for retransformation bias (Duan 1983; Kleisner et al., 2017; Morley et al., 2018). This suitable thermal habitat value does not directly represent species abundance, but rather the potential suitable habitat and species density solely based on model predictor variables (Kleisner et al., 2017; Morley et al., 2018).

The predictor variables for the models included tow and environmental information from the trawl surveys and more complex temperature variables from GLORYS. From the surveys, we used in situ surface and bottom temperature and the starting depth of the tow. We used GLORYS temperature data to characterize broader spatial and temporal aspects of thermal habitat that could influence species distribution, including the annual minimum and maximum surface and bottom temperature (that is, the minimum or maximum temperature over the preceding twelve months in the 0.25° × 0.25° cell where the tow was conducted) and spatial standard deviation of surface and bottom temperature (using a 3 × 3 window around each 0.25° × 0.25° cell) to approximate frontal activity (Abrahms et al., 2019; Morley et al., 2018), which is known to influence species distribution in Iceland’s waters (Astthorsson et al., 2007). We removed the annual minimum bottom temperature as a predictor variable because it was highly correlated with the survey in situ bottom temperature (Spearman’s rho = 0.77).

To account for habitat suitability factors not captured by depth and temperature, we added Bormicon regions (Figure 1), which are habitat zones used in MFRI’s multispecies modelling efforts (Stefansson, 2004), as a parametric term. These regions were delineated based on topography (the depth contour of the continental shelf and submarine ridges), hydrography (salinity distribution and mixing patterns), and general patterns of spawning, migration, and fishing effort for key demersal species (Stefansson & Palsson, 1997). We selected these regions based on their management relevance and in the absence of more detailed habitat quality data but recognize risks of tautology in using these regions to simulate species distribution. We performed additional tests of the suitability of this predictor during initial model selection and determined that it improved model performance in terms of prediction error, deviance explained, and Akaike’s Information Criterion (AIC) for the majority of species. The fact that there are relatively few survey tows in the offshore regions also limits predictive performance, so we focus our subsequent results and discussion on habitat dynamics along the continental shelf and provide the caveat that this variable should be interpreted with caution, particularly for any species for which models project offshore habitat changes.

Thus, the final model formulas were as follows, where s() represents a thin plate spline smooth term:

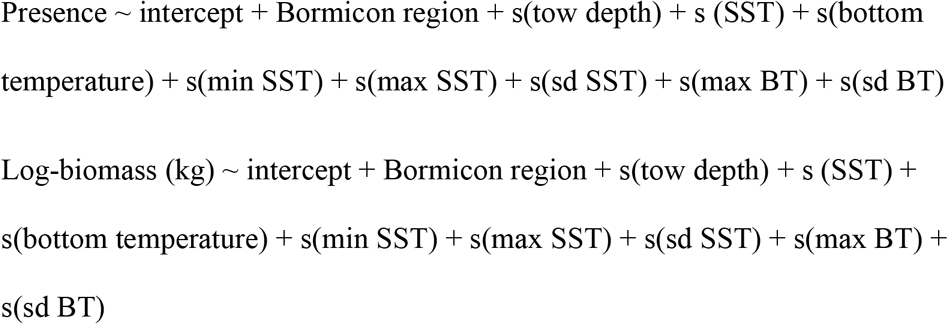

The GLORYS data were available from 1993-2018, so the final models were fitted with n = 19759 tows with all predictor data available. For species with no presences observed in a particular Bormicon region, we randomly replaced 10% of the tows in those regions with near-zero (1^−10^) biomass values to allow log-biomass projections in those areas, following Morely et al. (2018). Because these surveys were designed to sample demersal fish, we removed invertebrate and pelagic species for analysis. We ran models for 56 fish species with sufficient observations for the model parameters.

We assessed model performance by training the presence-absence models with tows through 2013 (approximately 80% of the tows), and testing predictions on the remaining 20% of the tows. We assessed presence-absence models using true skill statistic (TSS), a measure suited to spatial presence-absence prediction performance, where a score of 1 represents perfect agreement and a score < 0 is no better than random (Allouche et al., 2006). All species had a TSS score > 0.25, so we did not remove any species from analysis based on this metric. To evaluate the importance of each predictor variable in contributing to model explanatory power, we compared the percent deviance explained from the full model and a model with that covariate dropped for each species.

We considered the suitability of a temperature-based model by comparing the prediction error in the combined presence-absence and log-biomass thermal suitable habitat predictions with that of a “naïve” model with temperature variables removed (i.e., just depth and Bormicon region as predictors). Models were considered suitable if the ratio of the full model to naïve model error was < 1 and the Diebold-Mariano test p-value was < 0.05 (see Kleisner et al., 2017). We removed nine species from analysis based on this assessment and present results for the 47 remaining species. We performed these tests using the dismo package (version 1.3.2) in R (Hijmans et al., 2020). Other model results reported (deviance explained, suitable thermal habitat values) are based on the models fitted on all data, not split into training and testing. See supplemental information for model performance results for all species.

### Projecting future thermal habitat

We calculated the amount of available suitable habitat as the sum of all modeled thermal habitat values in the full EEZ projection grid (Morley et al., 2018). We compared the mean available habitat in the historical period (2000-2018) with the future period (2061-2080) for each species. Because some changes were quite large (orders of magnitude), we present log10 x-fold change (log (mean future thermal habitat / mean historical thermal habitat)) so that relative increases and decreases can be compared. We calculated the centroid of distribution for each species as mean latitude and longitude weighted by the modeled thermal habitat value, and calculated the distance (km) and direction (degrees) shifted between the historical and future period with the geosphere package (version 1.5.10) in R (Hijmans, 2019). Additionally, we calculated the shifts in warm and cold edges of species distributions as the difference in the 95^th^ and 5^th^ respective percentile latitude of thermal habitat values > 0.05 (Fredston-Hermann et al., 2020).

We grouped species by thermal affinity indices outlined in Campana et al. (2020), based on bottom temperatures and depths from the trawl surveys. We calculated thermal bias as the difference in median biomass-weighted bottom temperature for each species and the median bottom temperature of all survey tows, stenothermal index as the range of 5^th^ and 95^th^ percentiles of those biomass-weighted bottom temperatures for each species, and depth as the biomass-weighted median tow depth for each species. Because the spring and autumn surveys sample different areas and depths, we calculated separate index values for each season and took a weighted mean based on the number of observations in each season. Following Campana et al. (2020), we categorized species into warm water (positive thermal bias), cool water (−3°C to 0°C thermal bias), and cold water (< −3°C thermal bias) niches. We tested associations between thermal affinity indices and projected shifts in thermal habitat availability and distribution with ordinary least squares regression.

## Results

### Projected climate change

The projected climate model deltas resulted in a mean increase in surface temperatures in Iceland’s EEZ of 1.96°C for SSP 2-4.5 (ranging from 0.76°C, CNRM to 2.40°C, MOHC) and 2.40°C for SSP 5-8.5 (ranging from 1.00°C, CM2.6 to 3.40°C, IPSL) by 2061-2080. For bottom temperatures, the mean projected increase for 2061-2080 was 0.06°C for SSP 2-4.5 (ranging from −1.5°C, GFDL to 1.21°C, IPSL) and 0.20°C for SSP 5-8.5 (ranging from -1.35°C, GFDL to 1.47°C, IPSL) in 2061-2080 (see supplemental information for time series of individual climate model projections). Spatially, surface warming was projected throughout Iceland’s EEZ and most pronounced in offshore northeast waters for both scenarios (Figure 2a). The variation among climate model surface temperature projections was also highest in Iceland’s northern waters, which likely reflects uncertainty in the mixing dynamics between Atlantic and polar waters (Figure 2b). Bottom temperature deltas showed cooling in Iceland’s southern waters beyond the continental shelf, and concentrated warming along the northern shelf and east along the Iceland-Faroes ridge in the southeast (Figure 2c). Variation among models was highest in the east where warming is most pronounced, as well as offshore southern waters (Figure 2d).

**Figure 2:**
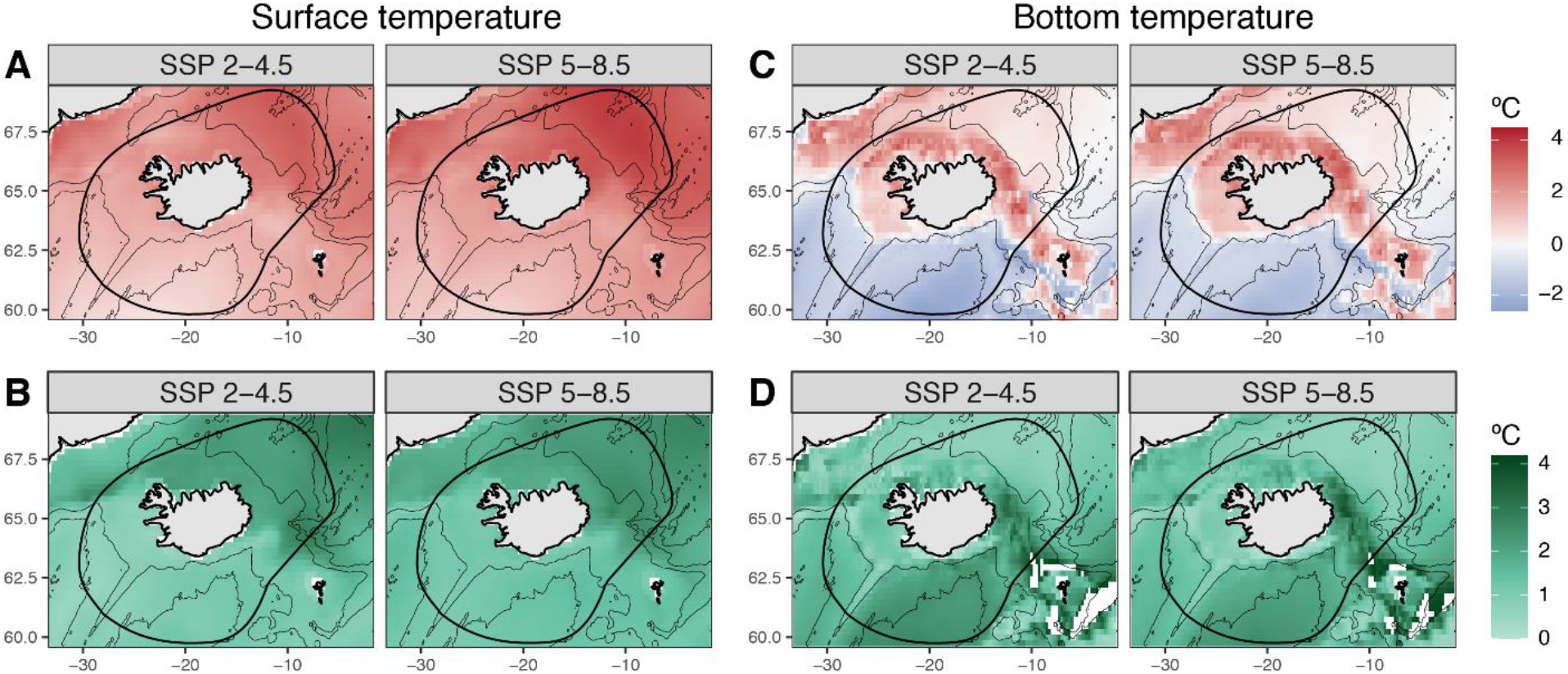
Projected ensemble means (A, C) and standard deviation (B, D) for sea surface and bottom temperature deltas for the years 2061-2080 for the middle of the road (SSP 2-4.5) and fossil-fueled development (SSP 5-8.5) scenarios. Thin lines indicate 1000m isobaths and thick lines indicate Iceland’s EEZ.

### Projected changes in suitable thermal habitat

The presence-absence GAMs explained mean 44% of deviance (sd 16; range 10-81) across the 47 species and had a mean TSS of 0.68 (sd 0.15, range 0.36-0.95). The log-biomass models explained mean 31% of deviance (sd 15, range 10-72). Tow depth and Bormicon region contributed the most to variance explained in both the presence-absence and log-biomass models. Models fitted without depth lost mean 12.9% variance explained for presence-absence (sd 9.11, range −2.05-32.1) and 5.5% for log-biomass (sd 4.64, range −0.05-16.3). Models fitted without Bormicon region lost mean 10.5% (sd 9.51, range −4.04-32.0) and 2.8% (sd 1.74, range −0.14 −7.02) variance explained in presence-absence and log-biomass models, respectively. For the presence-absence models, the remaining temperature variables contributed between 8.9-9.6% variance explained on average, whereas for log-biomass models, lost variance explained was minimal, ranging from 0.2% for SST standard deviation and maximum to 0.86% for bottom temperature.

Warmer water species were generally projected to see an increase in suitable thermal habitat in 2061-2080 relative to 2000-2018, while cold and cool water species were generally projected to see a decrease in habitat (Figure 3). The magnitude of decreases was higher than the magnitude of increases, and patterns were similar across the two scenarios. Thirteen species had a projected increase in habitat for all models for SSP 2-4.5 and fifteen for SSP 5-8.5, whereas fifteen species had a projected decrease in habitat for all models in SSP 2-4.5 and sixteen for SSP 5-8.5. Thermal bias was significantly positively correlated with median change in projected suitable habitat for SSP 5-8.5 (adjusted r^2^ = 0.12, p < 0.01; for SSP 2-4.5, adjusted r^2^ = 0.03 and p = 0.13), whereas stenothermic index and median depth did not show a significant relationship with projected suitable thermal habitat change in either scenario (stenothermic index: p = 0.13, 0.10; median depth: p = 0.12, 0.37).

**Figure 3:**
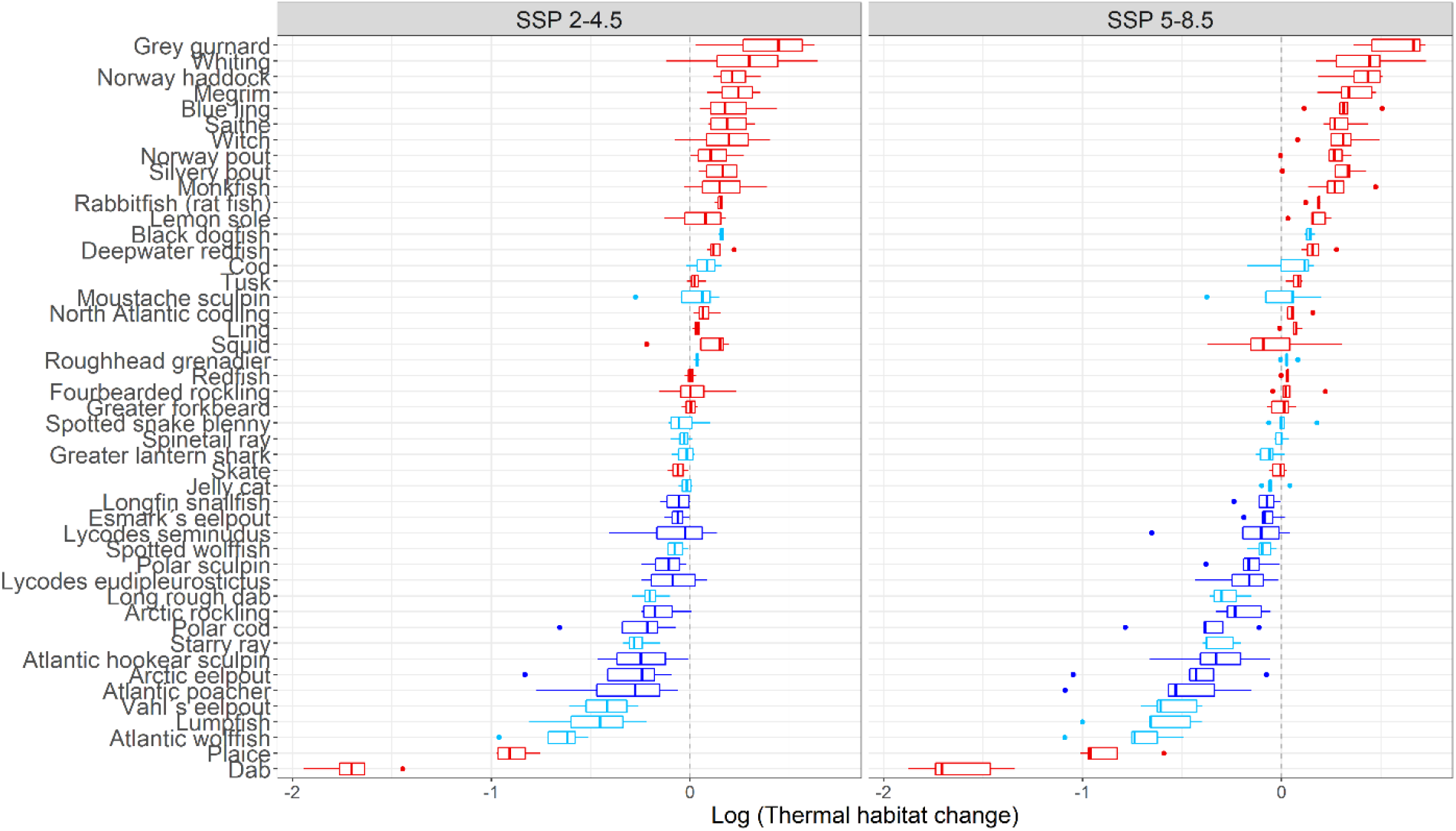
Projected changes in suitable thermal habitat for modeled species for 2061-2080 relative to 2000-2018 for middle of the road (SSP 2-4.5, left) and fossil-fueled development (SSP 5-8.5, right) scenarios. Boxplots represent the distribution of suitable habitat projections from each of the climate models. Colors represent species thermal niche: red species are warm water, light blue are cool water, and dark blue are cold water (see methods and Campana et al., 2020 for classifications). Note that x-axis is on a log10 scale, so a species with a value of −1 would have 10% of the total projected suitable thermal habitat in 2061-2080 relative to 2000-2018.

These projected habitat suitability shifts were spatially heterogeneous within Iceland’s waters, with most species showing relatively localized increases and/or decreases (Figure 4). Cod (*Gadus morhua*), saithe (*Pollachius virens*), and redfish (*Sebastes marinus*), key commercial species, showed increases along Iceland’s northern continental shelf. For cod, a cool water species, habitat suitability increases were more pronounced in the north and east, with potential suitability decreases in the southwest and southeast (Figure 4a). Redfish showed a pronounced decrease in southwest offshore waters. Long rough dab (*Hippoglossoides platessoides*), a cool water species, showed nearly the opposite pattern to cod, with decreases along the northern continental shelf and increases in the south/southeast (for SSP 2-4.5) (Figure 4b). Whiting (*Merlangius merlangus*), along with several other warm-water species including monkfish (*Lophius piscatorius*), witch (*Glyptocephalus cynoglossus*), megrim (*Lepidorhombus whiffiagonis*), and Norway haddock (*Sebastes viviparus*), showed habitat suitability increases along Iceland’s southern and western continental shelf (Figure 4c). Lumpfish (*Cyclopterus lumpus,* cool water), Atlantic wolffish (*Anarhichus lupus*, cool water), plaice (*Pleuronectes platessa*, warm water) and dab (*Limanda limanda*, warm water) all show inshore decreases, concentrated in the northwest fjords for lumpfish, wolffish, and dab and along the west coast bays for plaice (Figure 4d). Lemon sole (*Microstomus kitt*, warm water), on the other hand, shows nearshore increases around the island. Maps indicating projected habitat changes for all analyzed species and all projection periods are available in the supplemental information.

**Figure 4:**
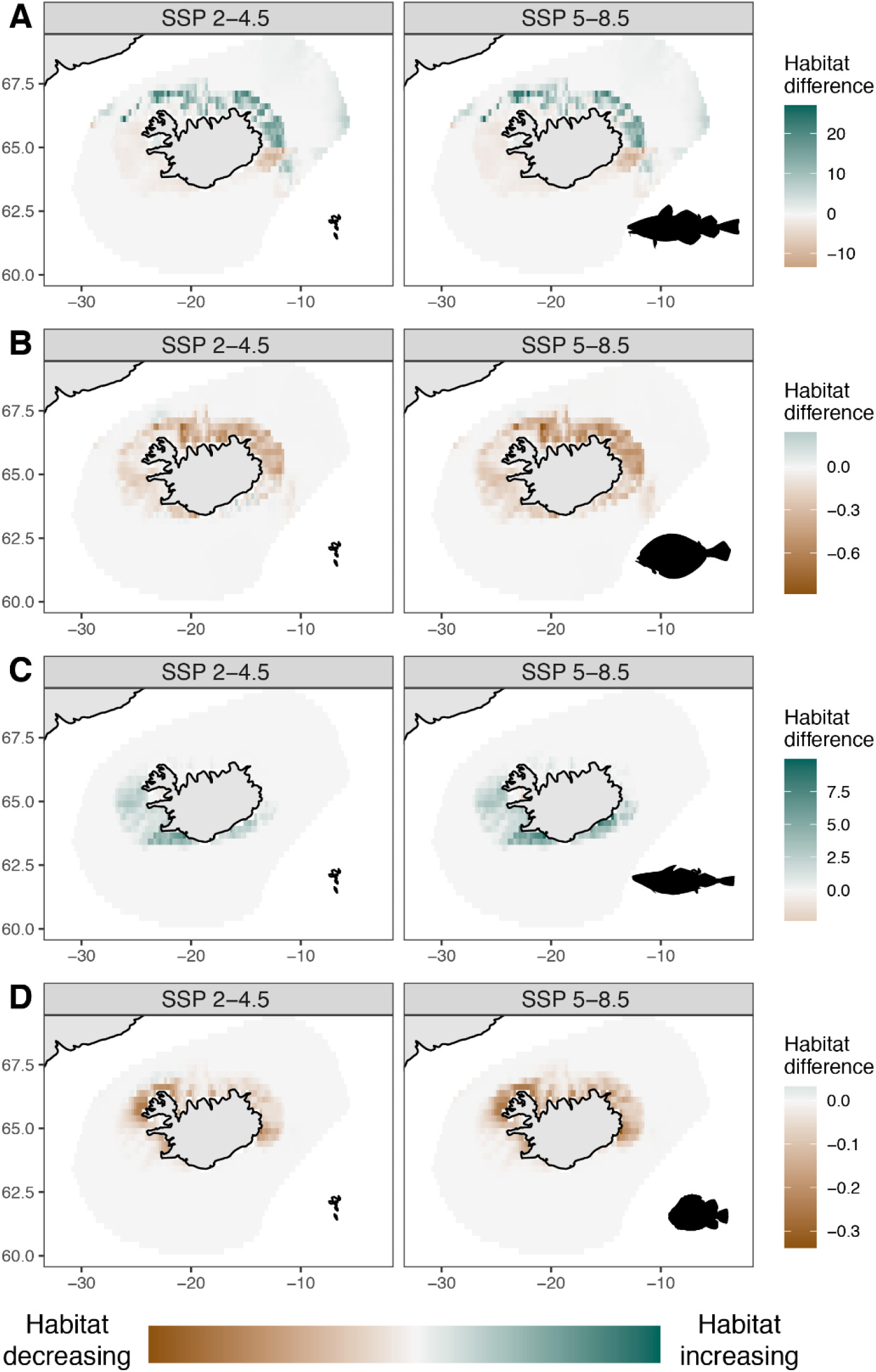
Projected changes in future habitat suitability for cod (A), long rough dab (B), whiting (C), and lumpfish (D) in 2061-2080 relative to 2000-2018 under the middle of the road (SSP 2-4.5, left) and fossil fueled development (SSP 5-8.5, right) scenarios. Green regions indicate increases in suitable habitat whereas brown regions indicate decreases in suitable habitat.

Taken together, these projected shifts in species distribution show general northward movement, trending north and northwest in SSP 2-4.5 and north and slightly northeast among warmer water species in SSP 5-8.5 (Figure 5). There was no significant relationship between thermal bias, stenothermic index, and depth with the distance or bearing of centroid change. Species with a higher thermal bias (i.e. warmer water species) were significantly more likely to see northward movement of the cold edge of their range (adj r^2^ = 0.10, p = 0.02 for SSP 2-4.5, adj r^2^ = 0.14, p = 0.005 for SSP 5-8.5), but there were no significant relationships observed for movement of the southern warm range edge or with other thermal affinity indices.

**Figure 5:**
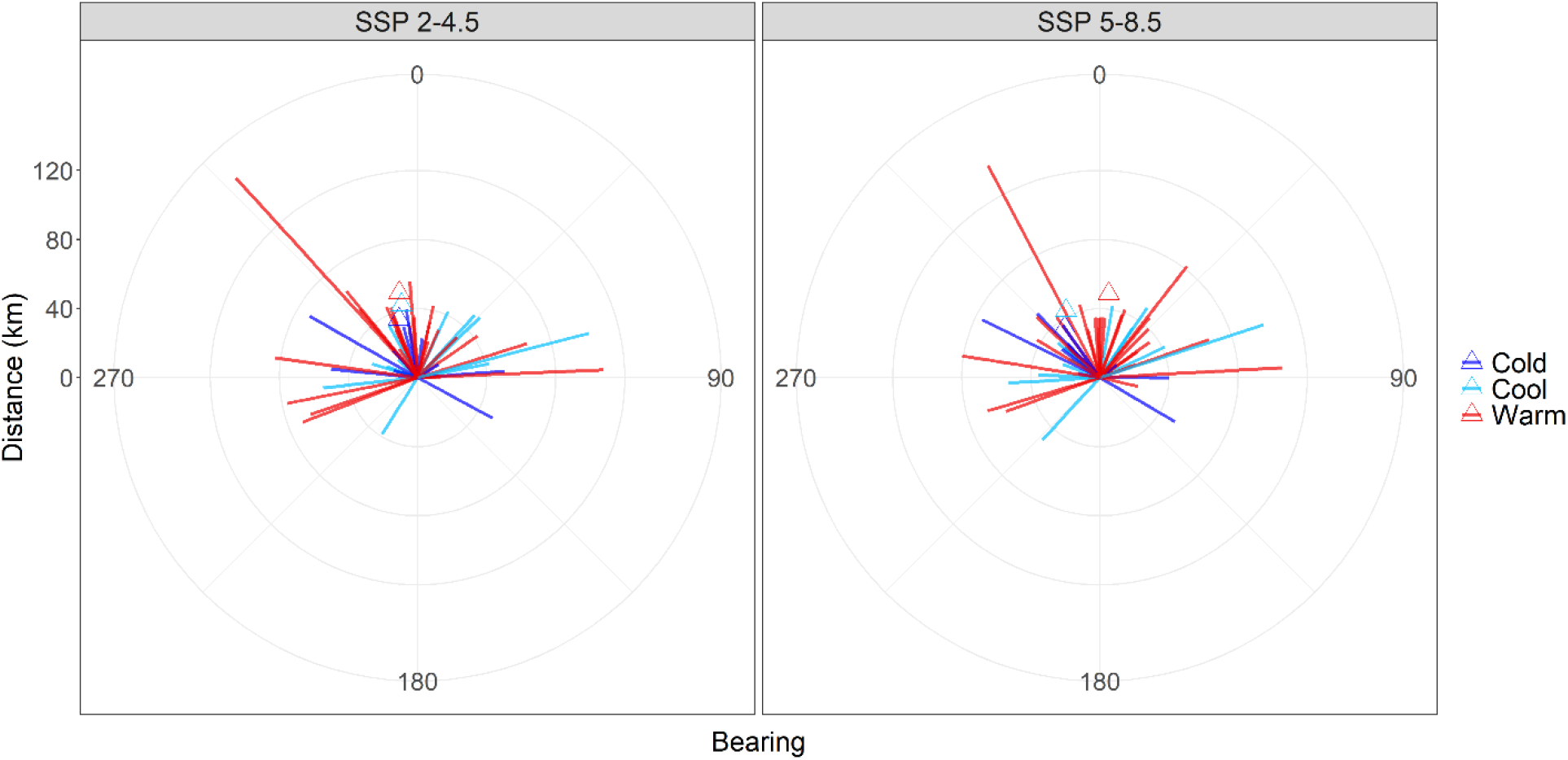
Projected distance (km) and direction (bearing, 0-360 degrees) of shifts in the centroid of suitable thermal habitat distribution between 2061-2080 and 2000-2018 under the middle of the road (SSP 2-4.5, left) and fossil fueled development (SSP 5-8.5, right) scenarios. Colors represent species thermal niche (see Figure 3). Triangles indicate the circular average value for the bearing change, grouped by thermal niche. Version with species labels available in the supplemental information.

## Discussion

These results corroborate previous projections that future climate change could result in significant shifts in fish species abundance and distribution in Iceland’s waters, and further illustrate potential variability in responses among species and regions. Consistent with global hypotheses and Campana et al. (2020)’s analyses, our results indicate an overall northward shift in center of biomass distribution for Iceland’s demersal fish species, with warmer water species more likely to expand the cold edge of their range. Species thermal bias was a more significant predictor of future habitat suitability change than stenothermic index or depth preference, results also similar to Campana et al. (2020)’s findings. In general, our analyses suggest similar trends to those observed during the mid-1990s warming period (Valtýsson & Jónsson, 2018) could be expected by midcentury, with many warmer water species likely to experience overall increases in suitable thermal habitat in Iceland’s EEZ. However, these patterns are localized and may be accompanied by regional decreases.

It is encouraging that many commercially important species are projected to see increases in suitable habitat in Iceland’s waters. However, regional heterogeneity in habitat suitability changes may be of interest to managers and the fishing industry. For example, the overall increase in habitat suitability for cod, a flagship commercial species for Iceland, is largely along the northern coast, where fishing capacity and infrastructure has declined following industry consolidation (Kokorsch & Benediktsson, 2018). Meanwhile, the capital region and western coast where cod catches have historically been highest may see decreases under the high-CO_2_ emission scenario. Fishers in these waters may see cod supplemented by warmer-water species such as monkfish and megrim. The impact of these shifts may be felt unevenly within the fishing industry as larger companies with more technological capacity are less dependent on proximity to fishing grounds than smaller companies (Edvardsson et al. 2018). Local small boat operations could be further disadvantaged by the projected decreased habitat suitability for several species in inshore areas, particularly in the fisheries-dependent northwest region. Declining inshore habitat suitability for lumpfish may be particularly challenging for communities where the fishery is a cultural mainstay (Chambers, 2016). The potential movement of lumpfish offshore may present an additional management concern if it increases lumpfish bycatch rates in other fisheries, but could be beneficial if it alleviates interactions with pinnipeds and seabirds. Thus, these multidecadal scale projections might be a consideration in long-term planning not only for fishery development but also broader infrastructure policy, as investment in energy transmission and roads is critical for the competitiveness of smaller and more remote processing facilities (Reynisson et al. 2012).

Further examination of the life history and ecology of these species may aid in interpreting the trends we present here and highlight any results that may warrant further, finer-scale investigation. For example, a potentially puzzling result is that the species with the largest projected suitable habitat decreases, dab and plaice, are both warm-water species. These species are found in shallow waters, and therefore may be particularly sensitive to temperature or limited in their habitat. Survey coverage and timing may also factor into the uncertainty of these results. For example, dab have an especially patchy distribution with few high-volume tows (MFRI, 2020), which likely reduces model predictive power. Plaice spawn during the early spring, concurrent with the spring surveys (Solmundsson et al., 2005), which may result in aggregations that could bias the survey. These spawning dynamics may in part explain why plaice show a different pattern from lemon sole, another warm and shallow water species that spawns later in the year. While our analyses were intended as an exploration of overall trends rather than fitting individual models for each species, a more tailored approach that incorporates these specific spawning dynamics would be advisable for any planning regarding these species. Similarly, a species-specific approach could reveal climate sensitivity in species we dropped from analysis based on MASE performance. For example, Greenland halibut appear to be sensitive to the interaction between bottom temperature and depth based on model performance in initial trials.

In addition to refining habitat models, more complex approaches are likely needed to accurately project future fish productivity and distribution in Iceland’s waters. Our estimates of suitable thermal habitat represent maximum potential abundances with all other factors held equal, and do not capture changes in physiology, species interactions, or fishing pressure, all of which could change in coming decades. Predator-prey interactions could be a dominant ecological driver in Iceland’s waters: During the mid-1990s warming period, increasing cod and other gadoid stocks along Iceland’s northern coast is believed to have triggered the decline of a commercially important shrimp fishery, as these fish preyed heavily on northern shrimp and other species in closed fjord systems (Jónsdóttir et al., 2019). More recently, mackerel predation has been associated with declines in other warm-water species despite continued warming (Valtýsson & Jónsson, 2018). Such predator-prey interactions, while not accounted for in our models, would be key management considerations under warming conditions.

Additionally, we did not differentiate size or age classes among species, nor other intraspecific population structures that may have distinct suitable thermal habitats and thus future climate-driven dynamics. Cod, for example, exhibit ontogenetic regional shifts that follow the clockwise flow of currents around Iceland (Astthorsson et al., 2007), as well as behavioral ecotypes with separate temperature and depth niches (Grabowski et al., 2011). Combining autumn and spring survey data in our models was a step toward reducing bias based on seasonal dynamics, but more specific models for species known to have age-differentiated temperature responses could provide more nuanced projections of how climate change could affect ecological interactions and fisheries productivity.

Another limitation of our study is that as the survey data are limited to Iceland’s EEZ, they may not capture the full thermal or depth range for these species. Thus, our characterizations of suitable habitat may be too restrictive, and our projections cannot account for potential migration of more southern stocks into Iceland’s waters. However, since many of Iceland’s demersal stocks are relatively constrained to Iceland’s continental shelf (Valtýsson & Jónsson, 2018), limiting this analysis to Iceland’s EEZ may be appropriate for the species discussed here. Expanding these models to include data from more southern surveys, such as the Western and Southern areas of the International Council for Exploration of the Sea International Bottom Trawl Survey, could more comprehensively capture species temperature niches and potential future climate responses. Such an analysis would be valuable for identifying dynamics of straddling stocks and informing priorities for international negotiations.

Finally, management decisions are likely to be more significant determinants of fish abundance and distribution than long-term climate trends, particularly for the next few decades (Mullon et al., 2016). Our habitat suitability projections can provide an idea of regions or species that could be important to develop or to protect, but the future outcomes will depend largely on human actions. This study demonstrates the value of using global climate models to make spatially explicit projections of fish suitable habitat, which may serve as guideposts for long-term scenario planning, investment in fisheries and infrastructure, and risk evaluation. For Iceland and other regions with available data to support habitat modeling, such forward-looking studies could be a valuable strategy for achieving climate-adaptive fisheries.

## Data statement

NOAA High Resolution SST data were provided by the NOAA/OAR/ESRL PSL, Boulder, Colorado, USA, from their website at https://www.ncdc.noaa.gov/oisst. GLORYS data were provided by the Copernicus Marine Environmental Monitoring Service at https://resources.marine.copernicus.eu/?option=com_csw&view=details&product_id=GLOBAL_REANALYSIS_PHY_001_031.

Code for downloading climate data and performing these analyses is available at https://github.com/juliagmason/iceland_climate_resilient_fisheries.

## Notes

https://github.com/juliagmason/iceland_climate_resilient_fisheries

### Competing Interest Statement

The authors have declared no competing interest.

## References

Abrahms, B., Welch, H., Brodie, S., Jacox, M. G., Becker, E. A., Bograd, S. J., Irvine, L. M., Palacios, D. M., Mate, B. R., & Hazen, E. L. (2019). Dynamic ensemble models to predict distributions and anthropogenic risk exposure for highly mobile species. Diversity and Distributions, 25(8), 1182–1193.

Allouche, O., Tsoar, A., & Kadmon, R. (2006). Assessing the accuracy of species distribution models: Prevalence, kappa and the true skill statistic (TSS). Journal of Applied Ecology, 43(6), 1223–1232.

Anandhi, A., Frei, A., Pierson, D. C., Schneiderman, E. M., Zion, M. S., Lounsbury, D., & Matonse, A. H. (2011). Examination of change factor methodologies for climate change impact assessment. Water Resources Research, 47(3). https://doi.org/10.1029/2010WR009104

Astthorsson, O., Gislason, A., & Jonsson, S. (2007). Climate variability and the Icelandic marine ecosystem. Deep Sea Research Part II: Topical Studies in Oceanography, 54(23–26), 2456–2477.

Astthorsson, O. S., & Palsson, J. (2006). New fish records and records of rare southern species in Icelandic waters in the warm period 1996–2005. ICES CM, 100, 20.

Astthorsson, Olafur S., Valdimarsson, H., Gudmundsdottir, A., & Óskarsson, G. J. (2012). Climate-related variations in the occurrence and distribution of mackerel (Scomber scombrus) in Icelandic waters. ICES Journal of Marine Science, 69(7), 1289–1297. https://doi.org/10.1093/icesjms/fss084

Chambers, C. P. (2016). FISHERIES MANAGEMENT AND FISHERIES LIVELIHOODS IN ICELAND. University of Alaska Fairbanks.

Cheung, W. W. L., Lam, V. W. Y., Sarmiento, J. L., Kearney, K., Watson, R., & Pauly, D. (2009). Projecting global marine biodiversity impacts under climate change scenarios. Fish and Fisheries, 10(3), 235–251. https://doi.org/10.1111/j.1467-2979.2008.00315.x

Duan, Naihua (1983). “Smearing Estimate: A Nonparametric Retransformation Method.” Journal of the American Statistical Association 78, no. 383: 605–10. https://doi.org/10.1080/01621459.1983.10478017.

Edvardsson, Kristinn Nikulás, Cezara Păstrăv, and Karl Benediktsson. “Mapping the Geographical Consolidation of Fishing Activities in Iceland during the Maturation of the ITQ Fisheries Management System.” Applied Geography 97 (August 2018): 85–97. https://doi.org/10.1016/j.apgeog.2018.05.013.

Eyring, V., Bony, S., Meehl, G. A., Senior, C. A., Stevens, B., Stouffer, R. J., & Taylor, K. E. (2016). Overview of the Coupled Model Intercomparison Project Phase 6 (CMIP6) experimental design and organization. Geoscientific Model Development, 9(5), 1937–1958. https://doi.org/10.5194/gmd-9-1937-2016

FAO. (2020). The State of World Fisheries and Aquaculture 2020 (Sustainability in Action). FAO. https://doi.org/10.4060/ca9229en

Fredston-Hermann, A., Selden, R., Pinsky, M., Gaines, S. D., & Halpern, B. S. (2020). Cold range edges of marine fishes track climate change better than warm edges. Global Change Biology, 26(5), 2908–2922. https://doi.org/10.1111/gcb.15035

Free, C. M., Mangin, T., Molinos, J. G., Ojea, E., Burden, M., Costello, C., & Gaines, S. D. (2020). Realistic fisheries management reforms could mitigate the impacts of climate change in most countries. PLOS ONE, 15(3), e0224347. https://doi.org/10.1371/journal.pone.0224347

Gaines, S. D., Costello, C., Owashi, B., Mangin, T., Bone, J., Molinos, J. G., Burden, M., Dennis, H., Halpern, B. S., Kappel, C. V., Kleisner, K. M., & Ovando, B. (2018). Improved fisheries management could offset many negative effects of climate change. Science Advances, 4(8), eaao1378.

Golden, C. D., Allison, E. H., Cheung, W. W. L., Dey, M. M., Halpern, B. S., McCauley, D. J., Smith, M., Vaitla, B., Zeller, D., & Myers, S. S. (2016). Nutrition: Fall in fish catch threatens human health. Nature, 534(7607), 317–320. https://doi.org/10.1038/534317a

Grabowski, T. B., Thorsteinsson, V., McAdam, B. J., & Marteinsdóttir, G. (2011). Evidence of Segregated Spawning in a Single Marine Fish Stock: Sympatric Divergence of Ecotypes in Icelandic Cod? PLOS ONE, 6(3), e17528. https://doi.org/10.1371/journal.pone.0017528

Hijmans, R. J. (2019). geosphere: Spherical Trigonometry. https://CRAN.R-project.org/package=geosphere

Hijmans, R. J., Phillips, S., Leathwick, J., & Elith, J. (2020). dismo: Species Distribution Modeling. https://CRAN.R-project.org/package=dismo

Hobday, A. J., Spillman, C. M., Eveson, J. P., Hartog, J. R., Zhang, X., & Brodie, S. (2018). A Framework for Combining Seasonal Forecasts and Climate Projections to Aid Risk Management for Fisheries and Aquaculture. Frontiers in Marine Science, 5. https://doi.org/10.3389/fmars.2018.00137

Holsman, K. K., Hazen, E. L., Haynie, A., Gourguet, S., Hollowed, A., Bograd, S. J., Samhouri, J. F., & Aydin, K. (2019). Towards climate resiliency in fisheries management. ICES Journal of Marine Science.

Hoyer, S., & Hamman, J. (2017). xarray: N-D labeled arrays and datasets in Python. Journal of Open Research Software, 5(1). https://doi.org/10.5334/jors.148

Jónsdóttir, I. G., Bakka, H., & Elvarsson, B. T. (2019). Groundfish and invertebrate community shift in coastal areas off Iceland. Estuarine, Coastal and Shelf Science, 219, 45–55.

Karp, M. A., Peterson, J. O., Lynch, P. D., Griffis, R. B., Adams, C. F., Arnold, W. S., Barnett, L. A. K., deReynier, Y., DiCosimo, J., Fenske, K. H., Gaichas, S. K., Hollowed, A., Holsman, K., Karnauskas, M., Kobayashi, D., Leising, A., Manderson, J. P., McClure, M., Morrison, W. E., … Link, J. S. (2019). Accounting for shifting distributions and changing productivity in the development of scientific advice for fishery management. ICES Journal of Marine Science, 76(5), 1305–1315. https://doi.org/10.1093/icesjms/fsz048

Kleisner, K. M., Fogarty, M. J., McGee, S., Hare, J. A., Moret, S., Perretti, C. T., & Saba, V. S. (2017). Marine species distribution shifts on the US Northeast Continental Shelf under continued ocean warming. Progress in Oceanography, 153, 24–36.

Kleisner, K. M., Ojea, E., Amoros, S., Battista, W., Burden, M., Cunningham, E., Fujita, R., Karr, K., Mason, J., Rader, D., Rovegno, N., Thomas-Smyth, A. (in press). Identifying policy approaches to build social-ecological resilience in marine fisheries with differing capacities and contexts. ICES Journal of Marine Science.

Kokorsch, M., & Benediktsson, K. (2018). Prosper or perish? The development of Icelandic fishing villages after the privatisation of fishing rights. Maritime Studies, 17(1), 69–83.

Lam, V. W. Y., Cheung, W. W. L., Reygondeau, G., & Sumaila, U. R. (2016). Projected change in global fisheries revenues under climate change. Scientific Reports, 6(1), 32607. https://doi.org/10.1038/srep32607

Marine and Freshwater Research Institute (2020). Technical report: Dab—Sandkoli (18 pp). https://www.hafogvatn.is/static/extras/images/27-dab_tr1206897.pdf

Molinos, J. G., Halpern, B. S., Schoeman, D. S., Brown, C. J., Kiessling, W., Moore, P. J., Pandolfi, J. M., Poloczanska, E. S., Richardson, A. J., & Burrows, M. T. (2016). Climate velocity and the future global redistribution of marine biodiversity. Nature Climate Change, 6(1), 83.

Morley, J. W., Frölicher, T. L., & Pinsky, M. L. (2020). Characterizing uncertainty in climate impact projections: A case study with seven marine species on the North American continental shelf. ICES Journal of Marine Science, fsaa103. https://doi.org/10.1093/icesjms/fsaa103

Morley, J. W., Selden, R. L., Latour, R. J., Frölicher, T. L., Seagraves, R. J., & Pinsky, M. L. (2018). Projecting shifts in thermal habitat for 686 species on the North American continental shelf. PLoS ONE, 13(5). https://doi.org/10.1371/journal.pone.0196127

Mullon, C., Steinmetz, F., Merino, G., Fernandes, J. A., Cheung, W. W. L., Butenschön, M., & Barange, M. (2016). Quantitative pathways for Northeast Atlantic fisheries based on climate, ecological–economic and governance modelling scenarios. Ecological Modelling, 320, 273–291. https://doi.org/10.1016/j.ecolmodel.2015.09.027

Ogilvie, A. E., & Jónsdóttir, I. (2000). Sea ice, climate, and Icelandic fisheries in the eighteenth and nineteenth centuries. Arctic, 53(4), 383–394.

Olafsdottir, A. H., Utne, K. R., Jacobsen, J. A., Jansen, T., Óskarsson, G. J., Nøttestad, L., Elvarsson, B. Þ., Broms, C., & Slotte, A. (2019). Geographical expansion of Northeast Atlantic mackerel (Scomber scombrus) in the Nordic Seas from 2007 to 2016 was primarily driven by stock size and constrained by low temperatures. Deep Sea Research Part II: Topical Studies in Oceanography, 159, 152–168. https://doi.org/10.1016/j.dsr2.2018.05.023

O’Neill, B. C., Kriegler, E., Ebi, K. L., Kemp-Benedict, E., Riahi, K., Rothman, D. S., van Ruijven, B. J., van Vuuren, D. P., Birkmann, J., Kok, K., Levy, M., & Solecki, W. (2017). The roads ahead: Narratives for shared socioeconomic pathways describing world futures in the 21st century. Global Environmental Change, 42, 169–180. https://doi.org/10.1016/j.gloenvcha.2015.01.004

Parmesan, C. (2006). Ecological and Evolutionary Responses to Recent Climate Change. Annual Review of Ecology, Evolution, and Systematics, 37(1), 637–669. https://doi.org/10.1146/annurev.ecolsys.37.091305.110100

Pinsky, M. L., Reygondeau, G., Caddell, R., Palacios-Abrantes, J., Spijkers, J., & Cheung, W. W. (2018). Preparing ocean governance for species on the move. Science, 360(6394), 1189–1191.

Pinsky, M., & Mantua, N. (2014). Emerging Adaptation Approaches for Climate-Ready Fisheries Management. Oceanography, 27(4), 146–159. https://doi.org/10.5670/oceanog.2014.93

Poloczanska, E. S., Brown, C. J., Sydeman, W. J., Kiessling, W., Schoeman, D. S., Moore, P. J., Brander, K., Bruno, J. F., Buckley, L. B., Burrows, M. T., Duarte, C. M., Halpern, B. S., Holding, J., Kappel, C. V., O’Connor, M. I., Pandolfi, J. M., Parmesan, C., Schwing, F., Thompson, S. A., & Richardson, A. J. (2013). Global imprint of climate change on marine life. Nature Climate Change, 3(10), 919–925. https://doi.org/10.1038/nclimate1958

Reynisson, Reinhard, Jona Matthiasdottir, Tryggvi Finnsson, and Ari Pall Palsson. “Northeast Iceland Infrastructure Analysis.” Husavik, Iceland: Northeast Iceland Development Agency, 2012.

Saba, V. S., Griffies, S. M., Anderson, W. G., Winton, M., Alexander, M. A., Delworth, T. L., Hare, J. A., Harrison, M. J., Rosati, A., & Vecchi, G. A. (2016). Enhanced warming of the Northwest Atlantic Ocean under climate change. Journal of Geophysical Research: Oceans, 121(1), 118–132.

Seidov, Dan; Baranova, Olga K.; Biddle, Mathew; Boyer, Tim P.; Johnson, Daphne R.; Mishonov, Alexey V.; Paver, Christopher R.; Zweng, Melissa M. (2013). Greenland-Iceland-Norwegian Seas Regional Climatology (NCEI Accession 0112824). NOAA National Centers for Environmental Information. Dataset. https://doi-org.proxy.library.cornell.edu/10.7289/v5gt5k30. Accessed 14-07-2020.

Solmundsson, J., Karlsson, H., Bjornsson, H., Jonsdottir, I. G., Jakobsdóttir, K., & Bogason, V. (2020). A manual for the Icelandic groundfish survey in spring 2020. Marine and Freshwater Research Institute.

Solmundsson, J., Palsson, J., & Karlsson, H. (2005). Fidelity of mature Icelandic plaice (Pleuronectes platessa) to spawning and feeding grounds. ICES Journal of Marine Science, 62(2), 189–200. https://doi.org/10.1016/j.icesjms.2004.11.012

Spijkers, J., & Boonstra, W. J. (2017). Environmental change and social conflict: The northeast Atlantic mackerel dispute. Regional Environmental Change, 17(6), 1835–1851. https://doi.org/10.1007/s10113-017-1150-4

Stefánsdóttir, L. (2008). Groundfish species diversity and assemblage structure in Icelandic waters during a period of rapid warming (1996-2007).

Stefánsdóttir, R. B. (2019). Climate change and northward distribution shifts in Icelandic groundfish species. University of Iceland.

Stefansson, G. (2004). Development of structurally detailed statistically testable models of marine populations (No. QLK5-CT1999-01609; p. 200).

Stefansson, G., & Palsson, O. K. (1997). BORMICON: A Boreal Migration and Consumption Model (No. 58; p. 233). Marine Research Institute. https://www.hafogvatn.is/static/research/files/fjolrit-058.pdf

Stock, C. A., Alexander, M. A., Bond, N. A., Brander, K. M., Cheung, W. W. L., Curchitser, E. N., Delworth, T. L., Dunne, J. P., Griffies, S. M., Haltuch, M. A., Hare, J. A., Hollowed, A. B., Lehodey, P., Levin, S. A., Link, J. S., Rose, K. A., Rykaczewski, R. R., Sarmiento, J. L., Stouffer, R. J., … Werner, F. E. (2011). On the use of IPCC-class models to assess the impact of climate on Living Marine Resources. Progress in Oceanography, 88(1), 1–27. https://doi.org/10.1016/j.pocean.2010.09.001

Sumaila, U. R., Cheung, W. W. L., Lam, V. W. Y., Pauly, D., & Herrick, S. (2011). Climate change impacts on the biophysics and economics of world fisheries. Nature Climate Change, 1(9), 449–456. https://doi.org/10.1038/nclimate1301

Valdimarsson, H., Astthorsson, O. S., & Palsson, J. (2012). Hydrographic variability in Icelandic waters during recent decades and related changes in distribution of some fish species. ICES Journal of Marine Science, 69(5), 816–825.

Valtýsson, H. Þ., & Jónsson, S. (2018). Impacts of a Changing Climate on Icelandic Marine Stocks. In F. J. Mueter, M. R. Baker, S. C. Dressel, & A. B. Hollowed (Eds.), Impacts of a Changing Environment on the Dynamics of High-latitude Fish and Fisheries. Alaska Sea Grant, University of Alaska, Fairbanks.

Wood, S. N. (2006). Generalized additive models: An introduction with R,.(Chapman and Hall/CRC: Florida).

Wood, S. N. (2011). Fast stable restricted maximum likelihood and marginal likelihood estimation of semiparametric generalized linear models. Journal of the Royal Statistical Society (B), 73(1), 3–36.

