## Supplementary figures and images for "Projecting climate-driven shifts in demersal fish habitat in Iceland’s waters"

### Supplemental Figure 1

#### Mean projected annual temperature in Iceland's EEZ

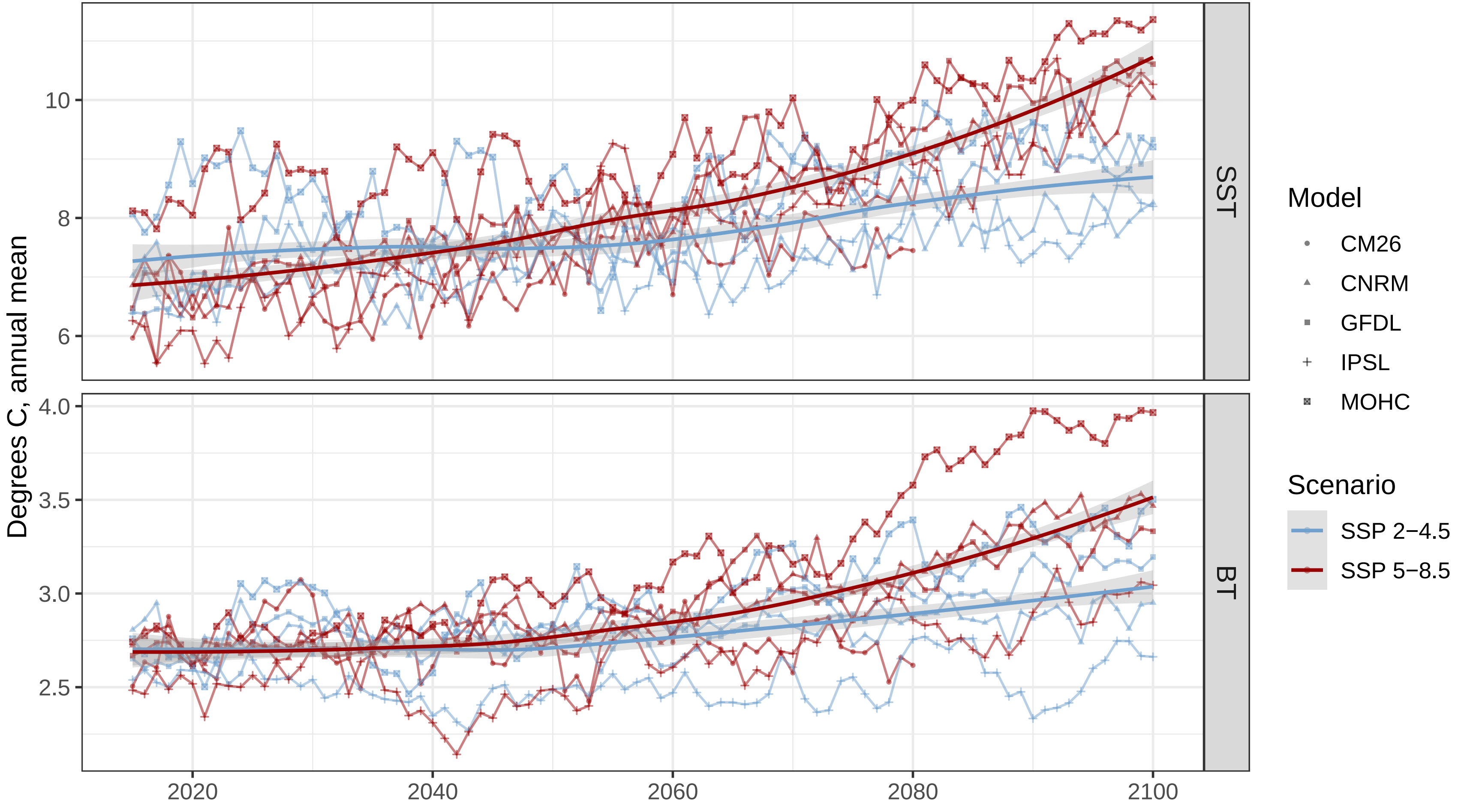

### Supplemental Figure 3

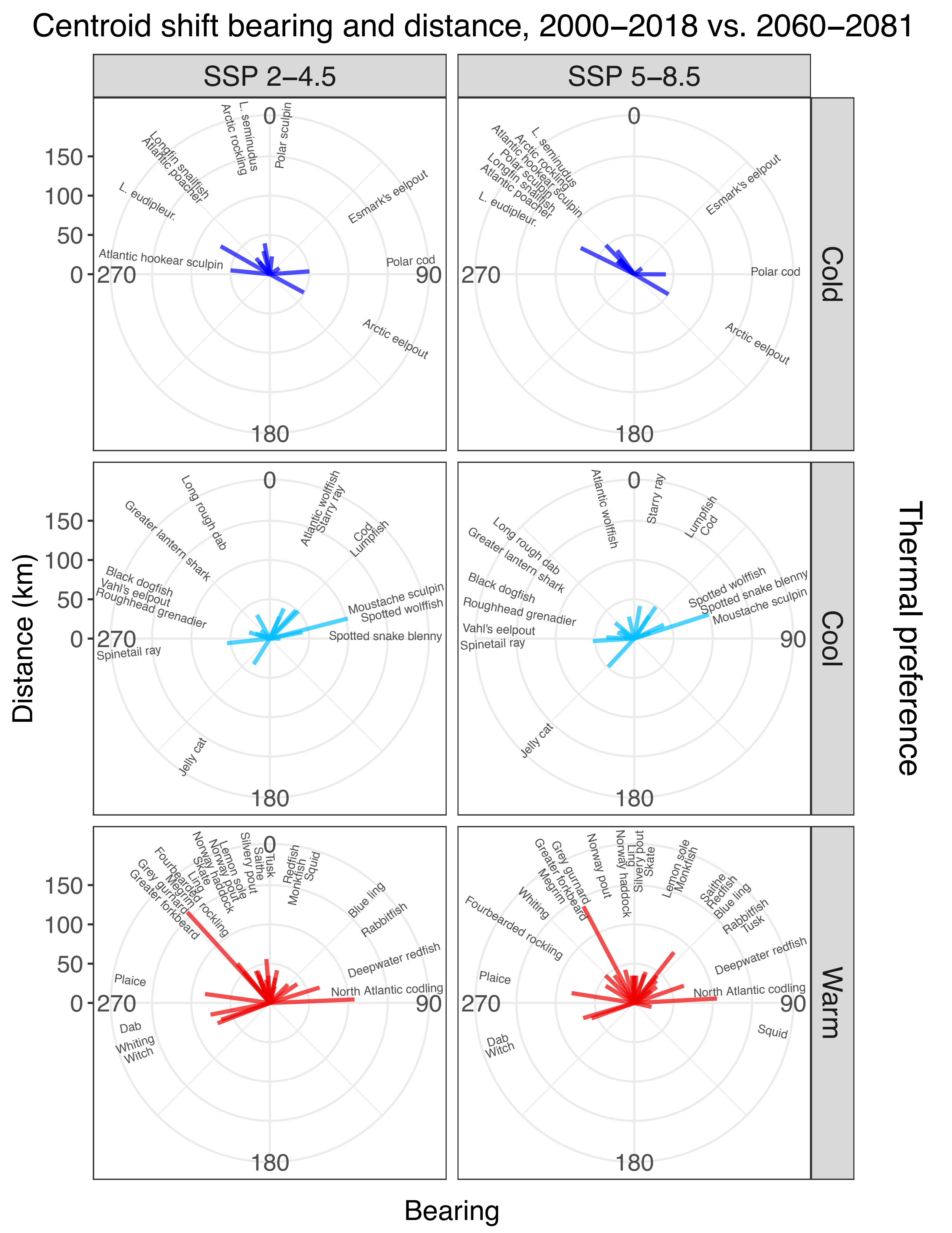
