## Supplemental Figure 2 for "Projecting climate-driven shifts in demersal fish habitat in Iceland’s waters"

### Starry ray, thorny skate, *Amblyraja radiata*

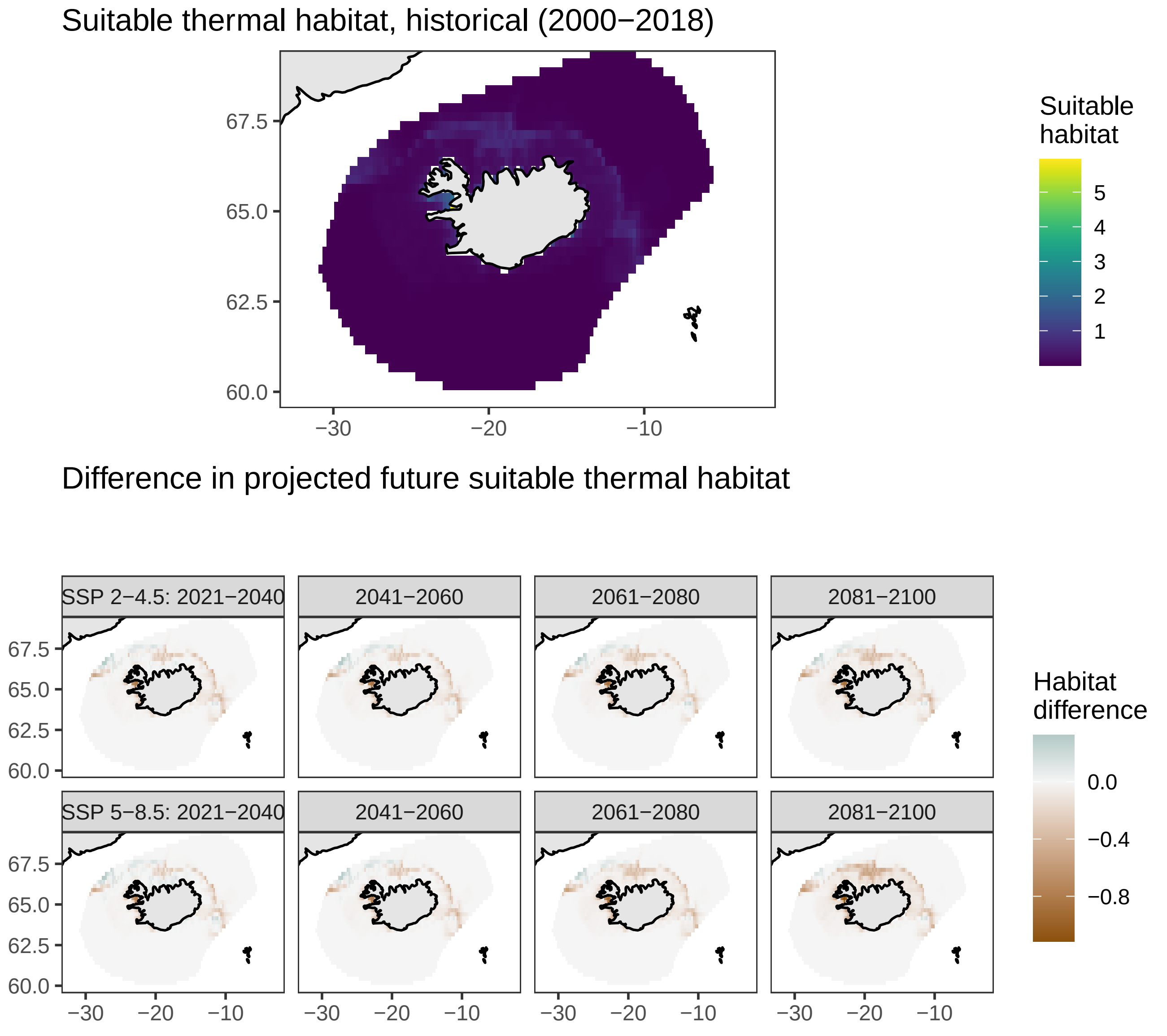

### Jelly cat, Arctic wolffish, *Anarhichas denticulatus*

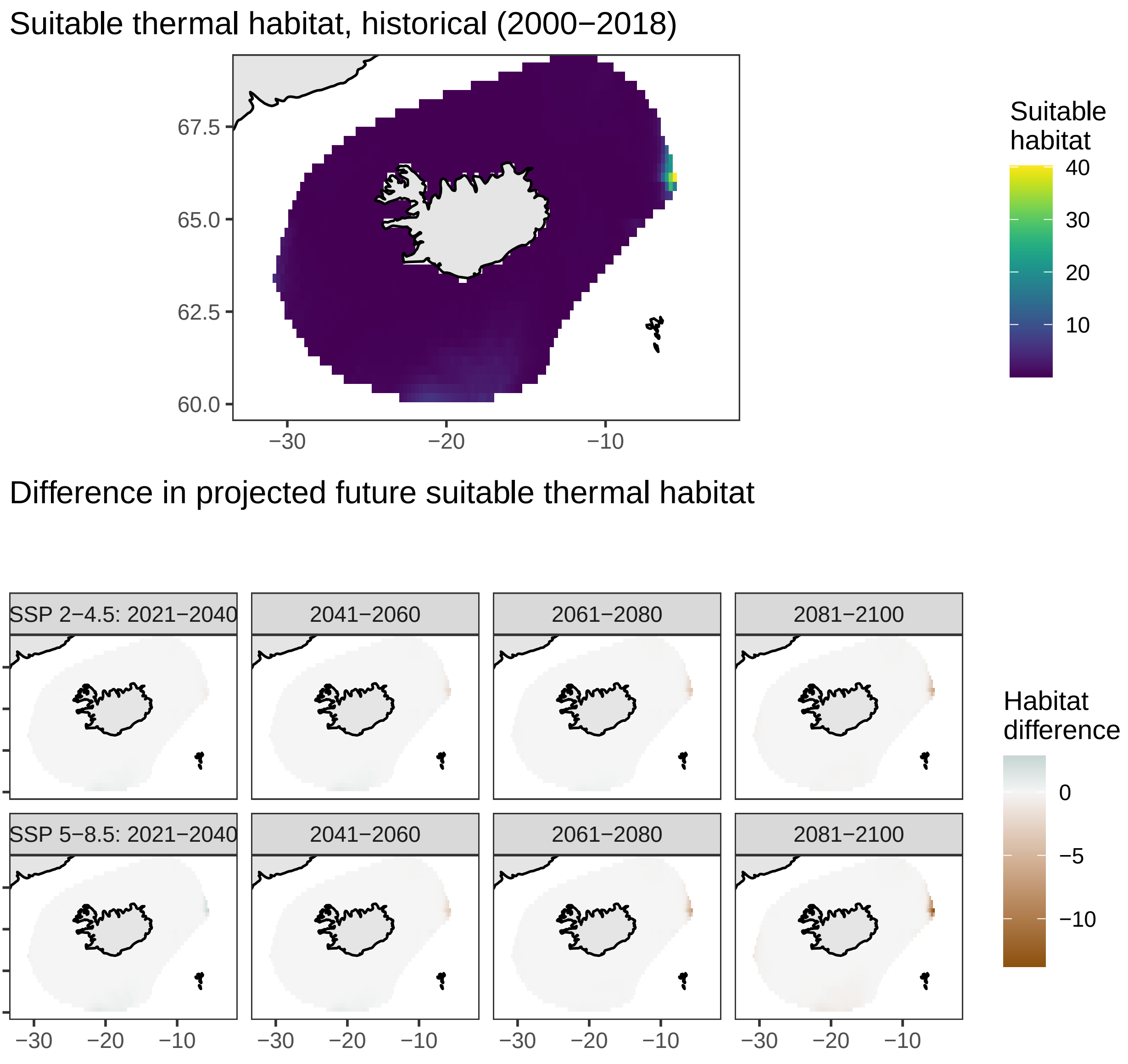

### Atlantic wolffish, *Anarhichas lupus*

Suitable thermal habitat, historical (2000−2018)

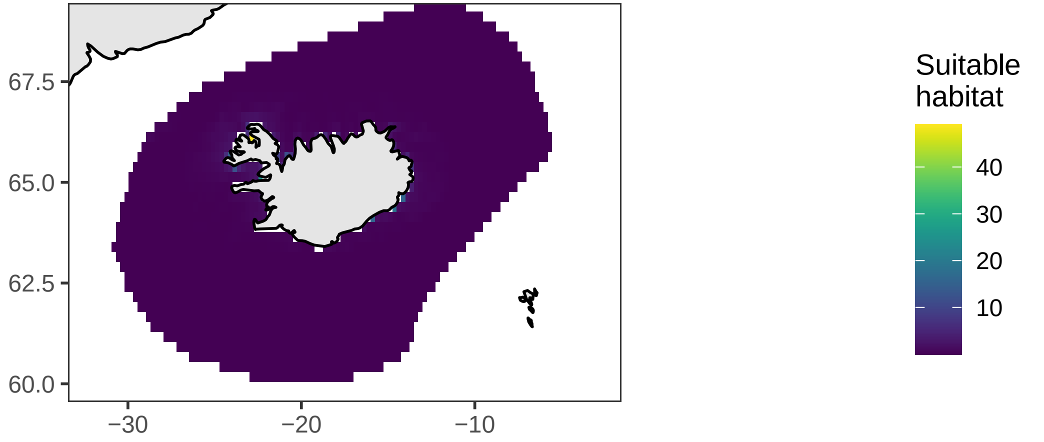


Difference in projected future suitable thermal habitat

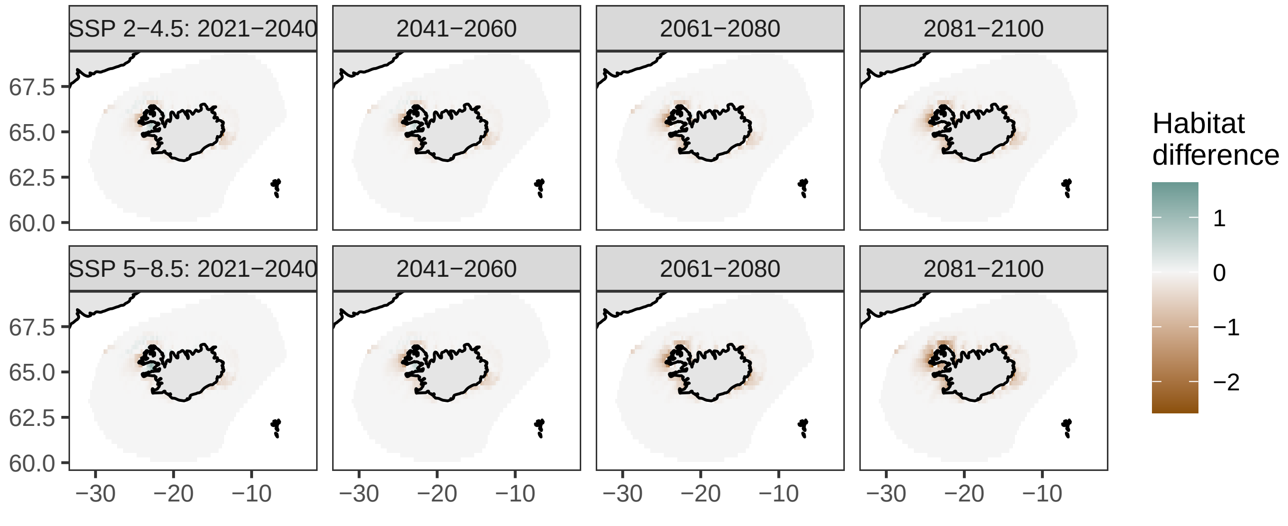

### Spotted wolffish, leopardfish, *Anarhichas minor*

Suitable thermal habitat, historical (2000−2018)

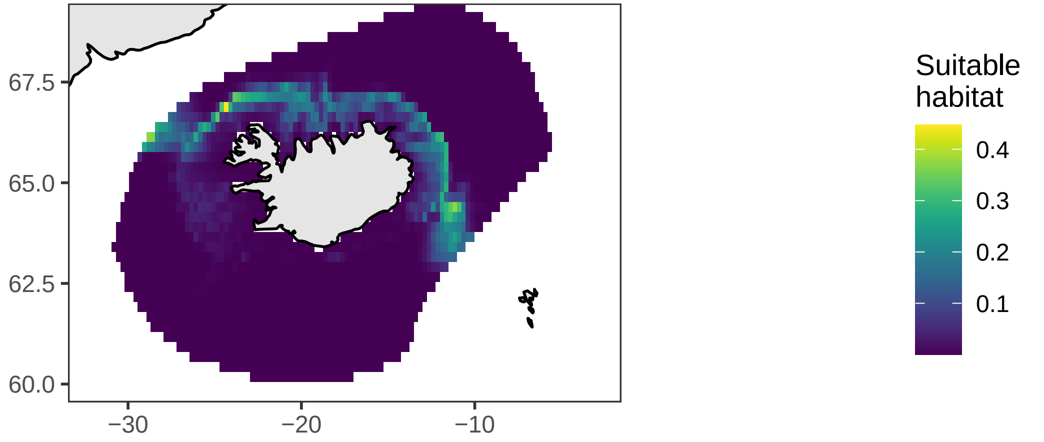


Difference in projected future suitable thermal habitat

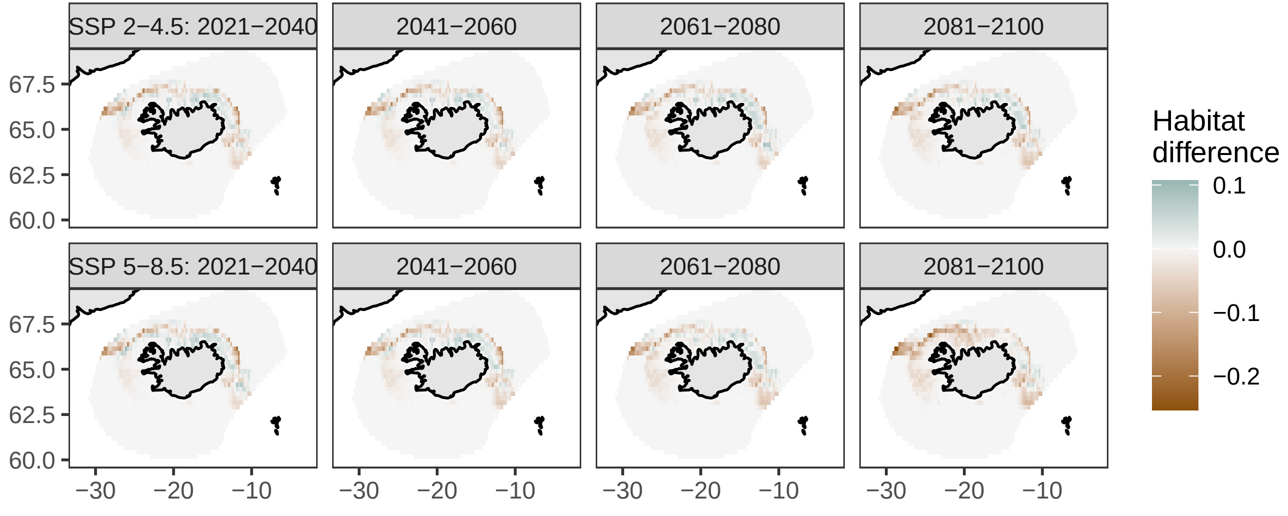

### Atlantic hookear sculpin, *Artediellus atlanticus*

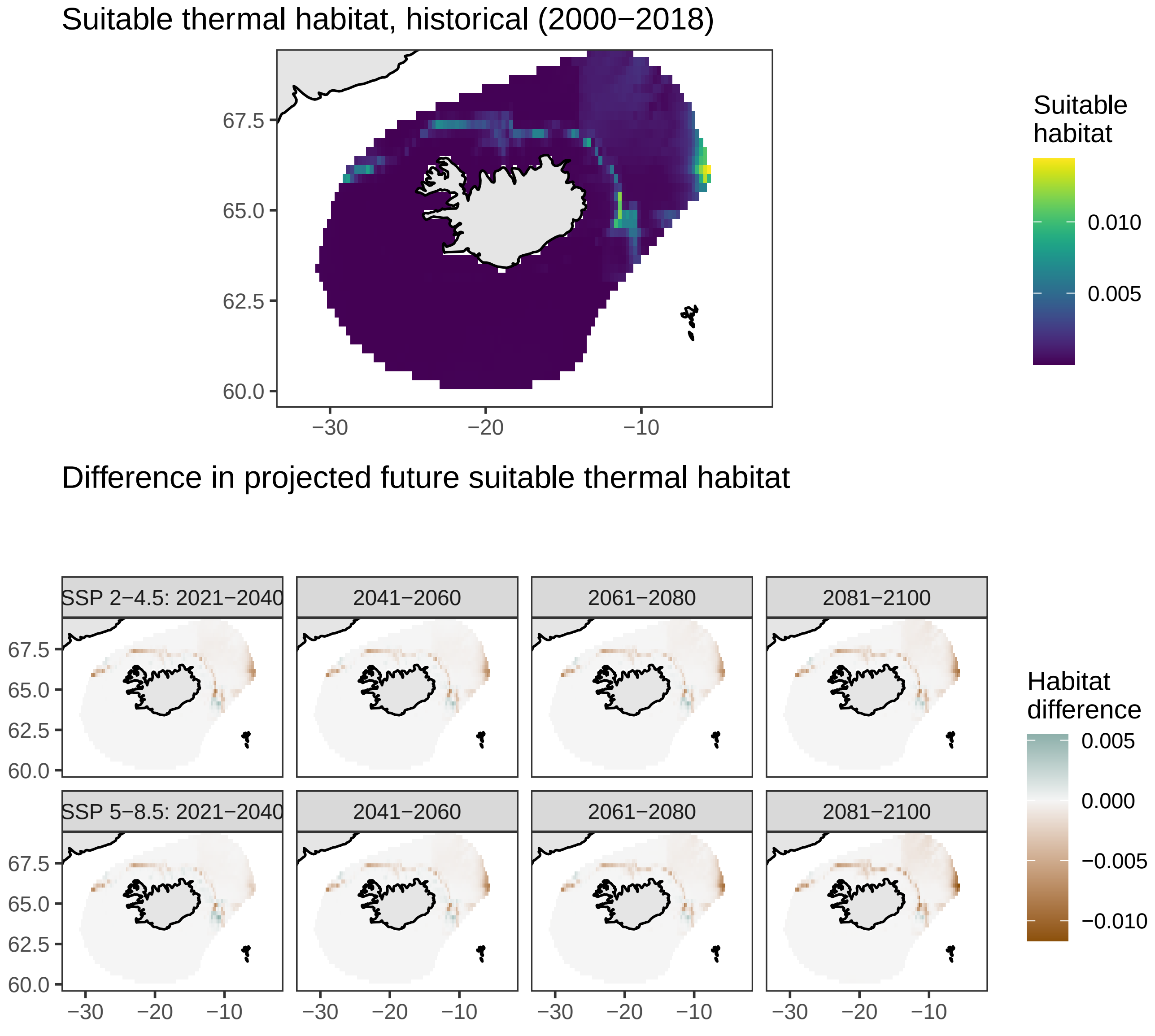

### Spinetail ray, spinetail skate, *Bathyraja spinicauda*

Suitable thermal habitat, historical (2000−2018)

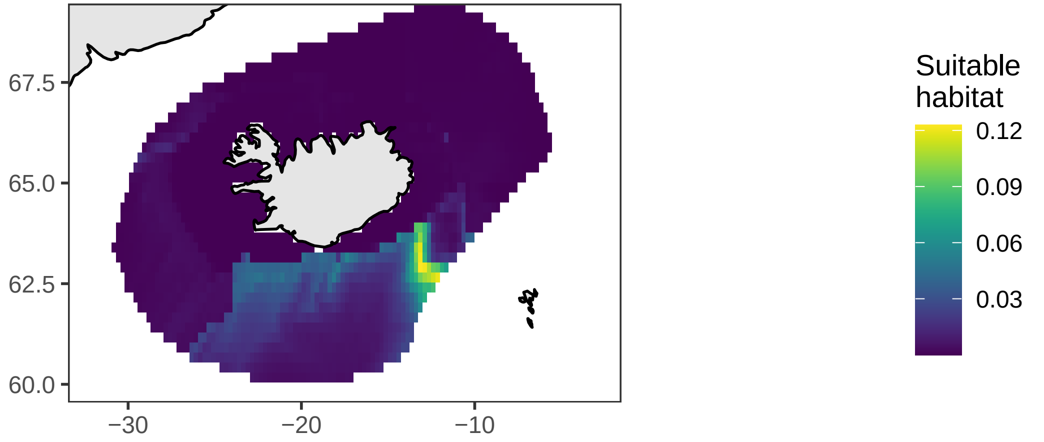


Difference in projected future suitable thermal habitat

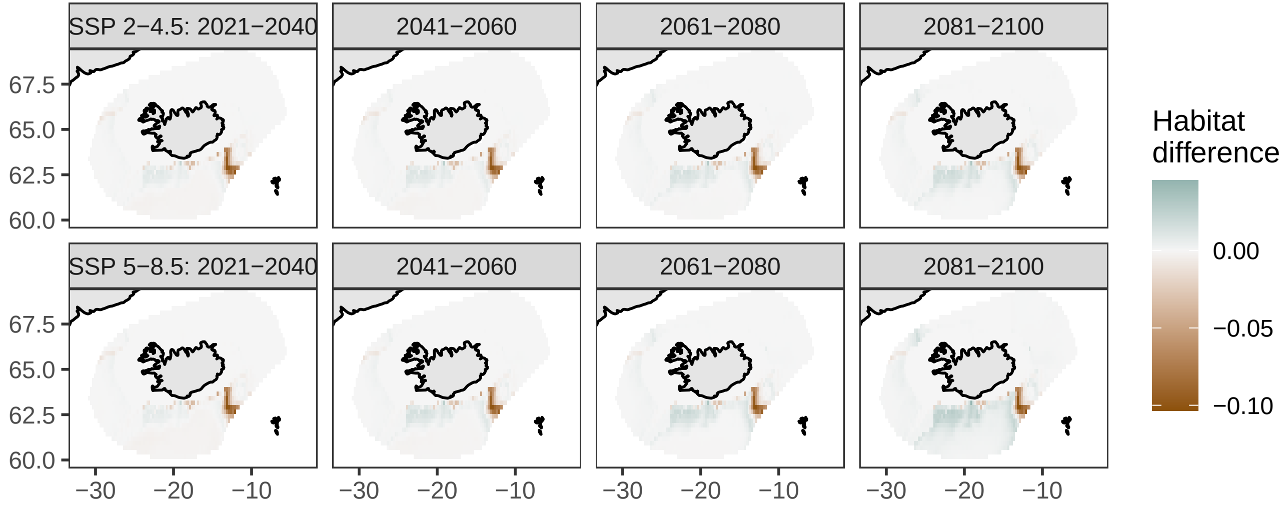

### Polar cod, *Boreogadus saida*

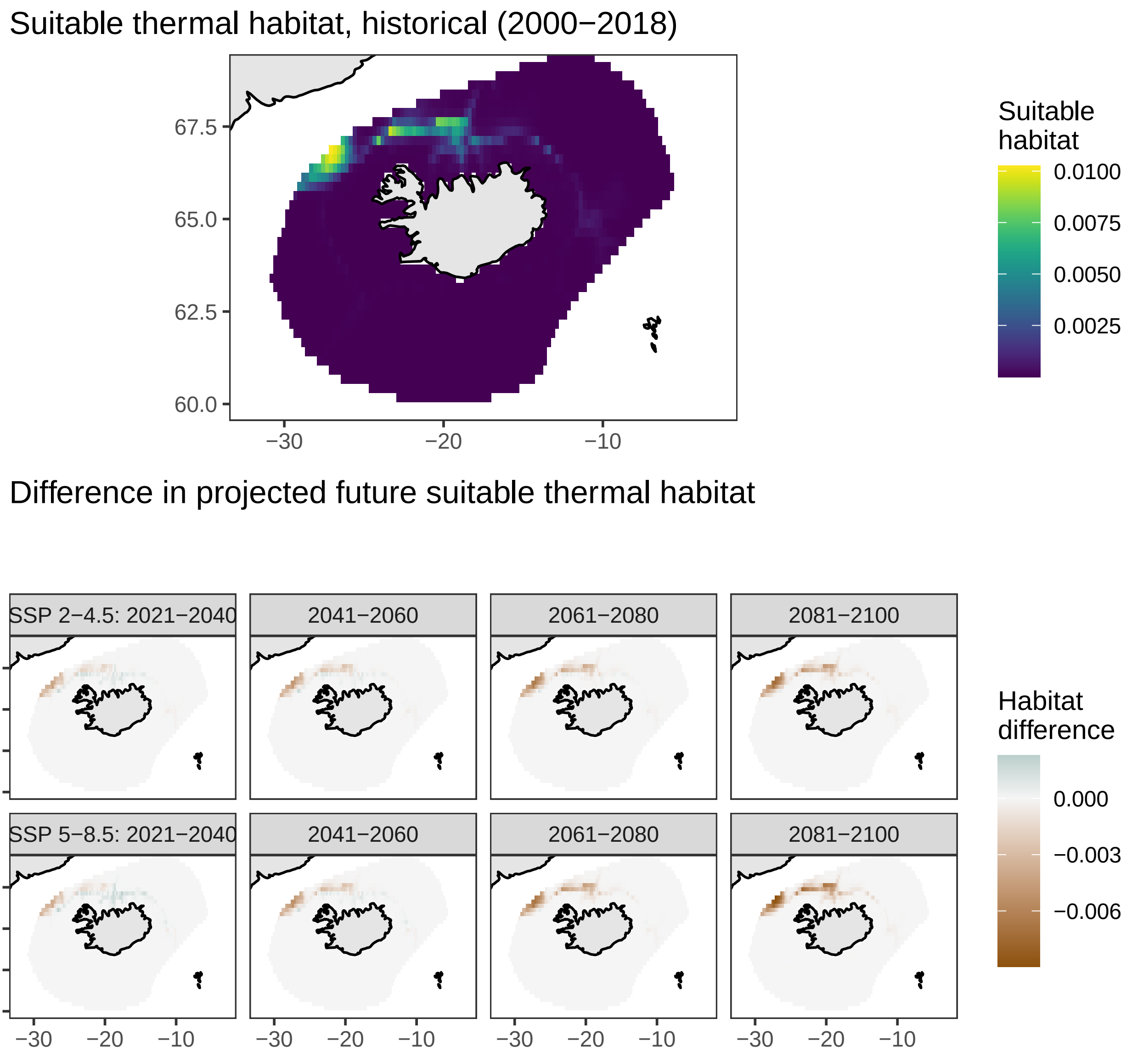

### Tusk, torsk, cusk, *Brosme brosme*

#### Suitable thermal habitat, historical (2000−2018)

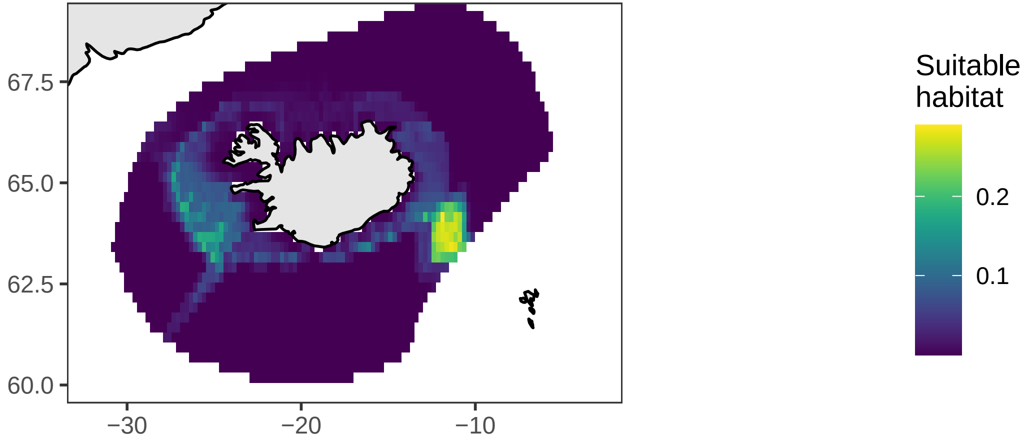


#### Difference in projected future suitable thermal habitat

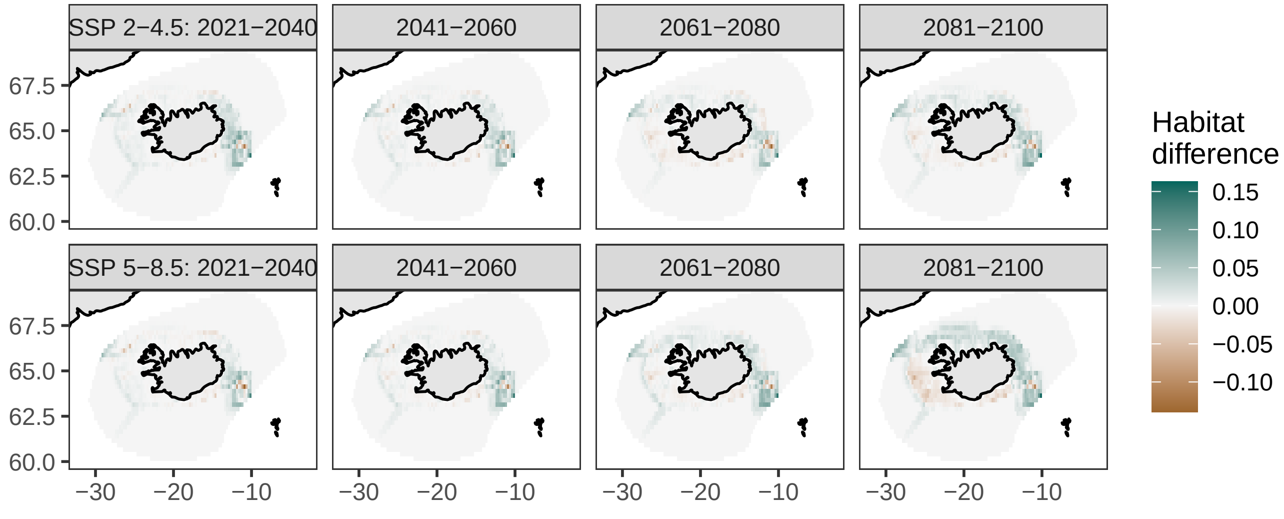

### Longfin snailfish, sea tadpole, *Careproctus reinhardti*

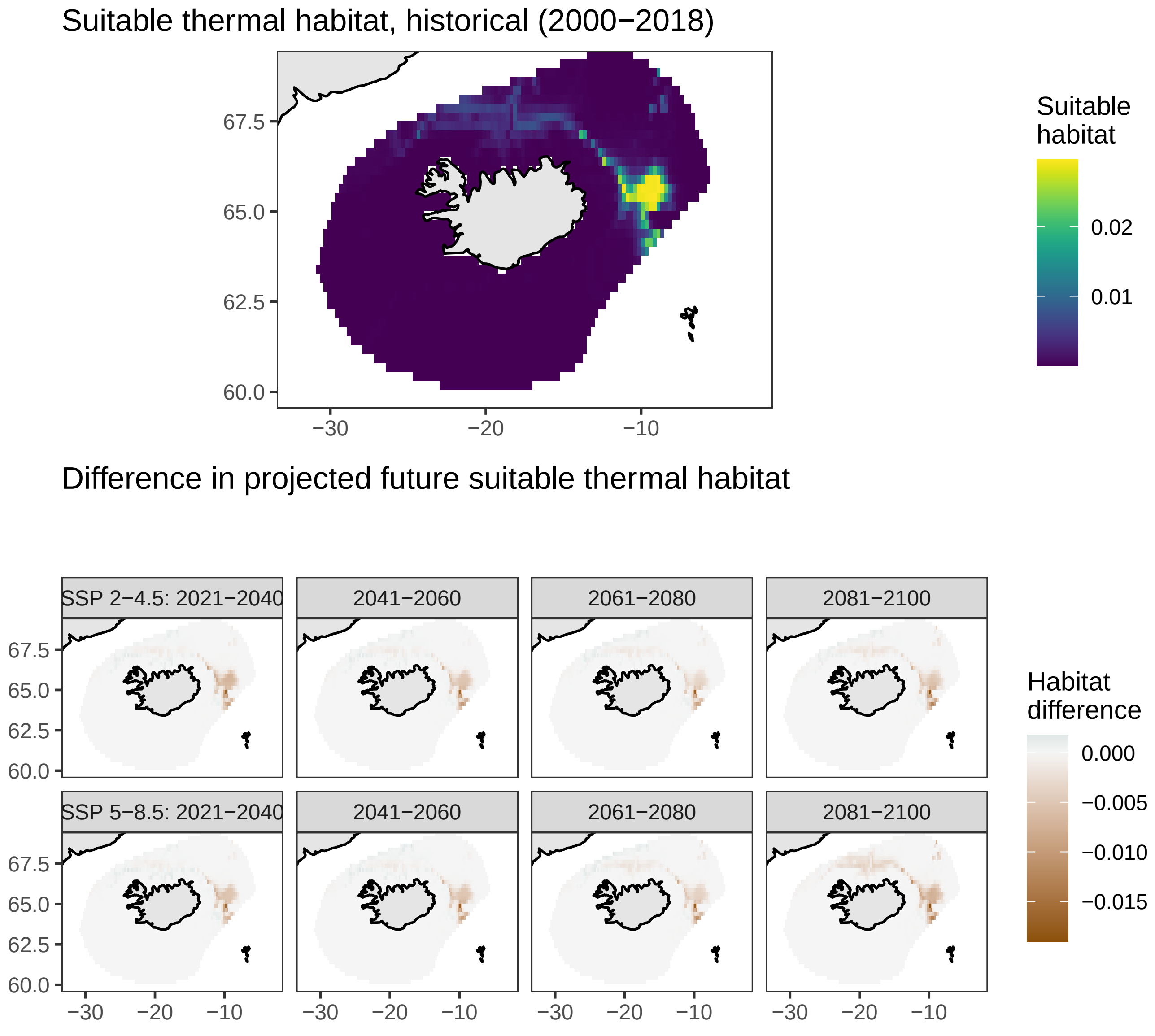

### Black dogfish, *Centroscyllium fabricii*

Suitable thermal habitat, historical (2000−2018)

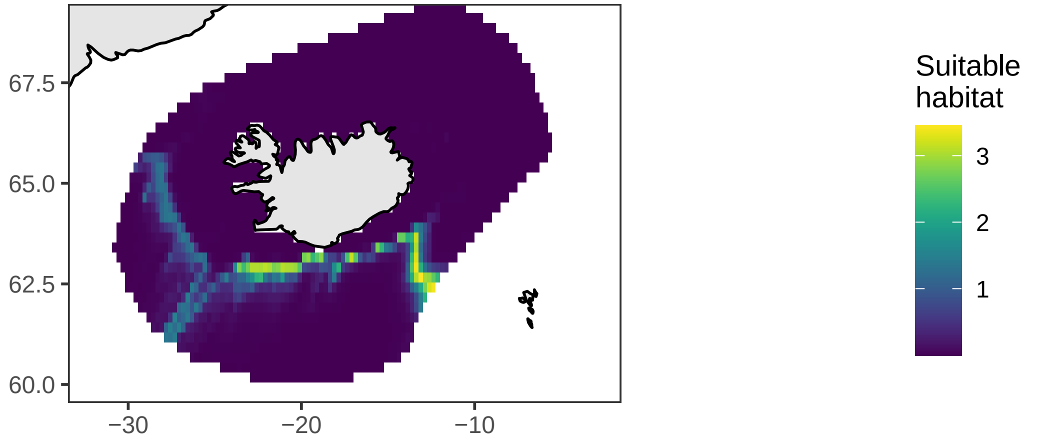


Difference in projected future suitable thermal habitat

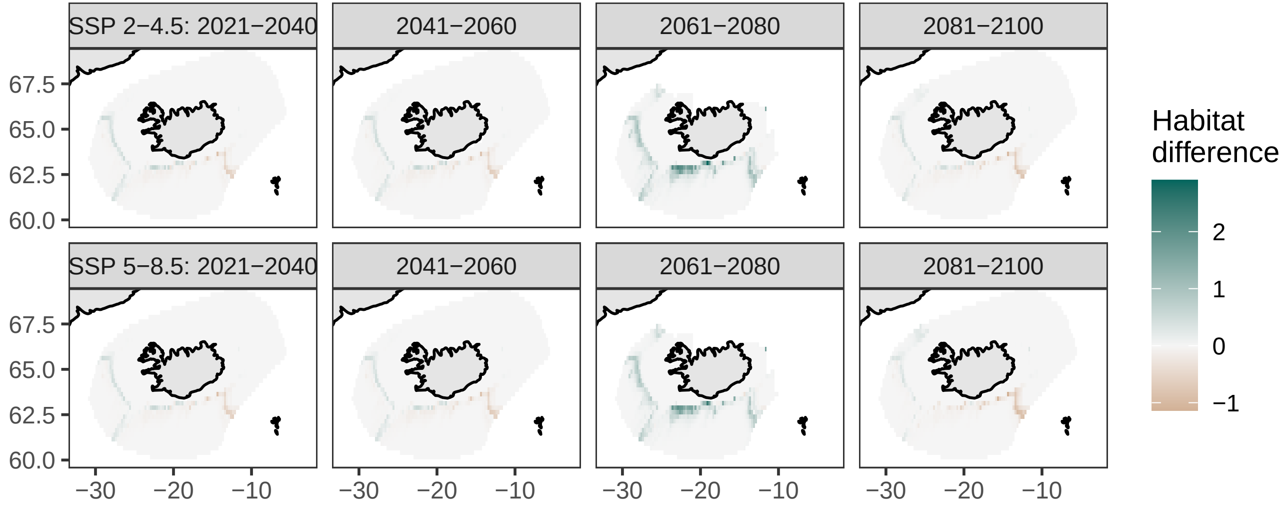

### Rabbitfish (rat fish), *Chimaera monstrosa*

#### Suitable thermal habitat, historical (2000−2018)

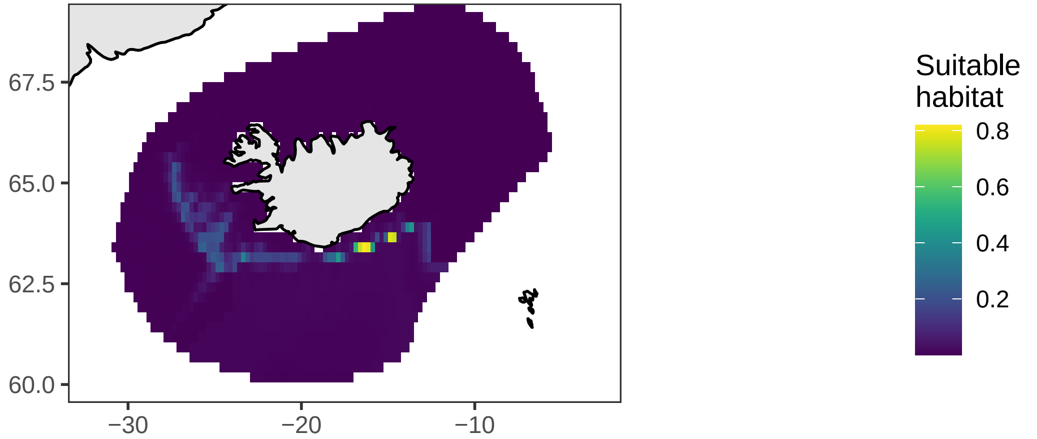


#### Difference in projected future suitable thermal habitat

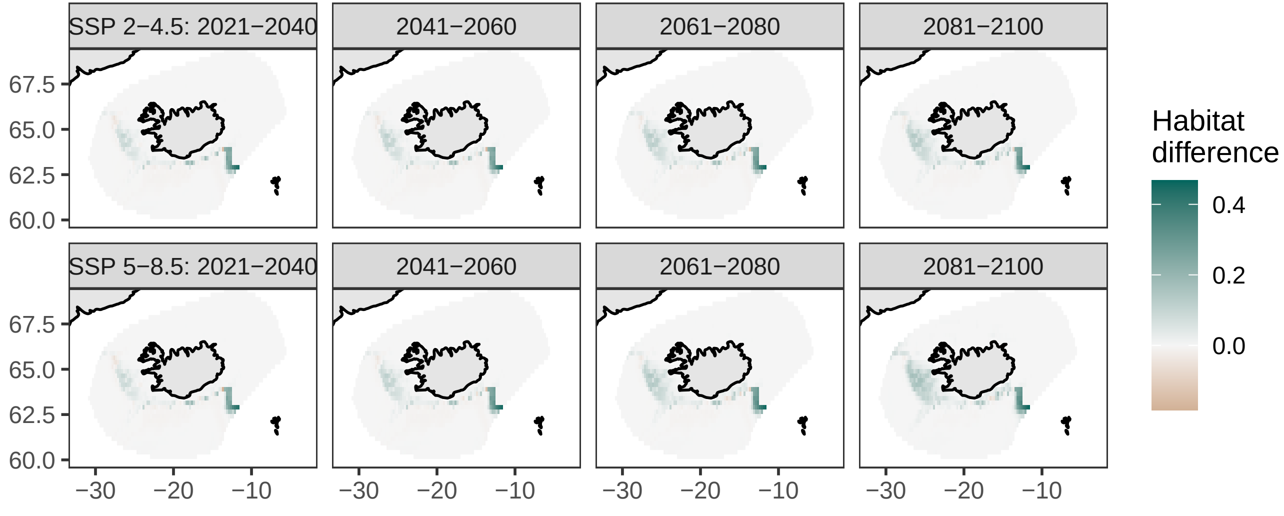

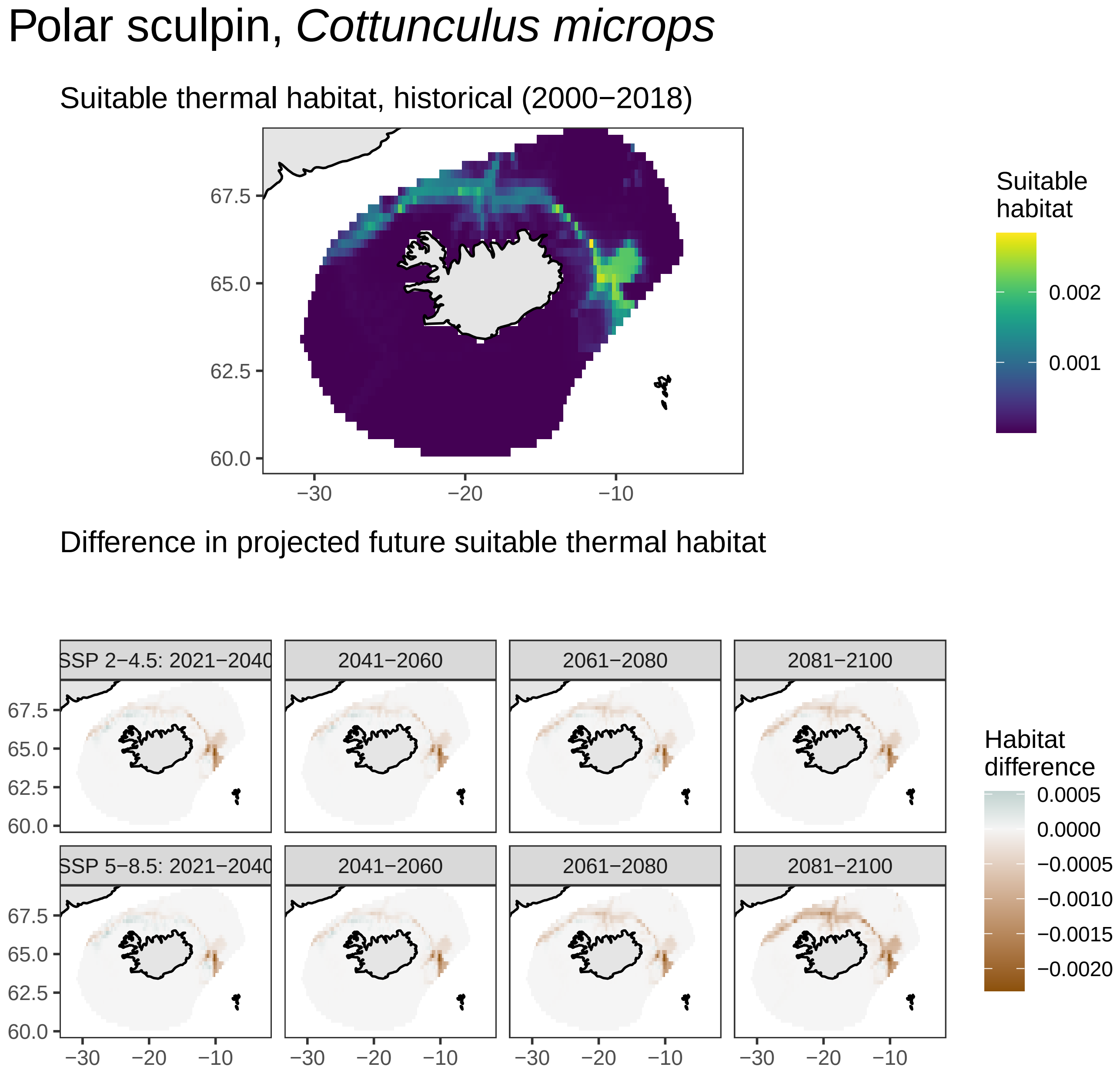

Polar sculpin, *Cottunculus microps*
Suitable thermal habitat, historical (2000−2018)
67.5
65.0
62.5
60.0
−30
−20
−10
Suitable habitat
0.002
0.001
Difference in projected future suitable thermal habitat
SSP 2−4.5: 2021−2040
2041−2060
2061−2080
2081−2100
SSP 5−8.5: 2021−2040
2041−2060
2061−2080
2081−2100
Habitat difference
0.0005
0.0000
−0.0005
−0.0010
−0.0015
−0.0020

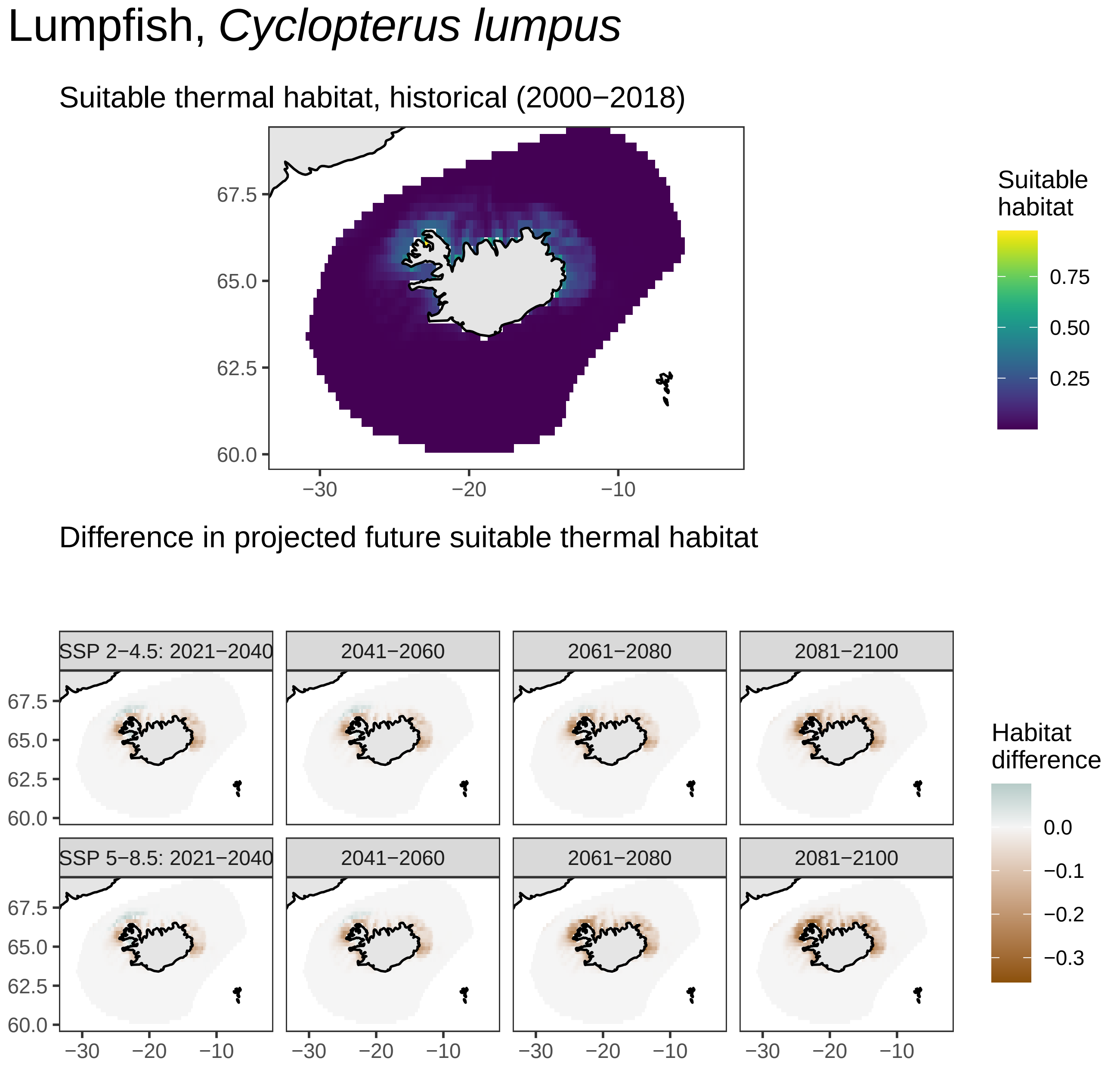

Lumpfish, *Cyclopterus lumpus*
Suitable thermal habitat, historical (2000−2018)
67.5
65.0
62.5
60.0
−30
−20
−10
Suitable habitat
0.75
0.50
0.25
Difference in projected future suitable thermal habitat
SSP 2−4.5: 2021−2040
2041−2060
2061−2080
2081−2100
SSP 5−8.5: 2021−2040
2041−2060
2061−2080
2081−2100
Habitat difference
0.0
−0.1
−0.2
−0.3

### Skate, *Dipturus batis*

Suitable thermal habitat, historical (2000−2018)

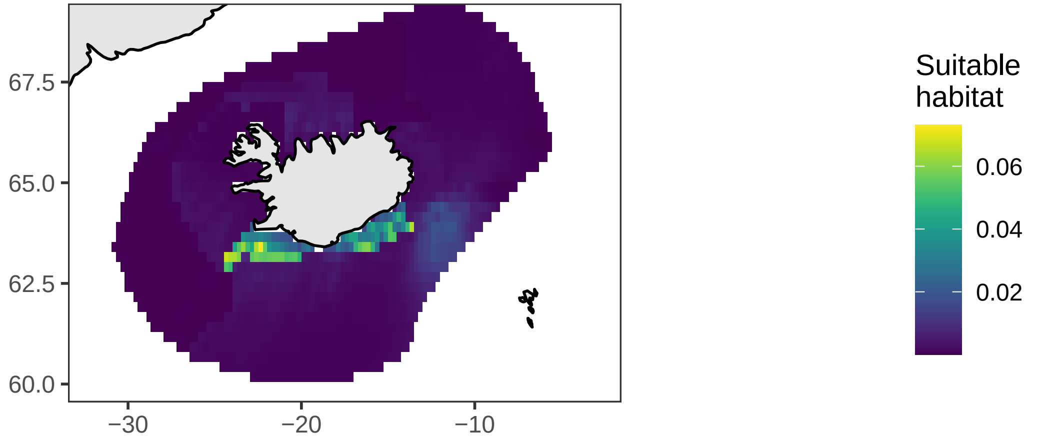


Difference in projected future suitable thermal habitat

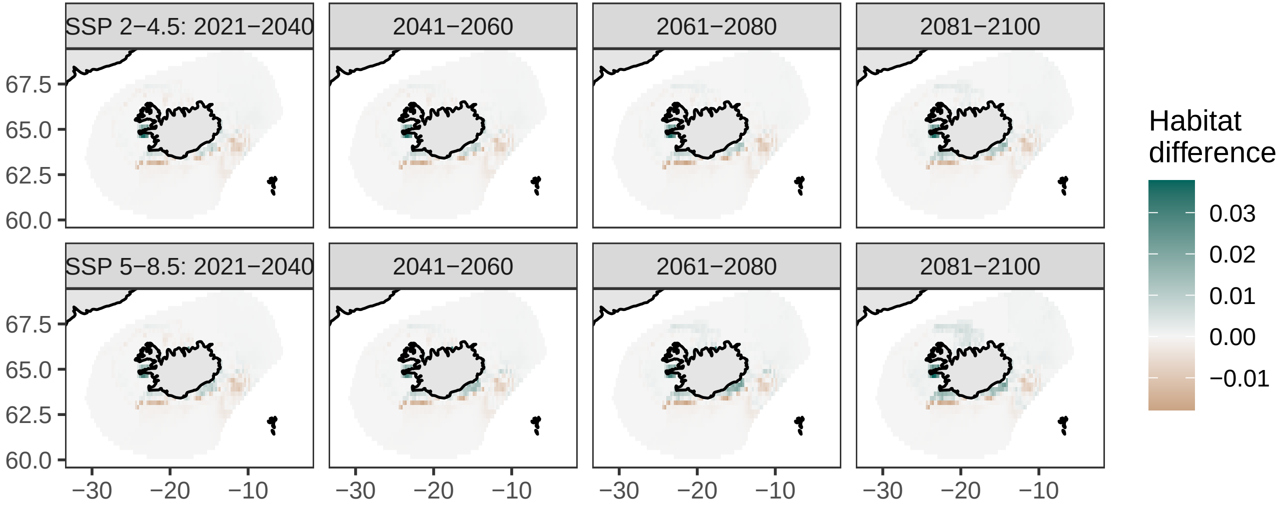

### Greater lantern shark, *Etmopterus princeps*

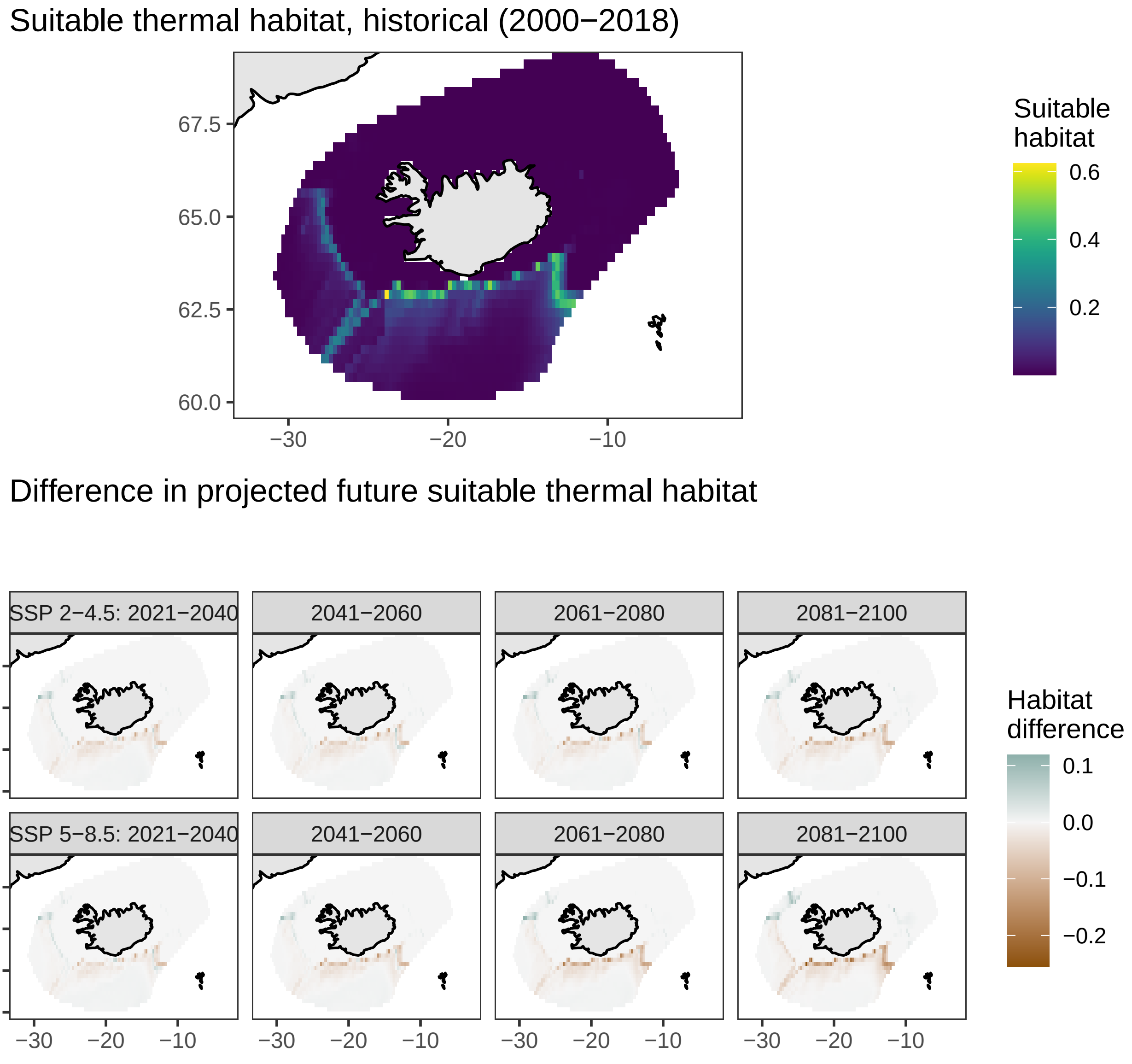

### Grey gurnard, *Eutrigla gurnardus*

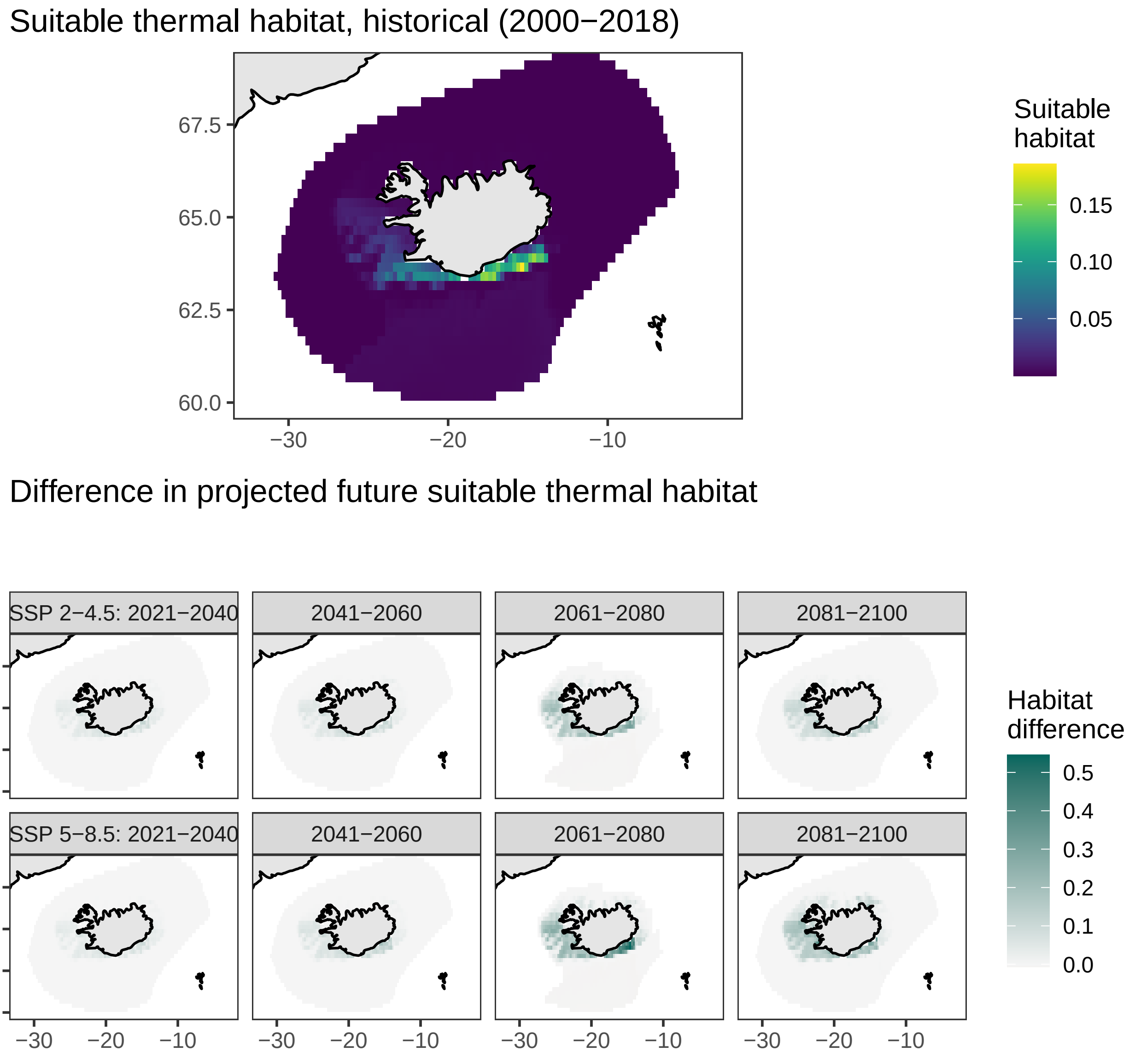

### Silvery pout, *Gadiculus argenteus*

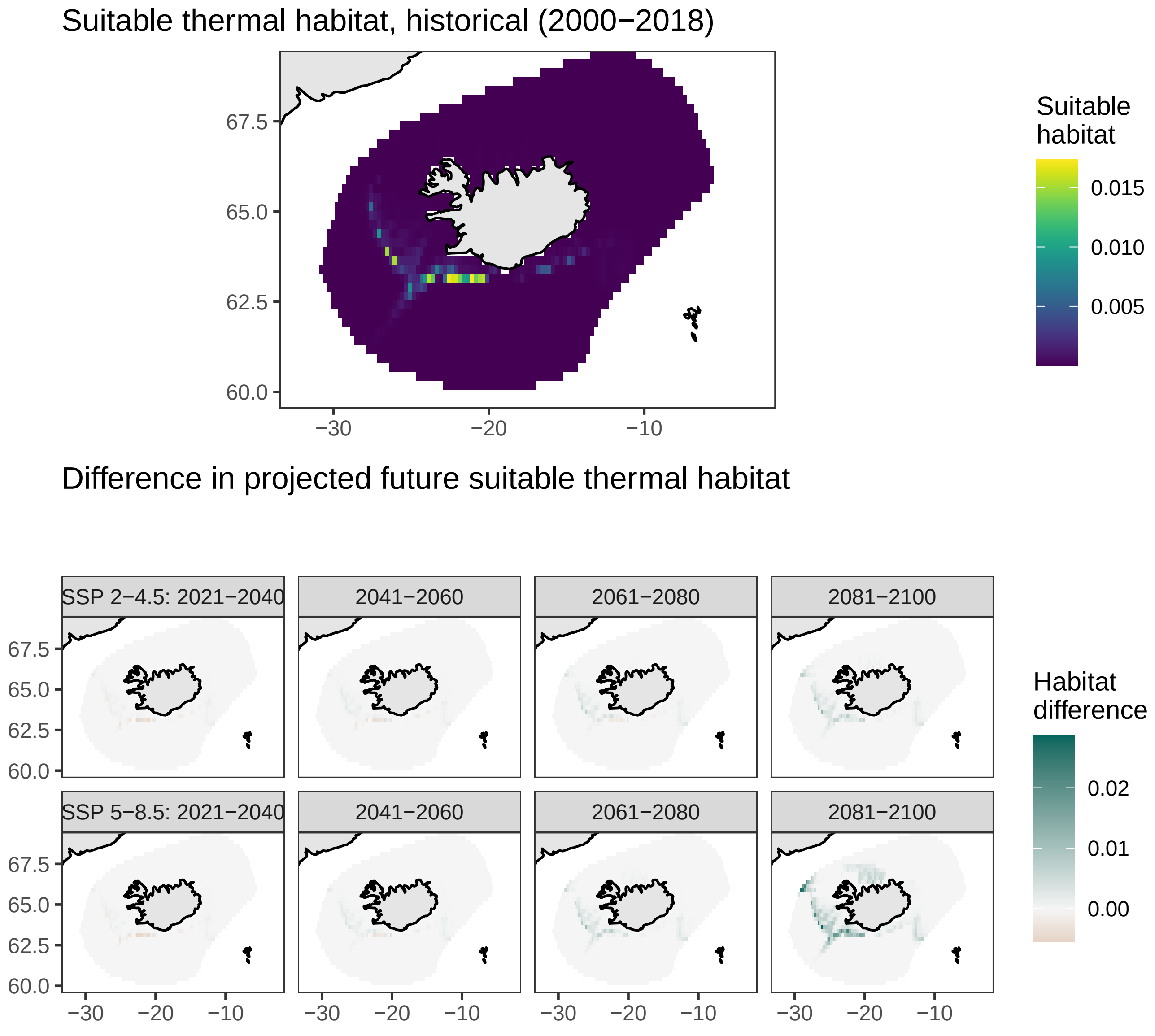

### Cod, *Gadus morhua*

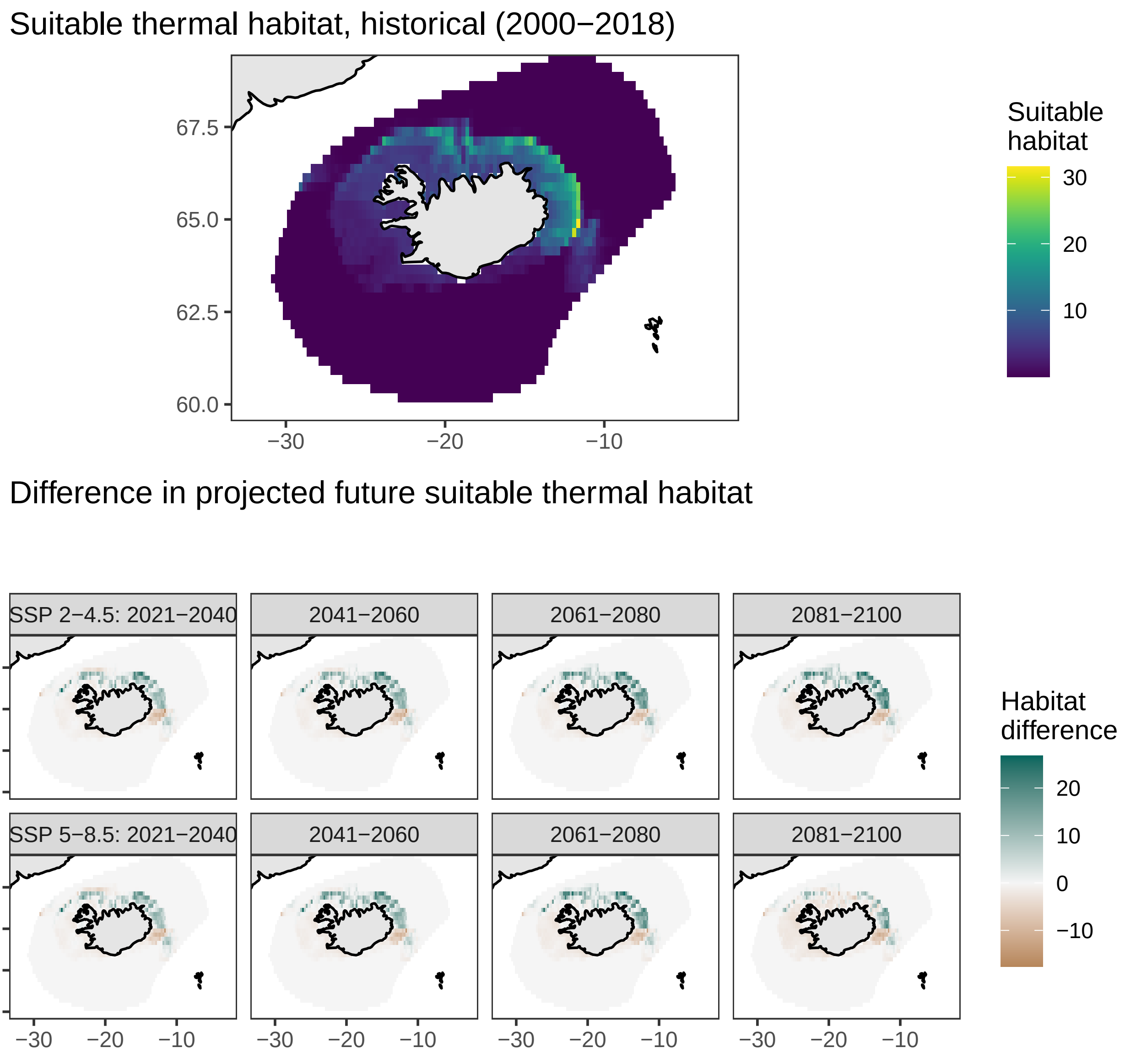

### Arctic rockling, *Gaidropsarus argentatus*

Suitable thermal habitat, historical (2000−2018)

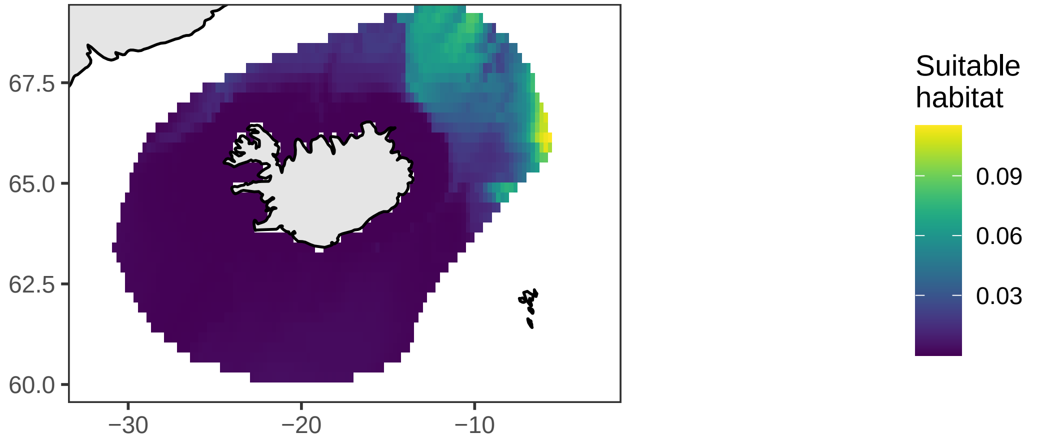


Difference in projected future suitable thermal habitat

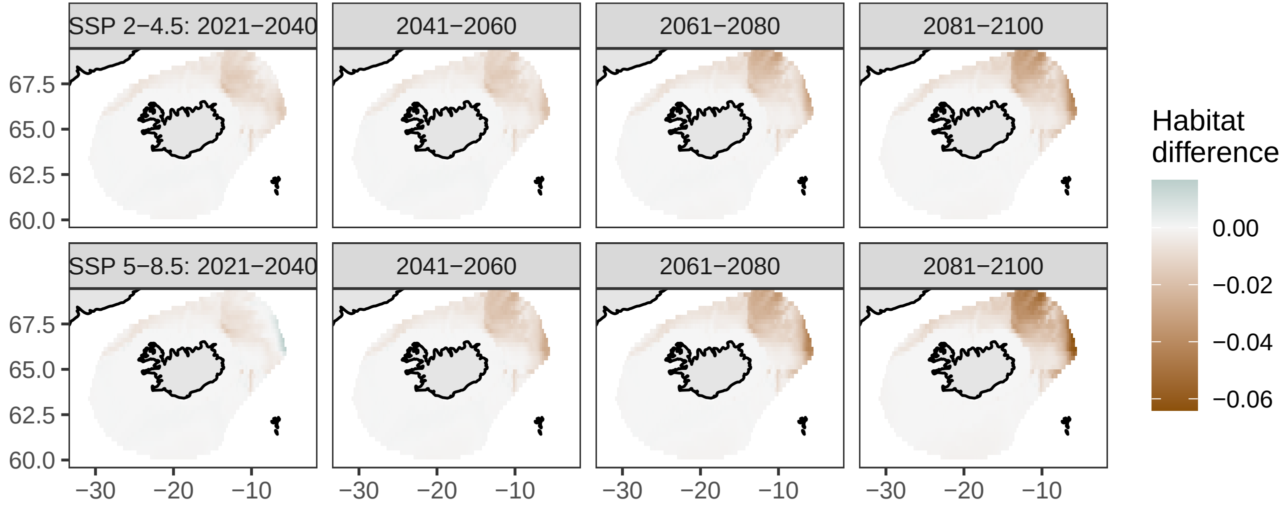

### Witch, *Glyptocephalus cynoglossus*

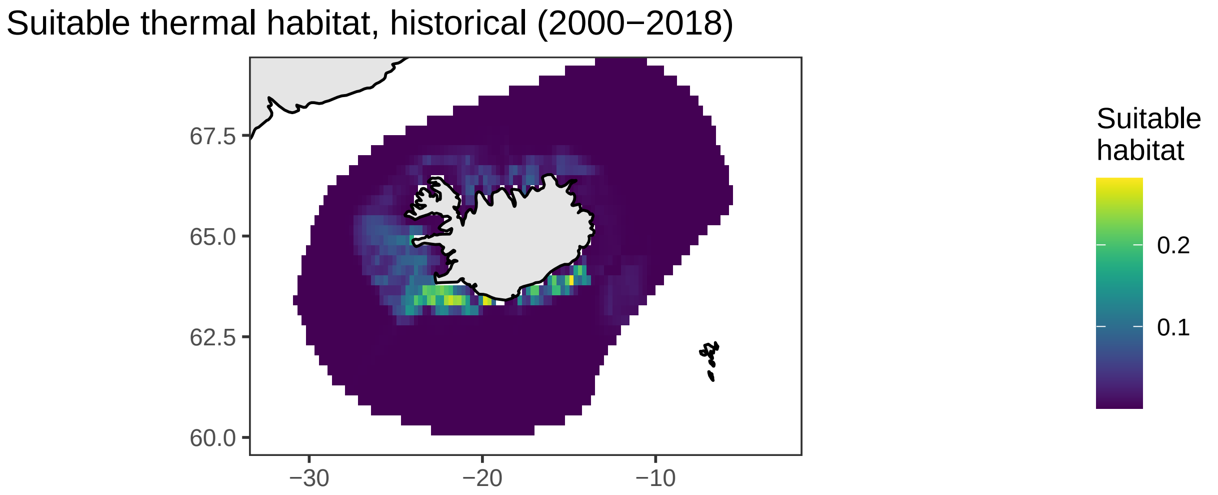


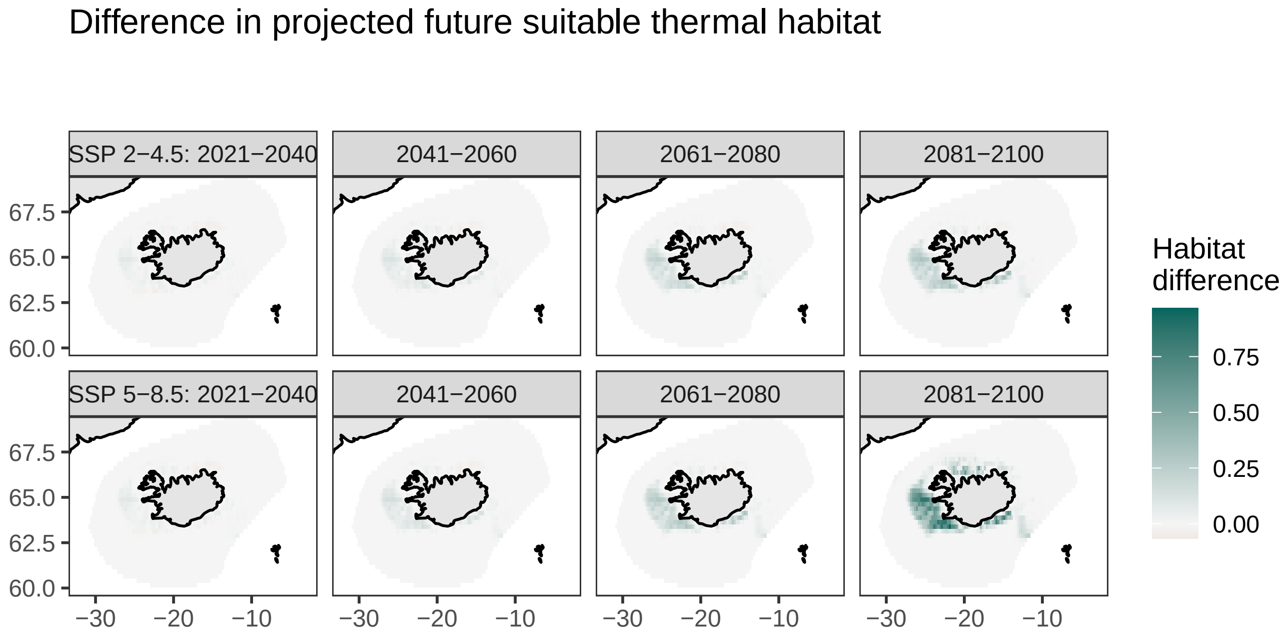

### Long rough dab, *Hippoglossoides platessoides*

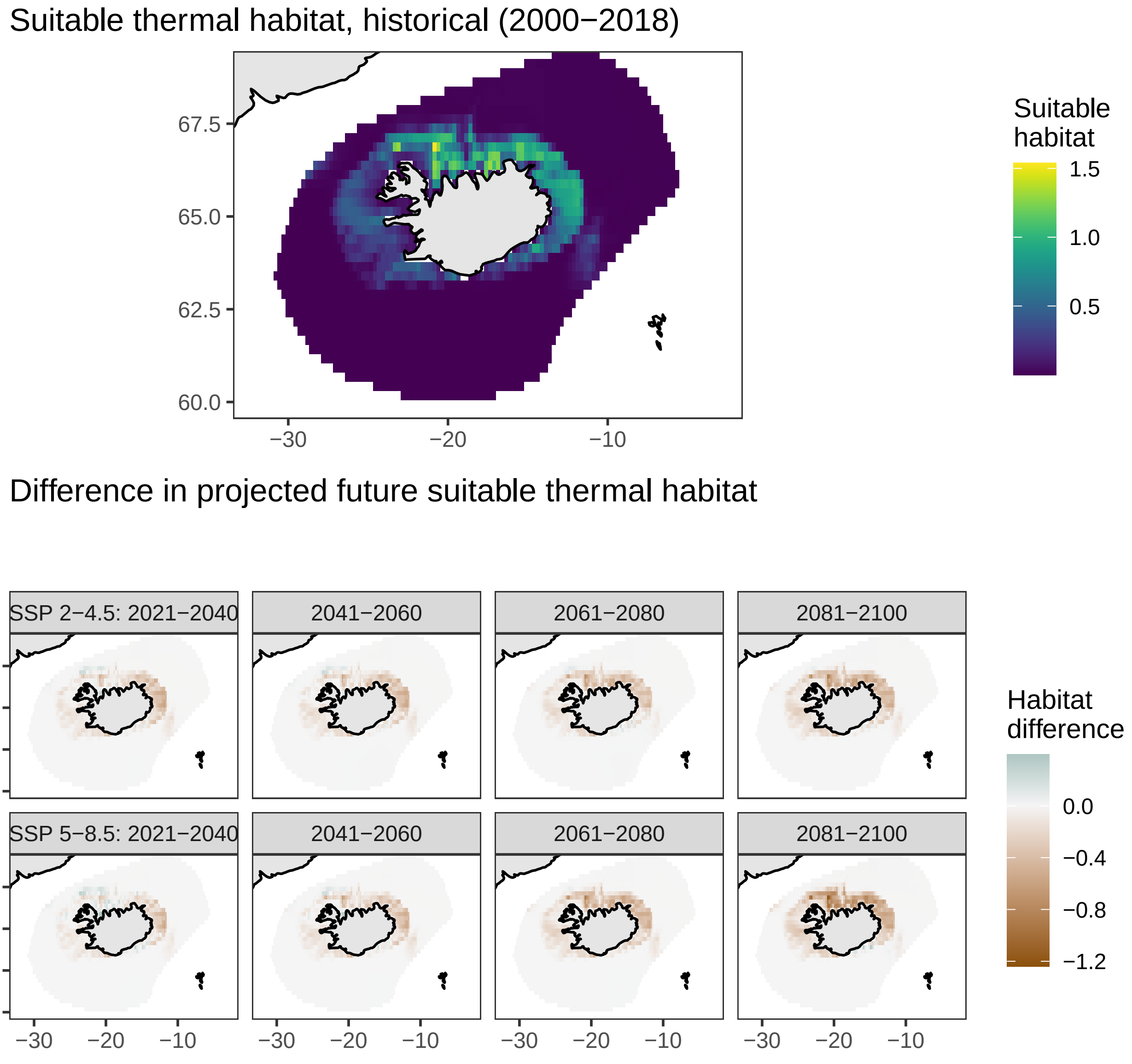

### North Atlantic codling, *Lepidion eques*

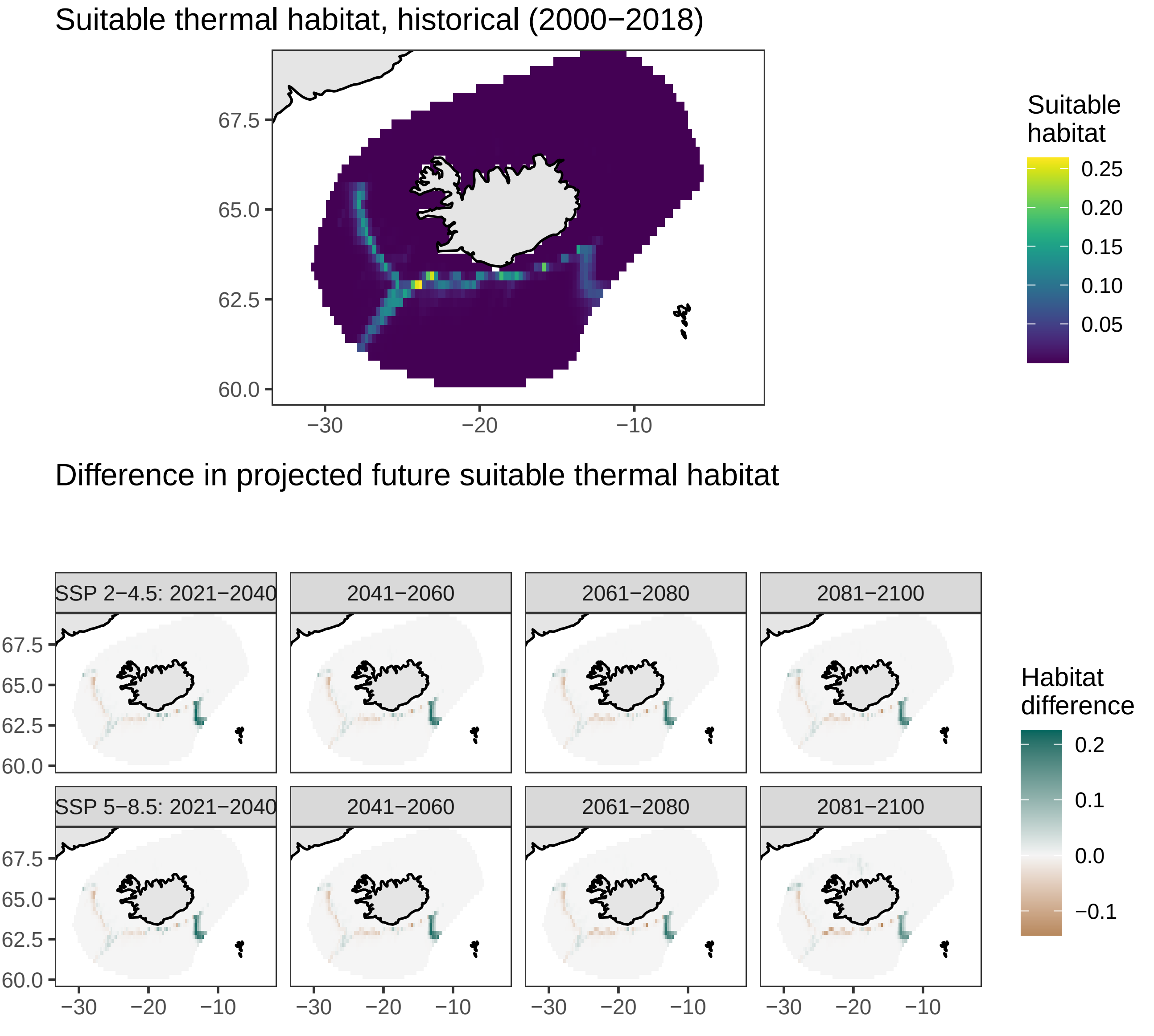

### Megrim, *Lepidorhombus whiffiagonis*

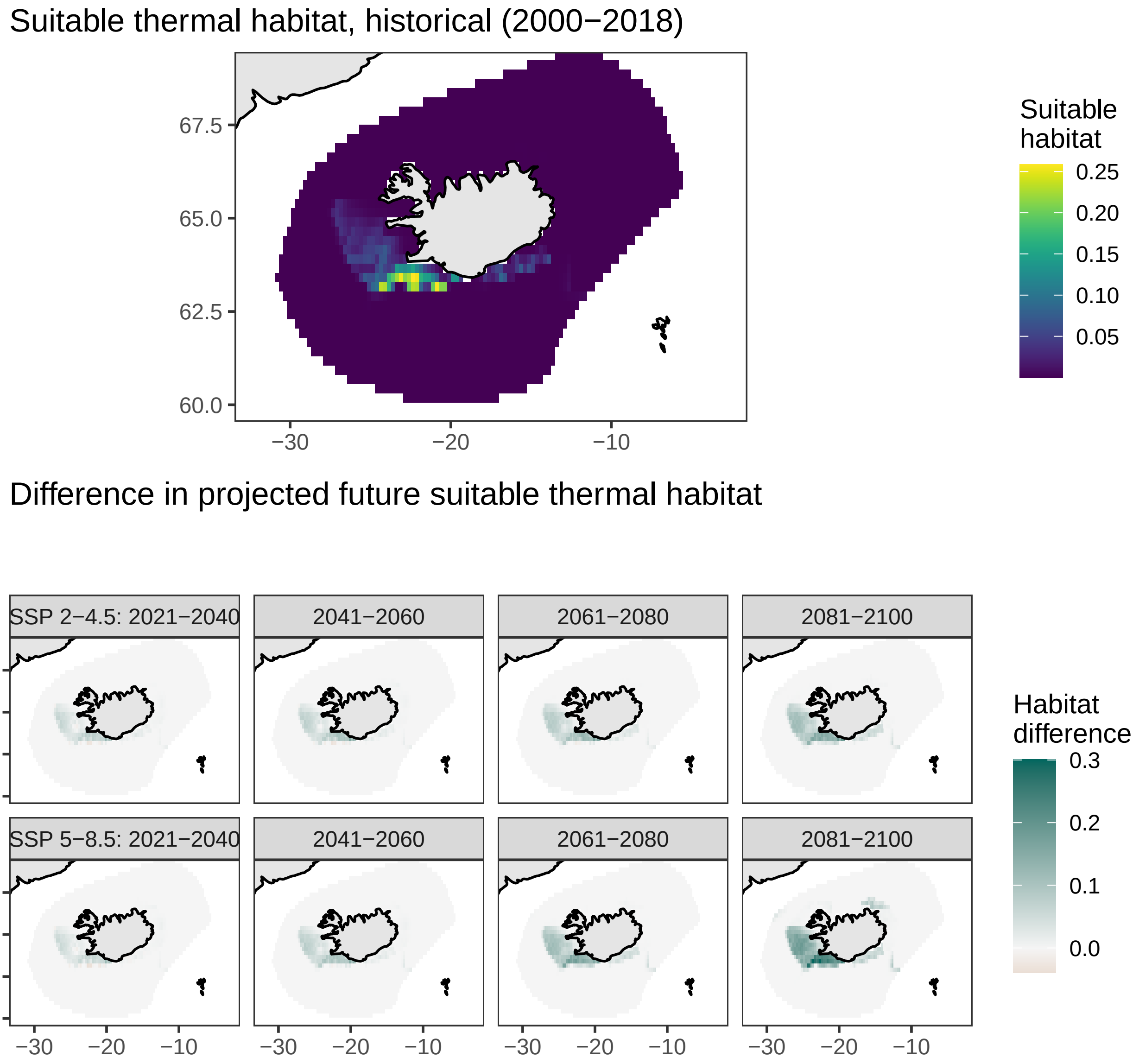

### Atlantic poacher, *Leptagonus decagonus*

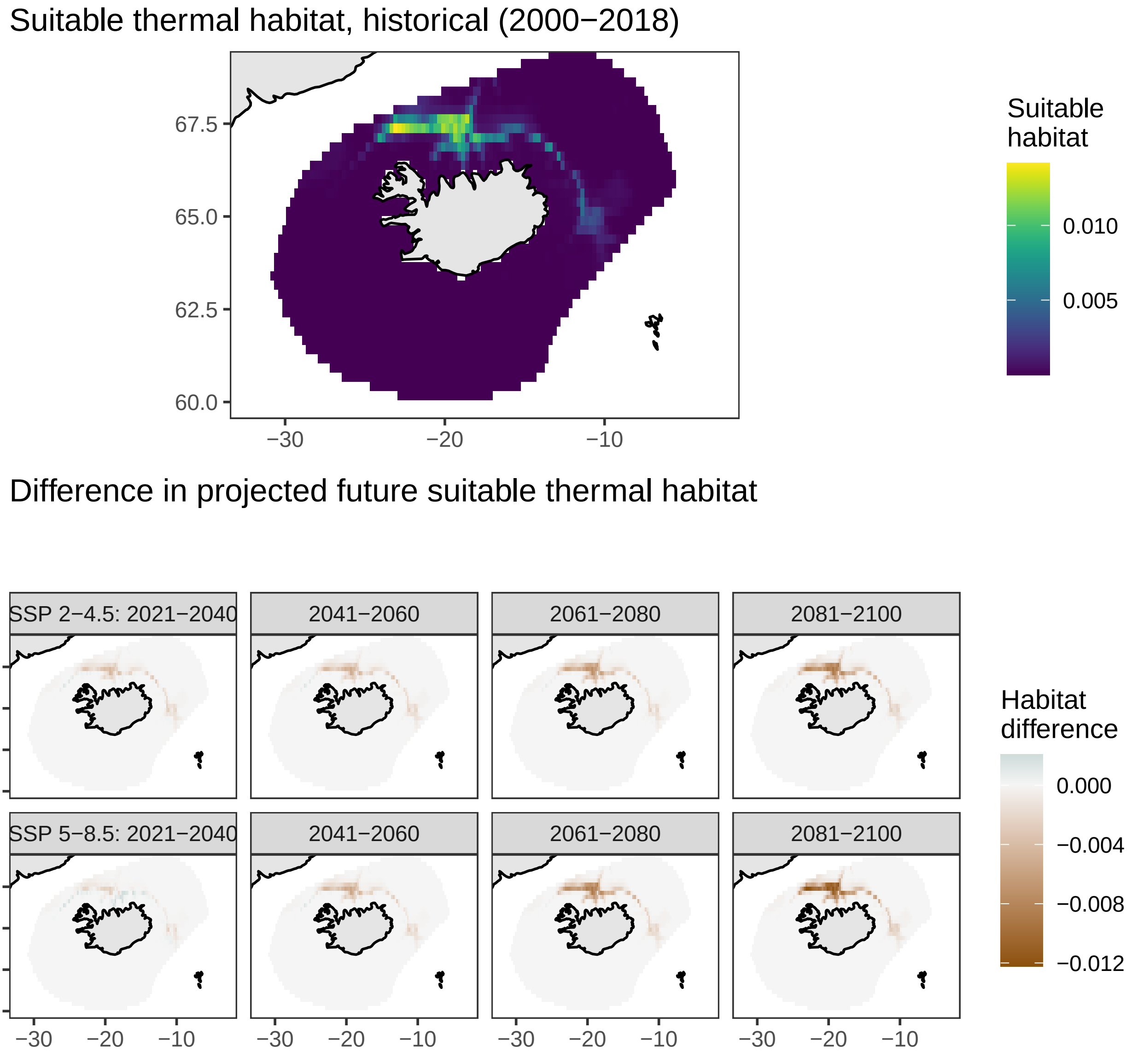

### Spotted snake blenny, *Leptoclinus maculatus*

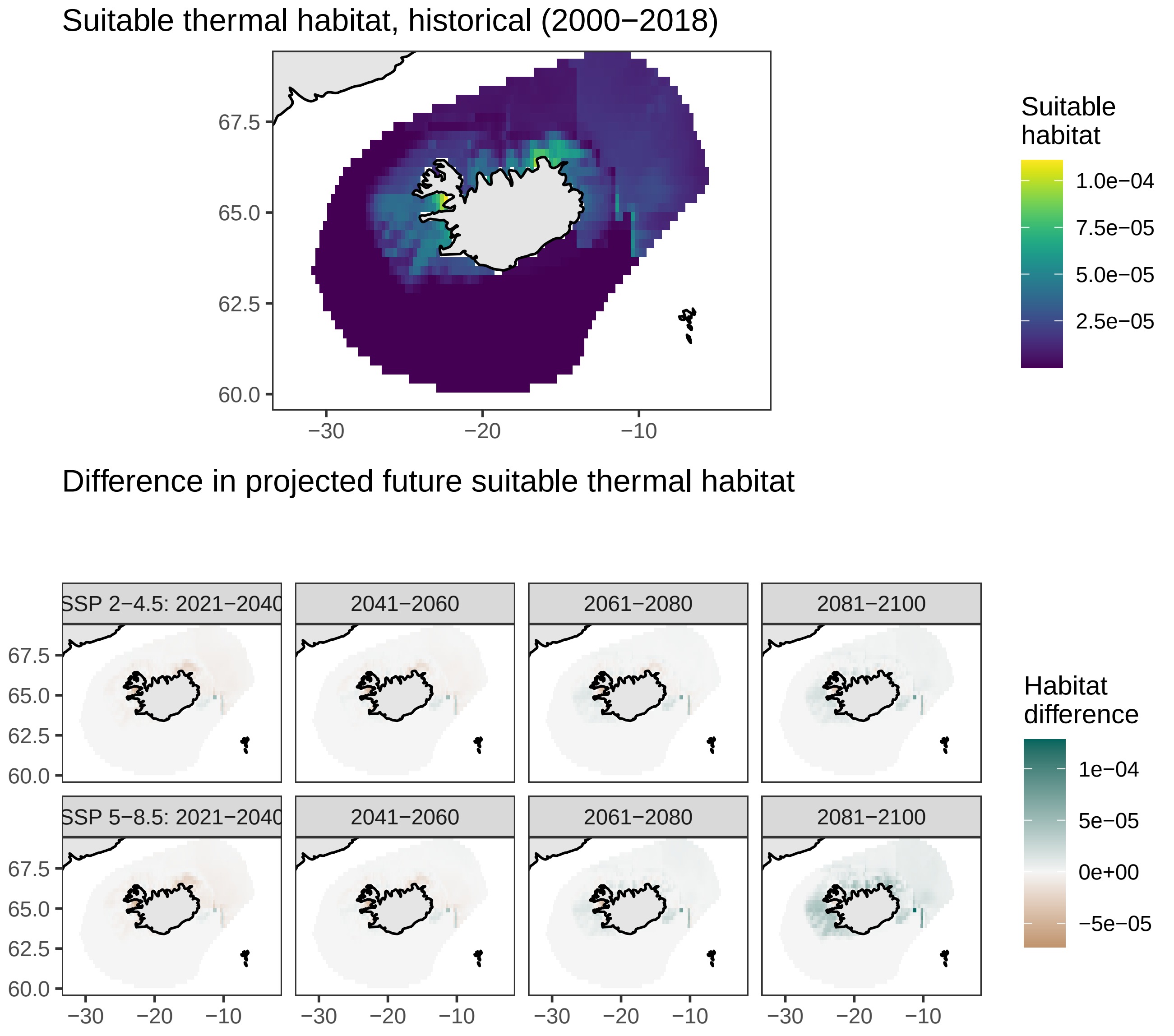

### Dab, *Limanda limanda*

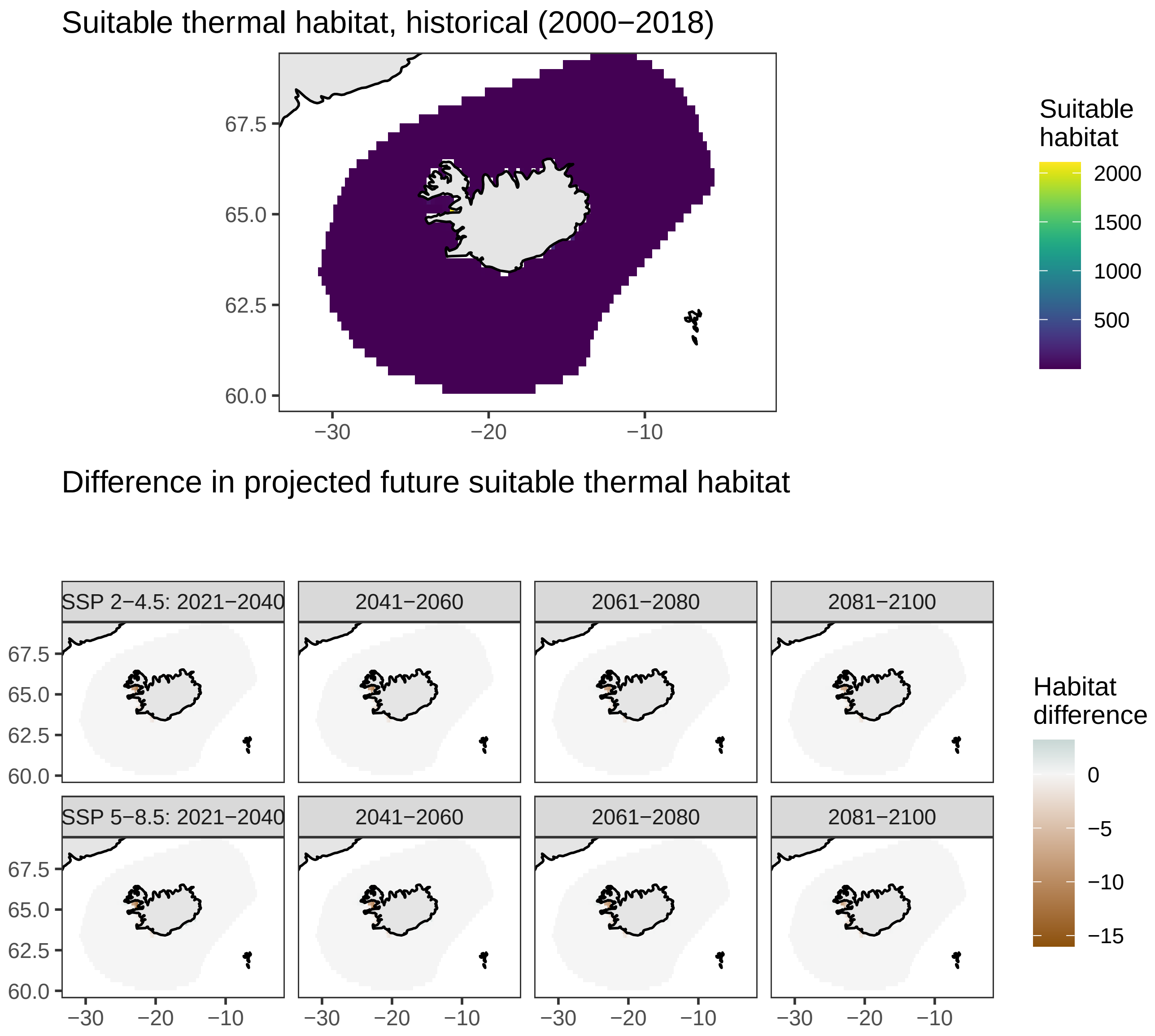

### Monkfish, *Lophius piscatorius*

Suitable thermal habitat, historical (2000−2018)

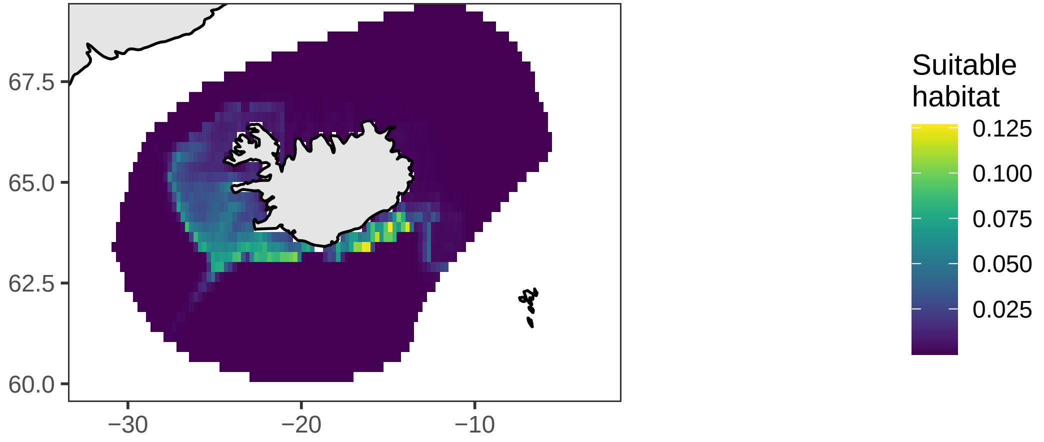


Difference in projected future suitable thermal habitat

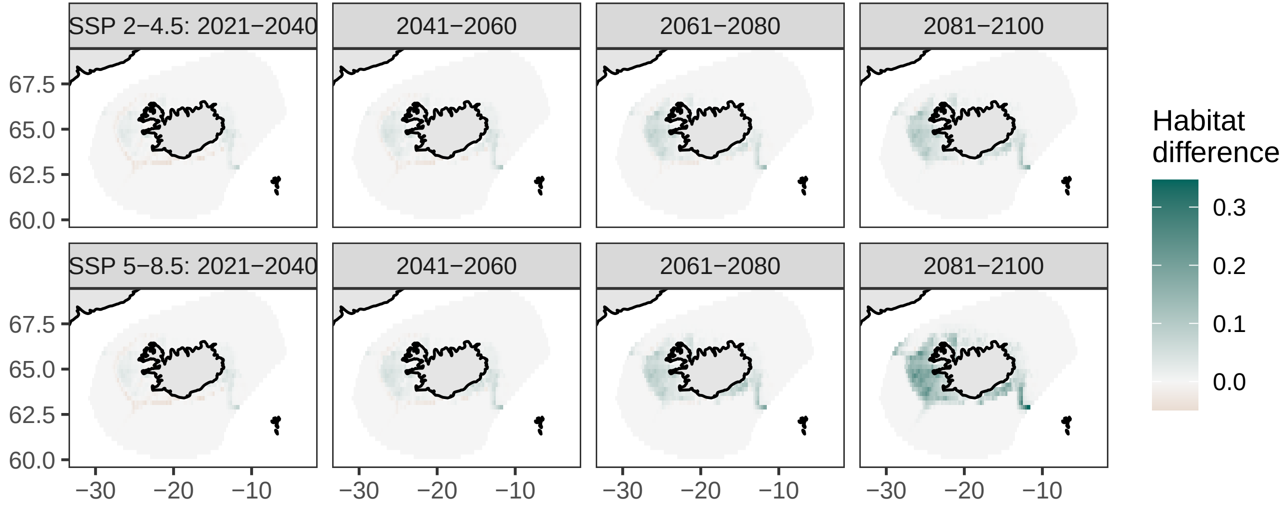

### Esmark's eelpout, *Lycodes esmarkii*

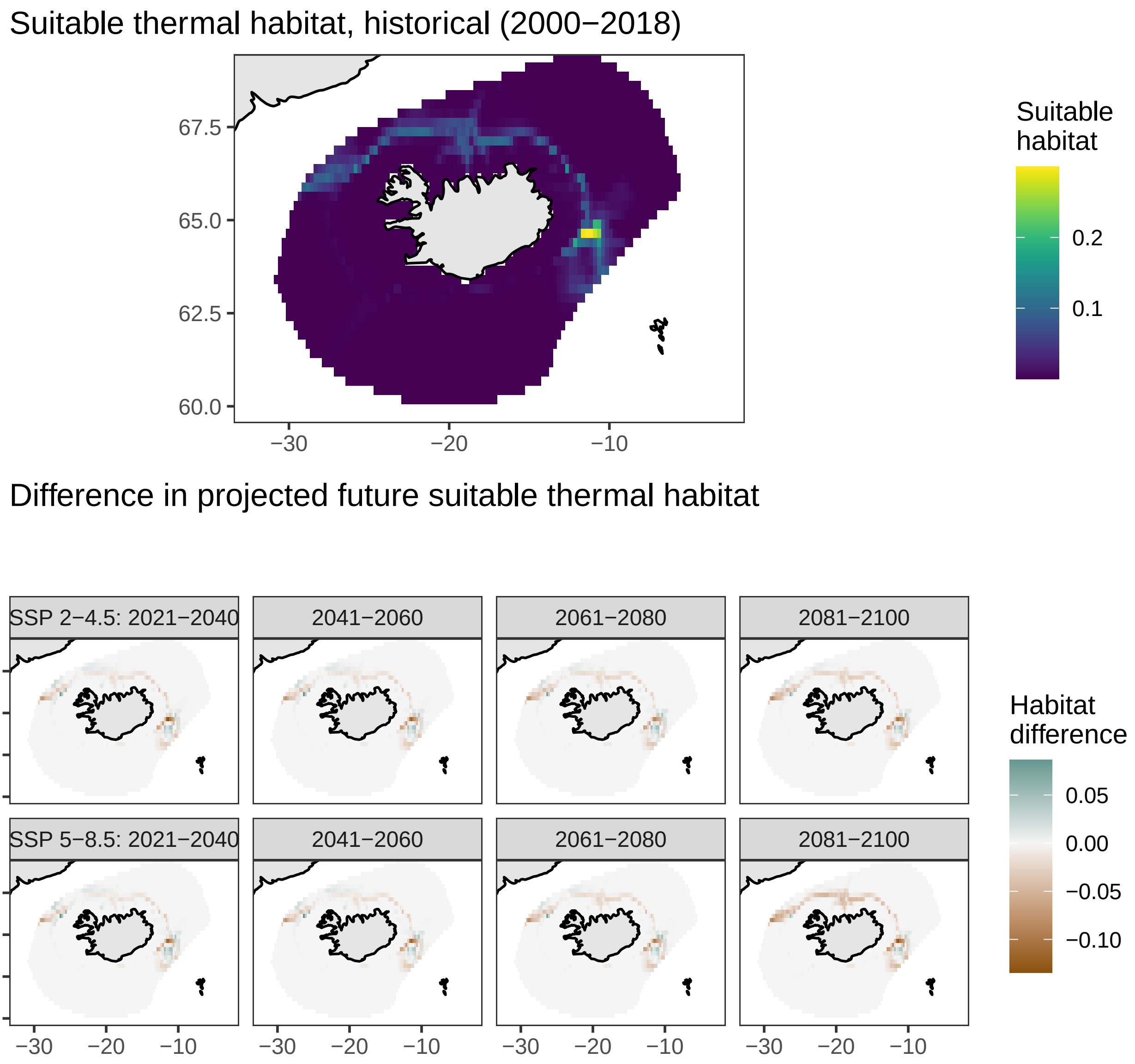

### Lycodes eudipleurostictus, *Lycodes eudipleurostictus*

Suitable thermal habitat, historical (2000−2018)

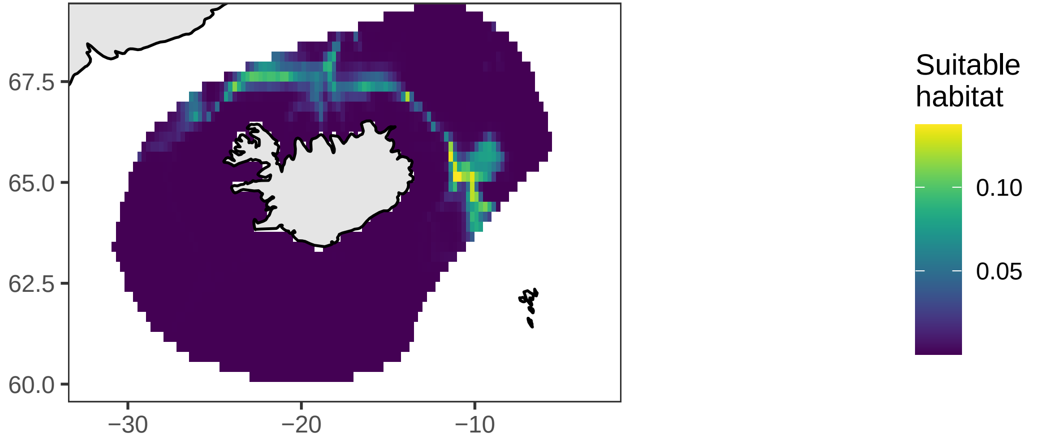


Difference in projected future suitable thermal habitat

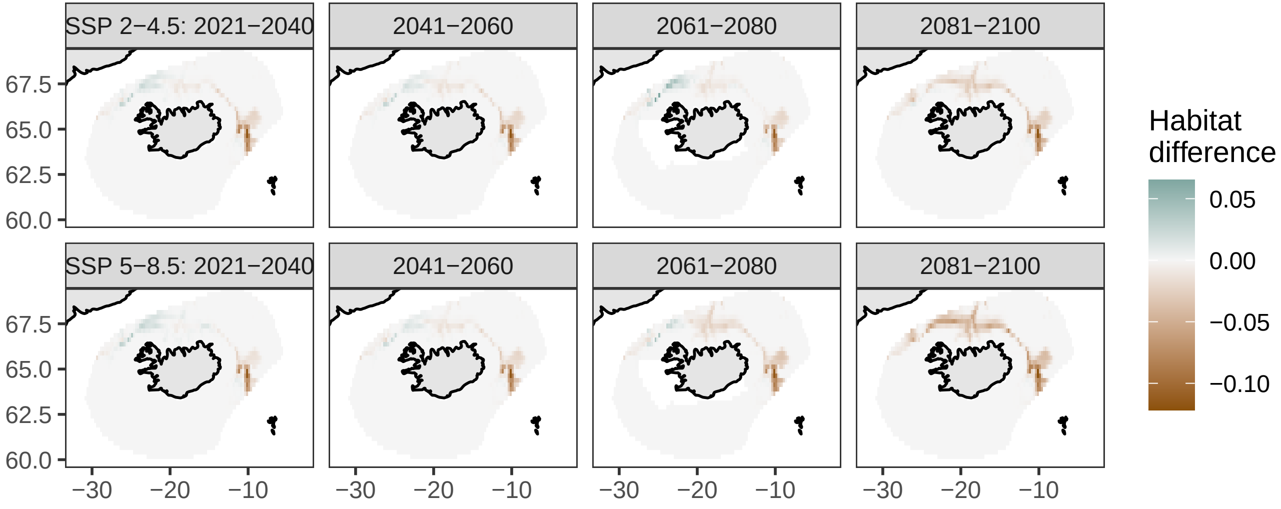

### Vahl's eelpout, checker eelpout, *Lycodes gracilis*

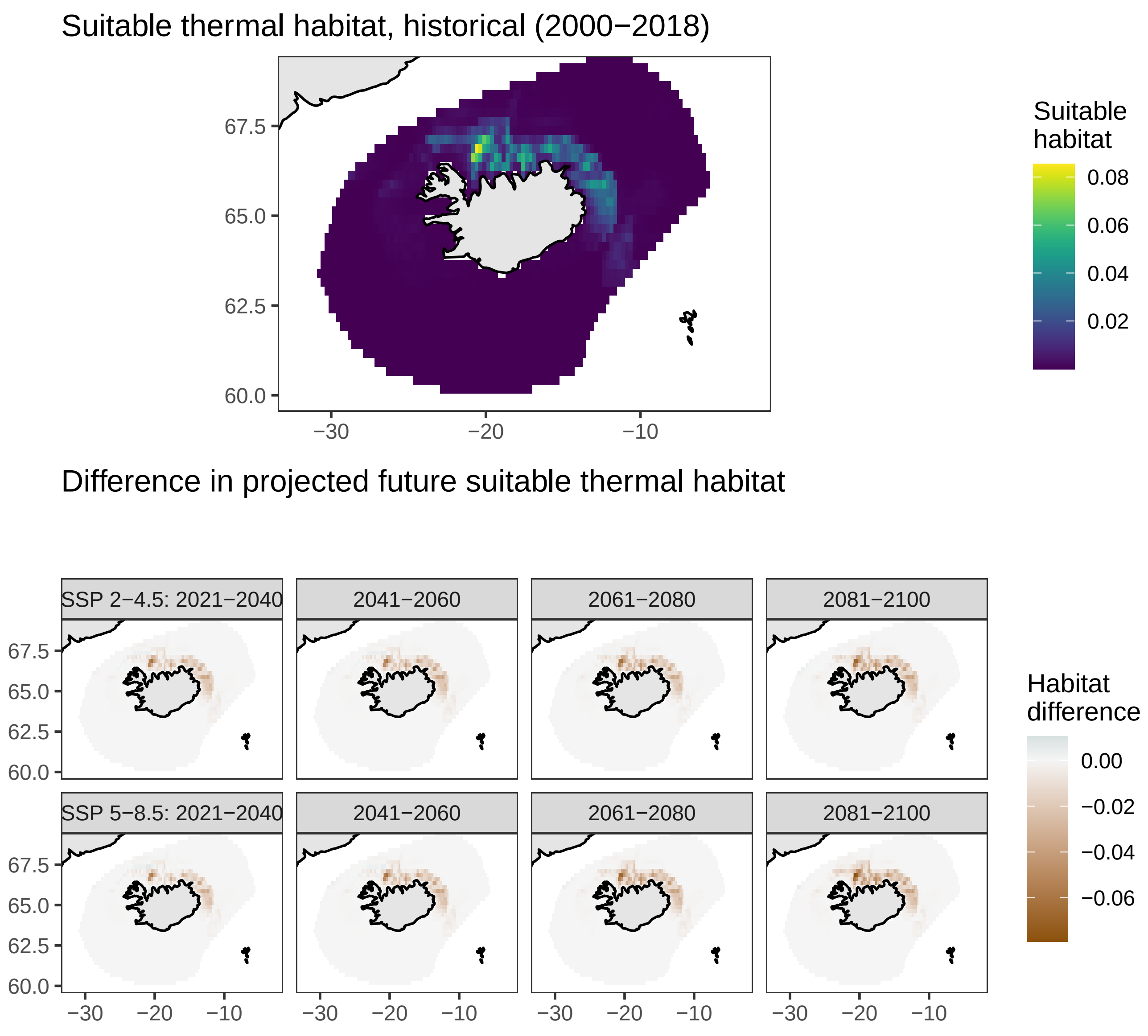

### Arctic eelpout, *Lycodes reticulatus*

#### Suitable thermal habitat, historical (2000−2018)

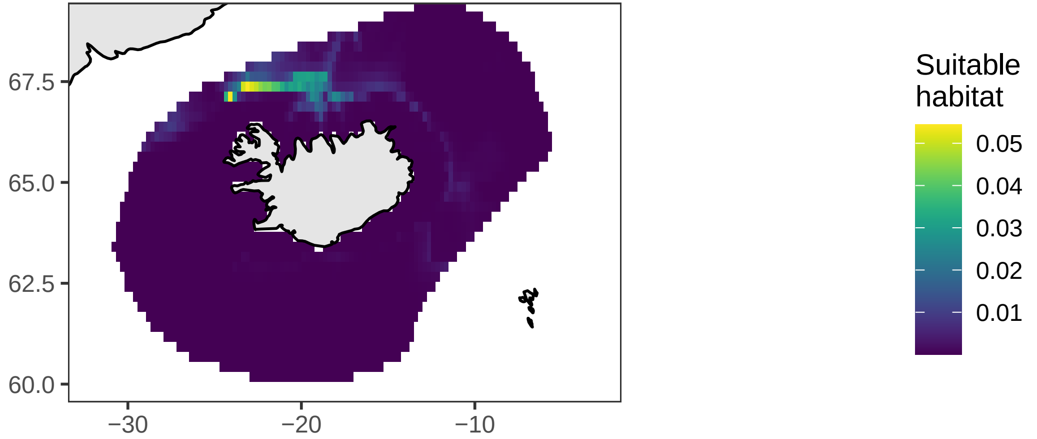


#### Difference in projected future suitable thermal habitat

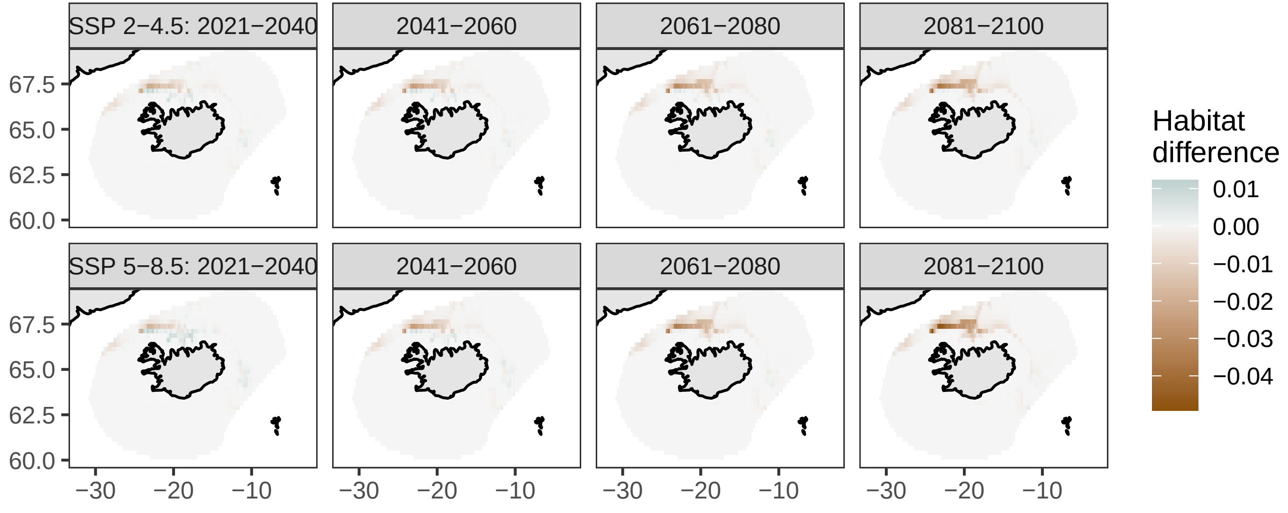

### Lycodes seminudus, *Lycodes seminudus*

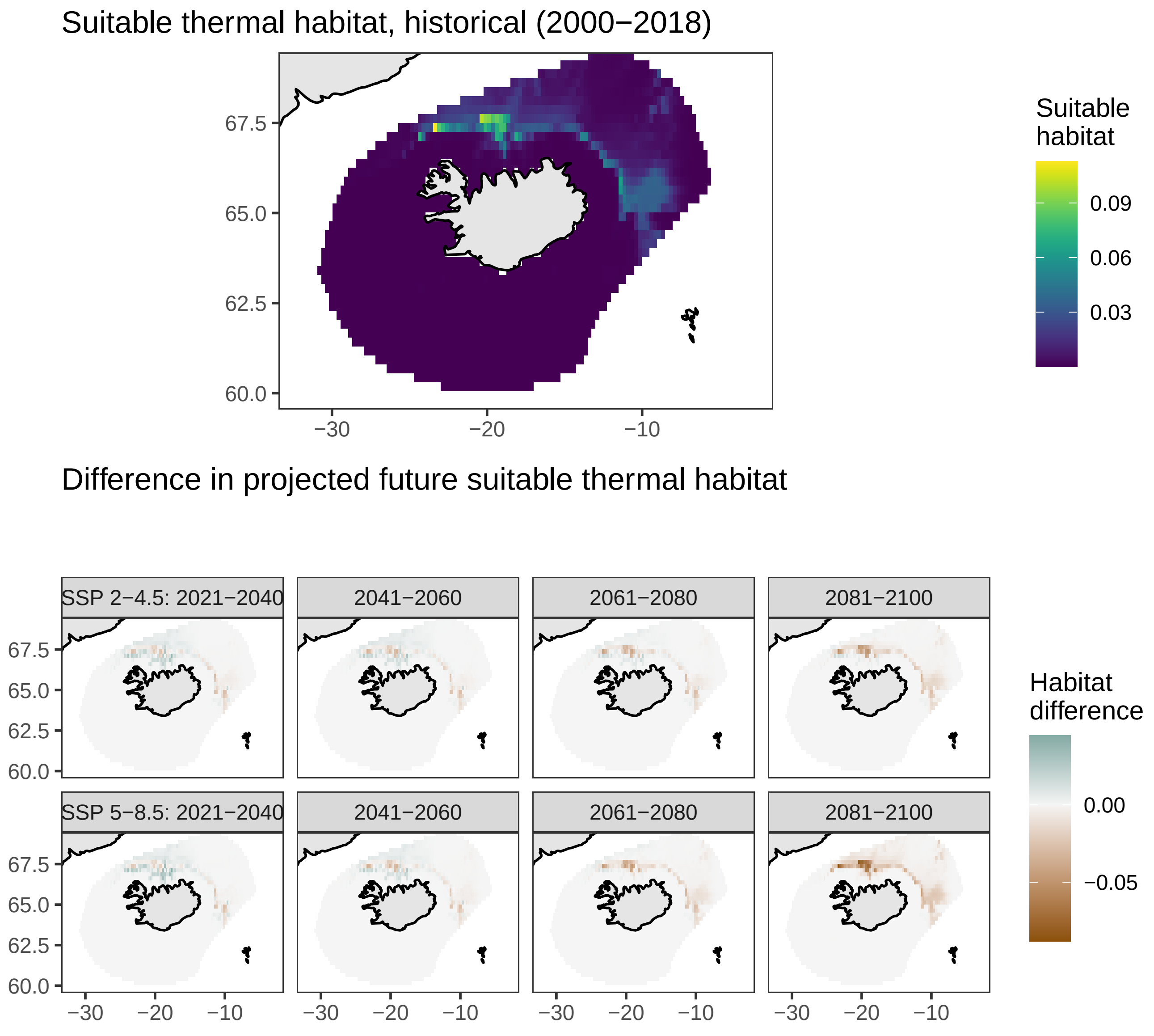

### Roughhead grenadier, *Macrourus berglax*

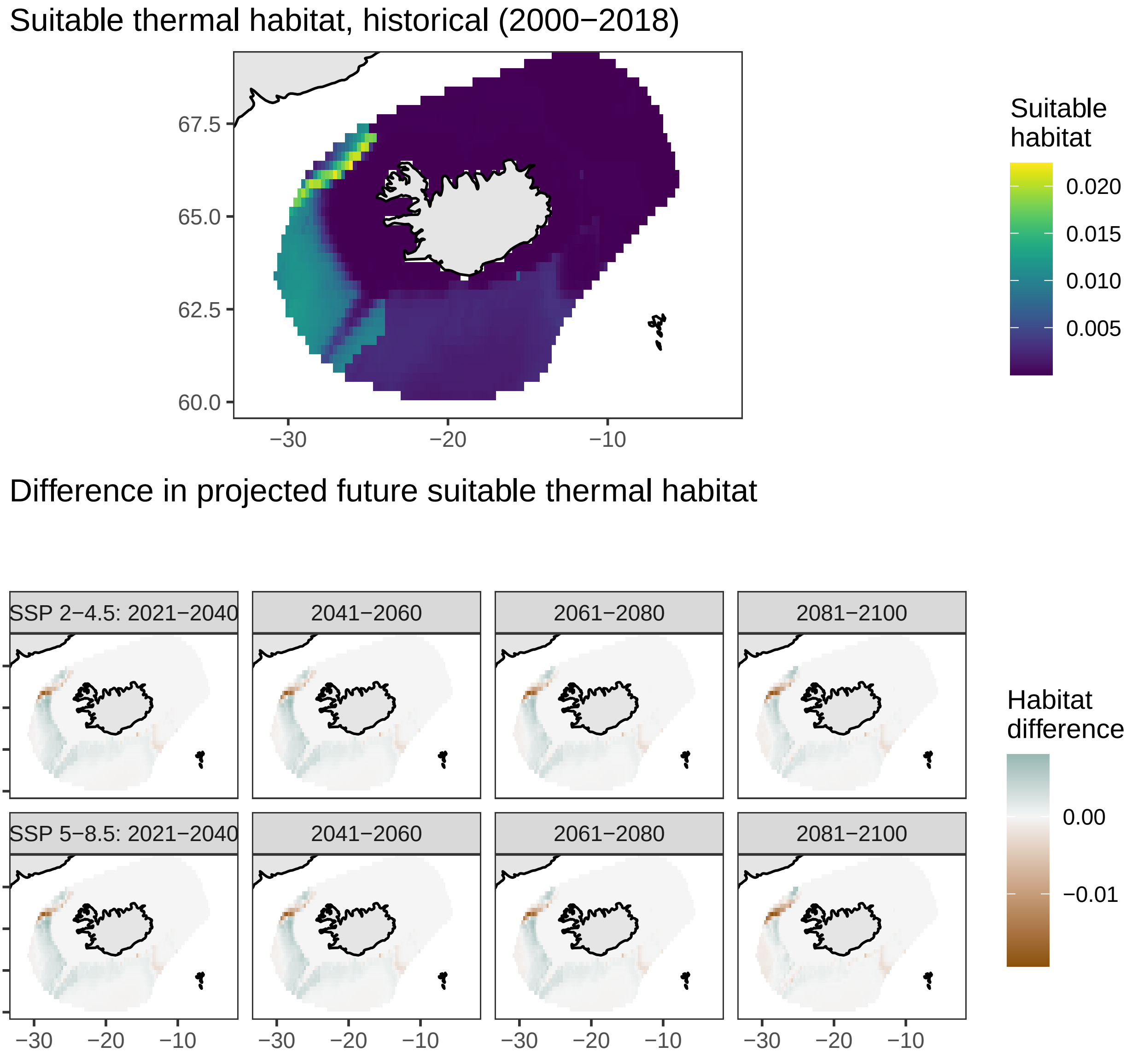

### Whiting, *Merlangius merlangus*

Suitable thermal habitat, historical (2000−2018)

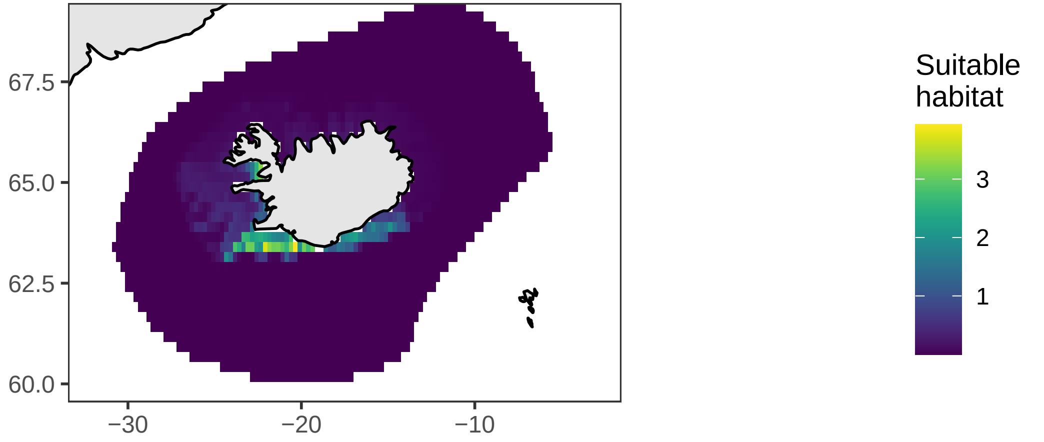


Difference in projected future suitable thermal habitat

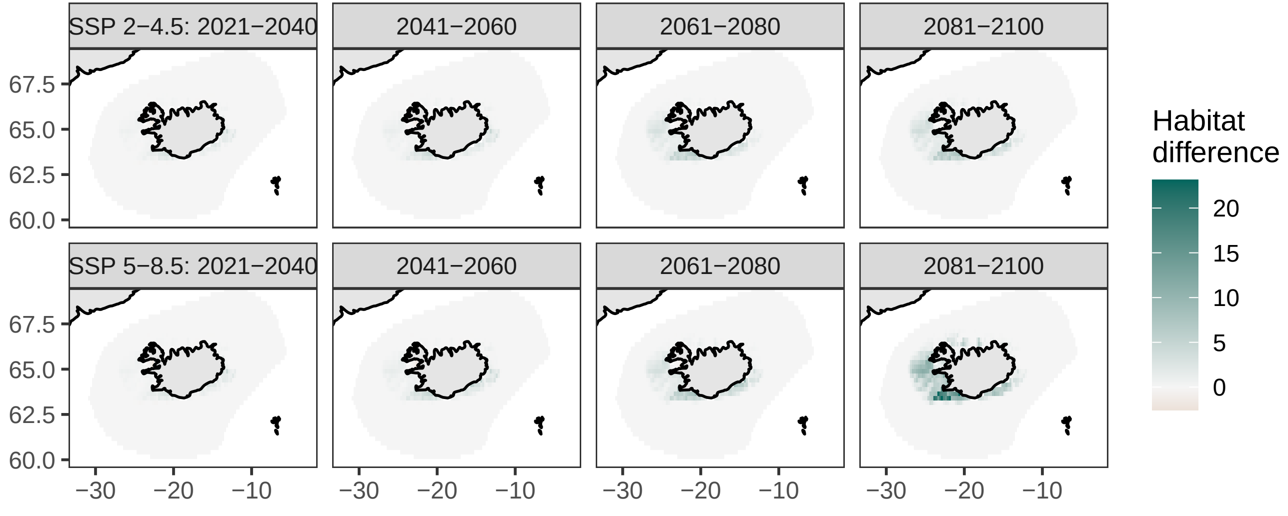

### Lemon sole, *Microstomus kitt*

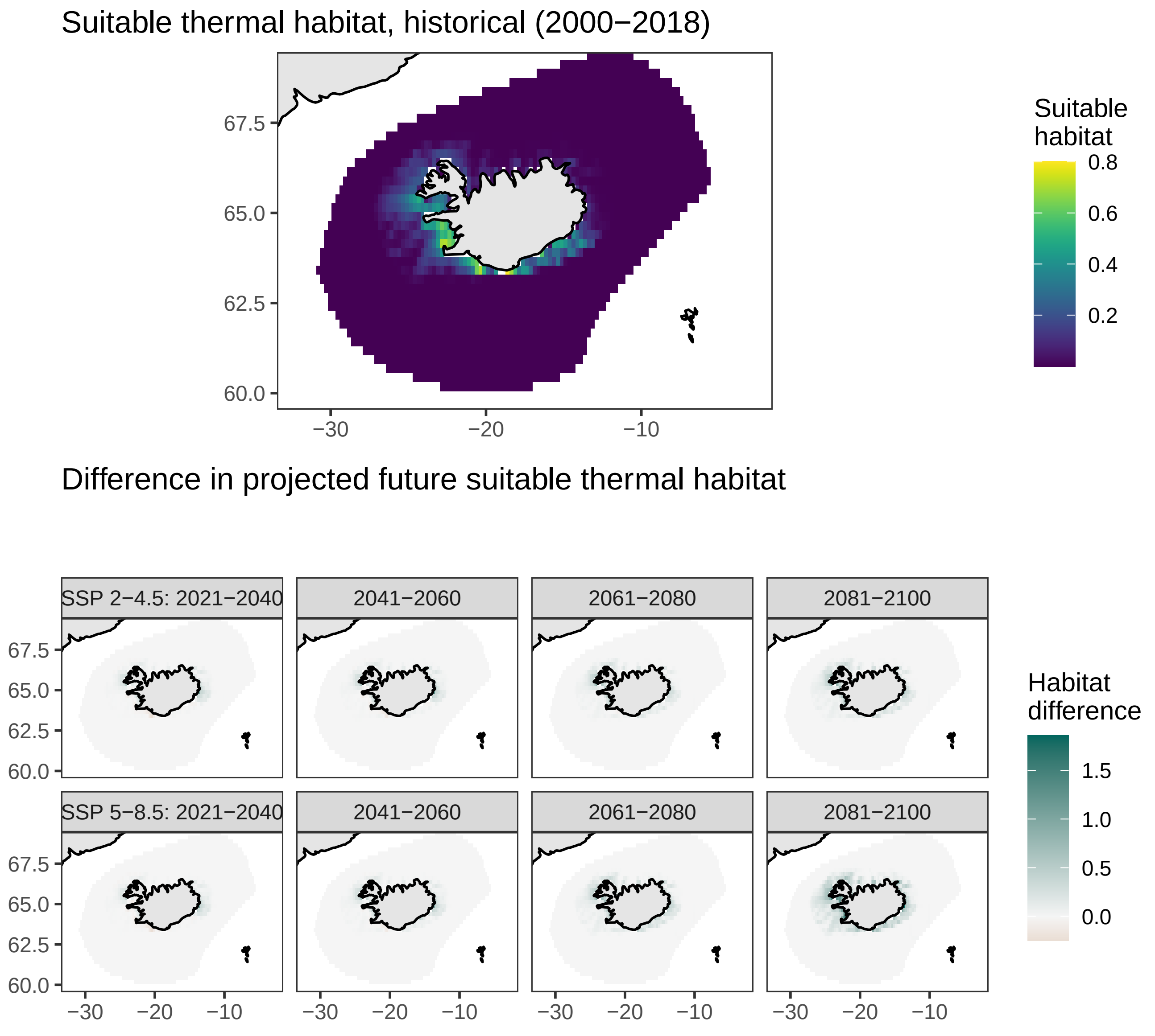

### Blue ling, European ling, *Molva dypterygia*

Suitable thermal habitat, historical (2000−2018)

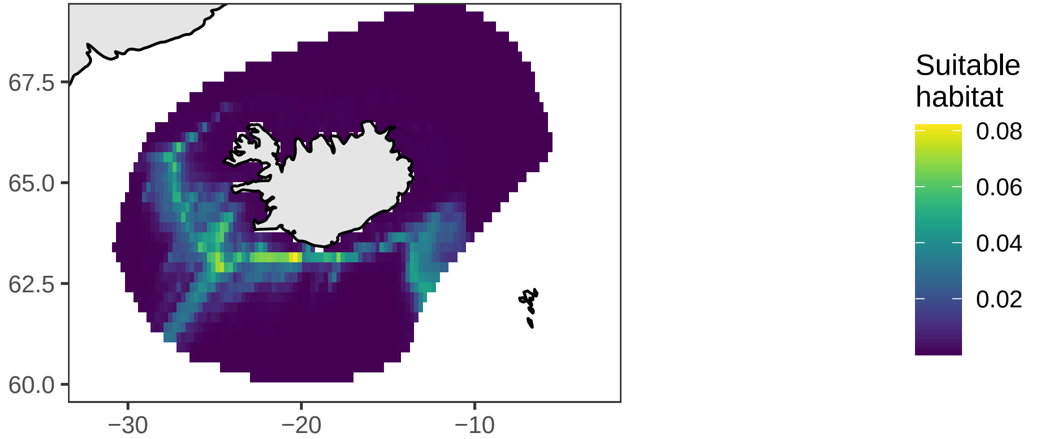


Difference in projected future suitable thermal habitat

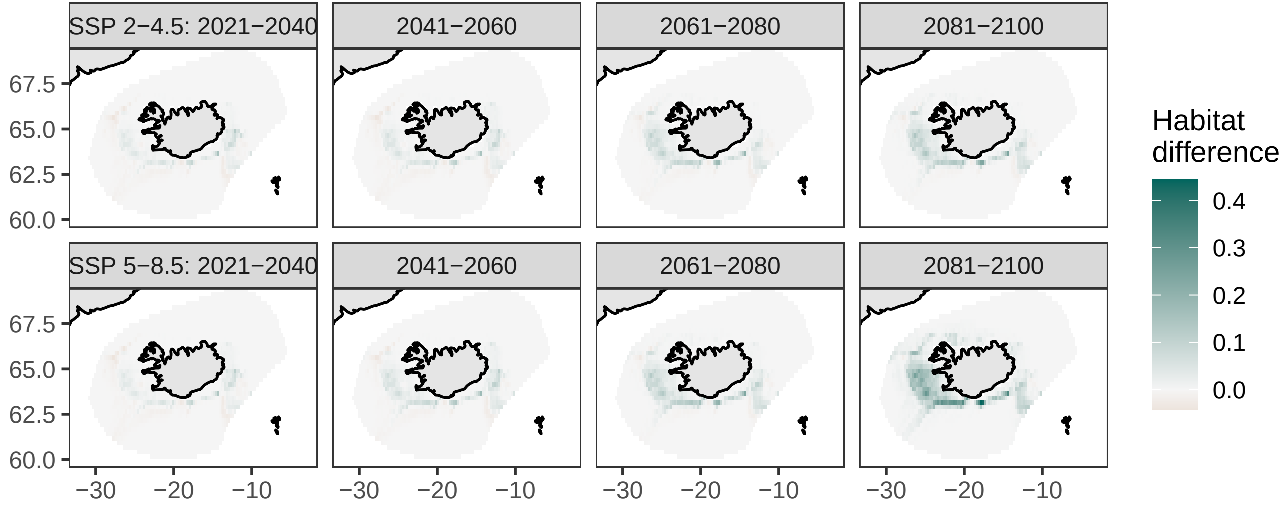

### Ling, *Molva molva*

Suitable thermal habitat, historical (2000−2018)

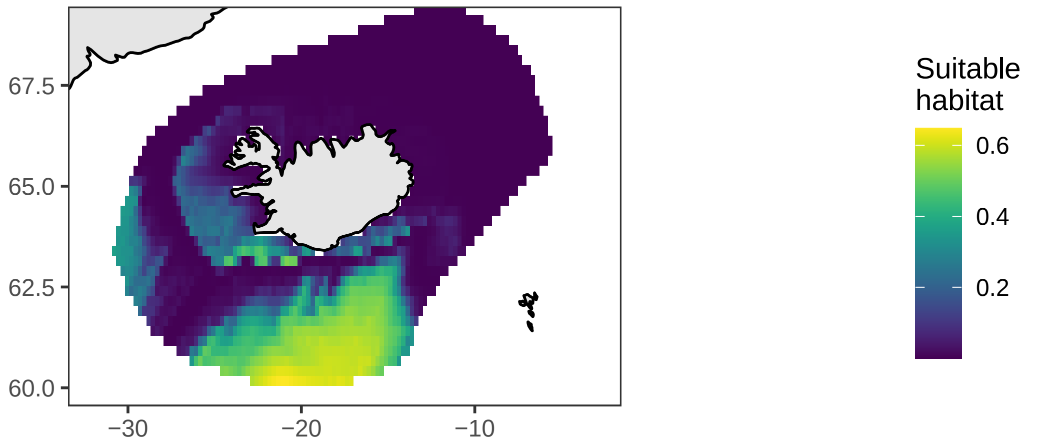


Difference in projected future suitable thermal habitat

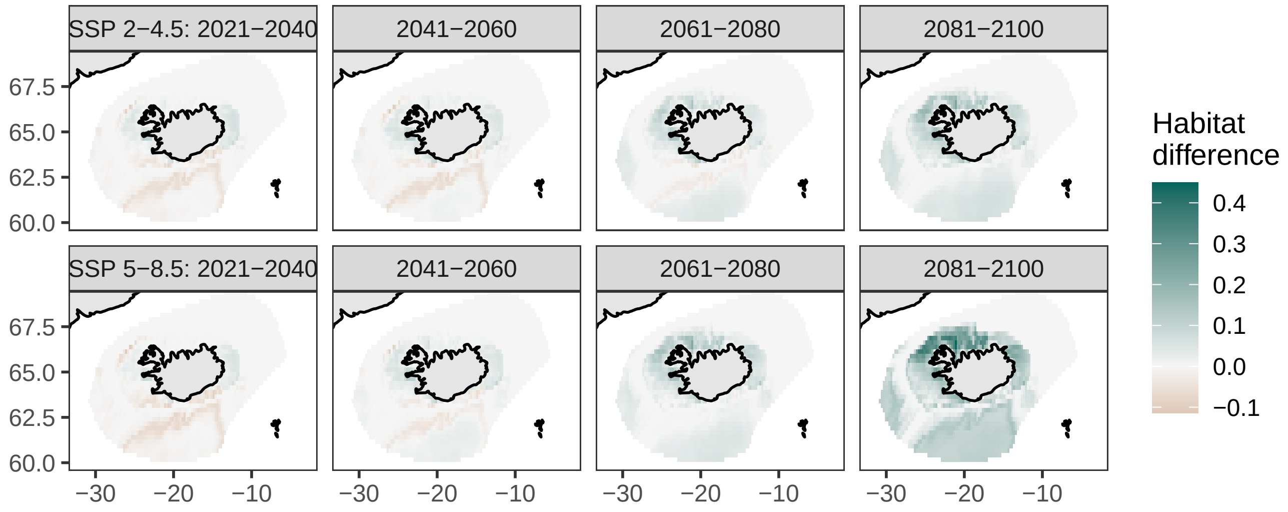

### Greater forkbeard, *Phycis blennoides*

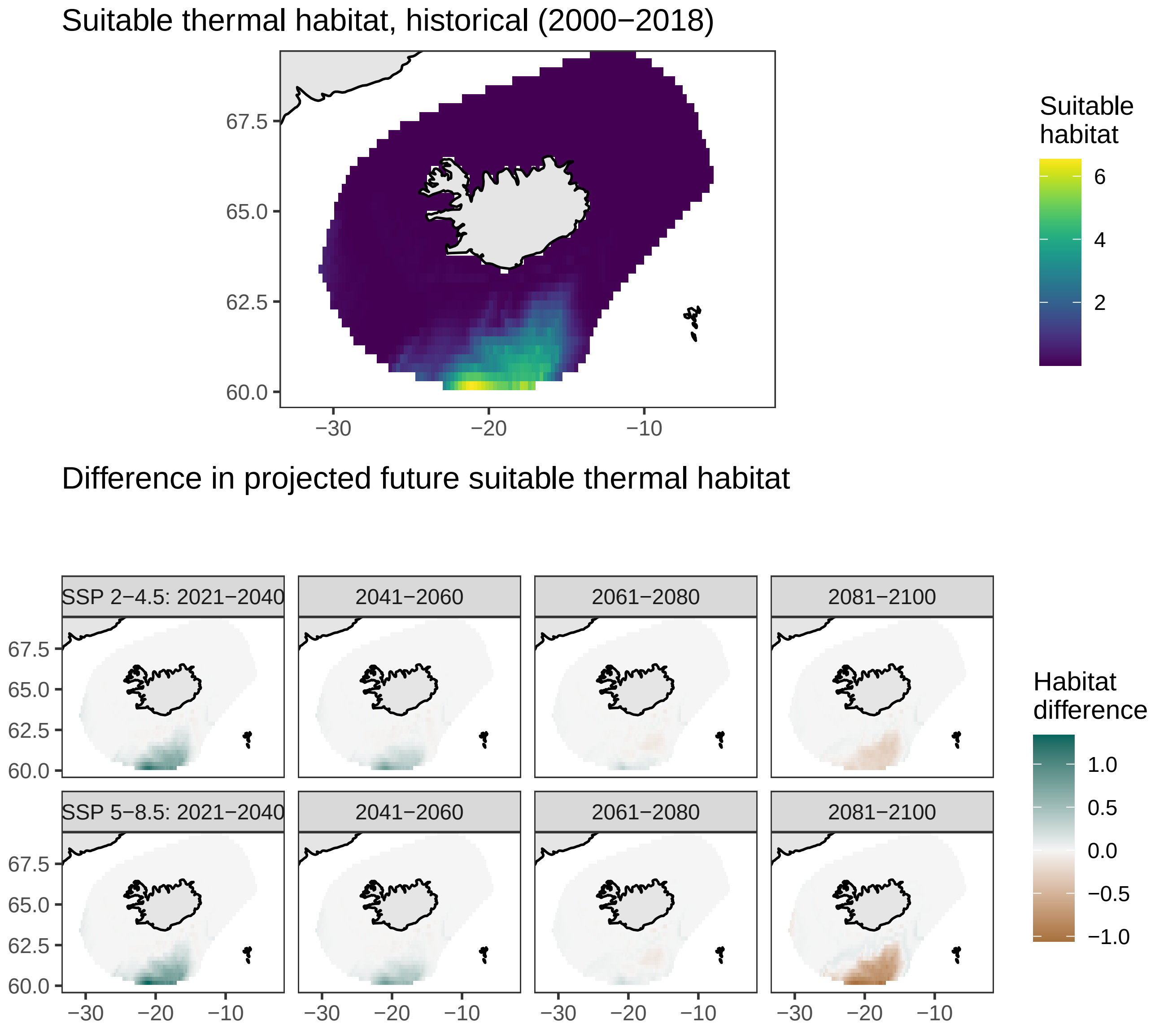

### Plaice, *Pleuronectes platessa*

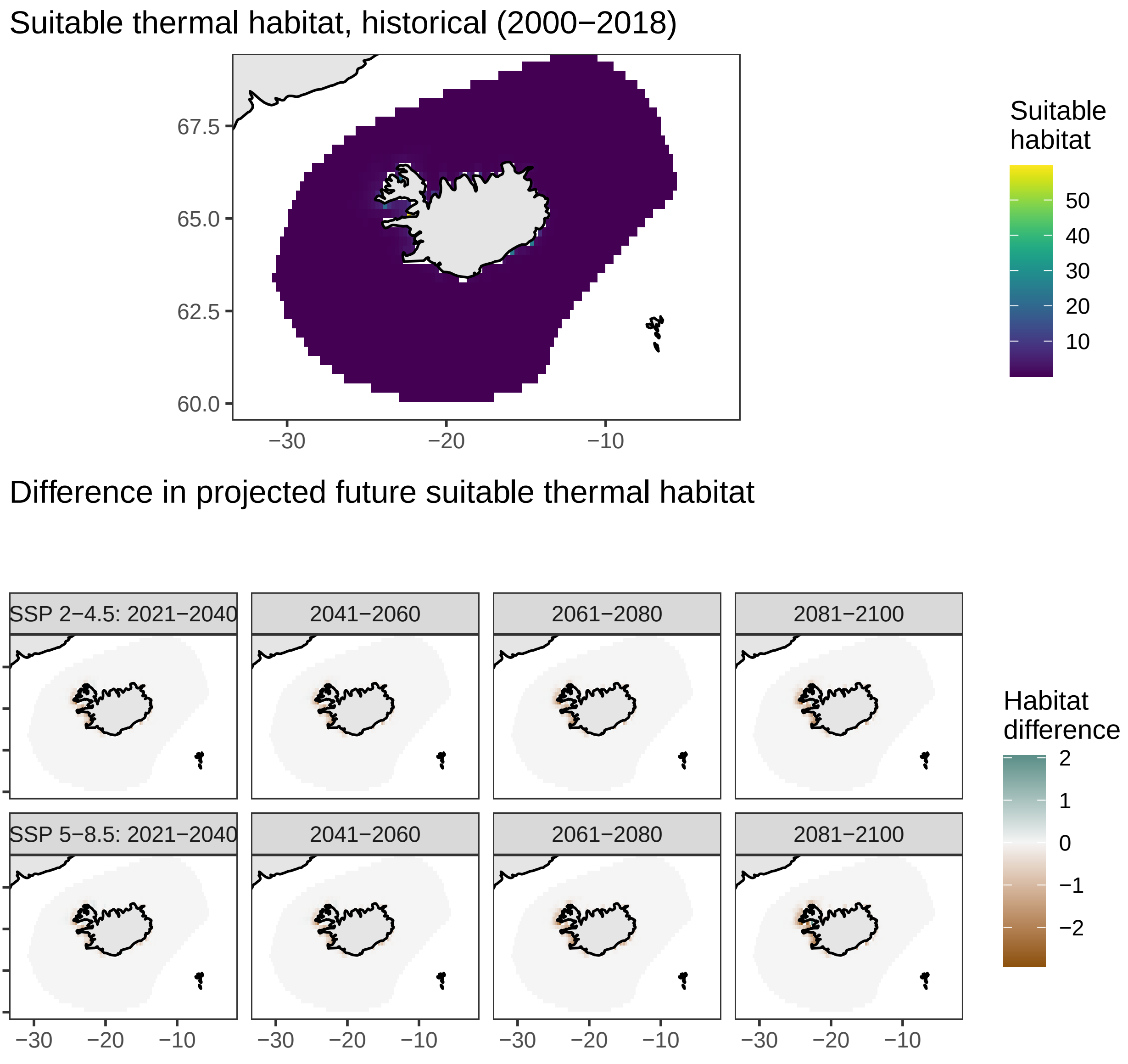

### Saithe, *Pollachius virens*

#### Suitable thermal habitat, historical (2000−2018)

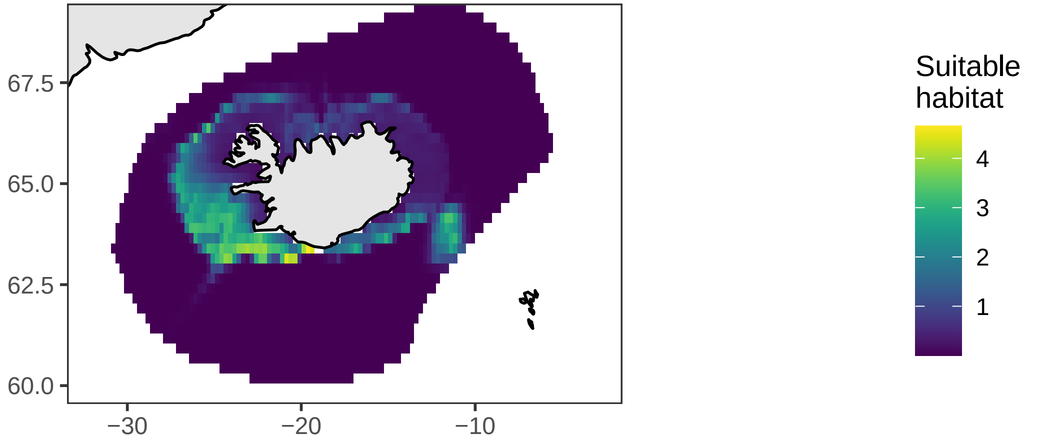


#### Difference in projected future suitable thermal habitat

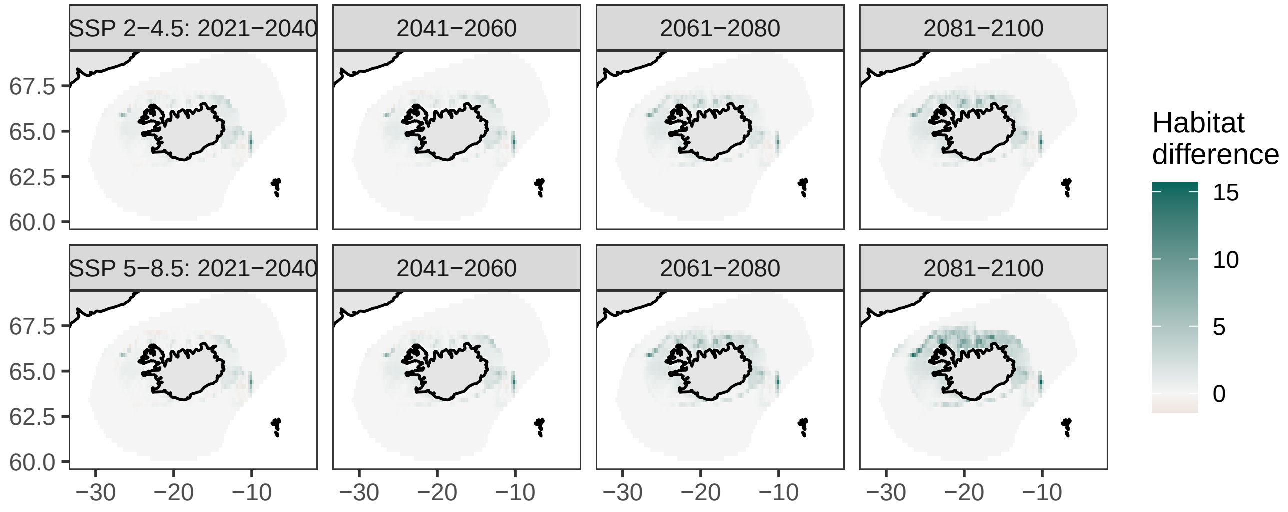

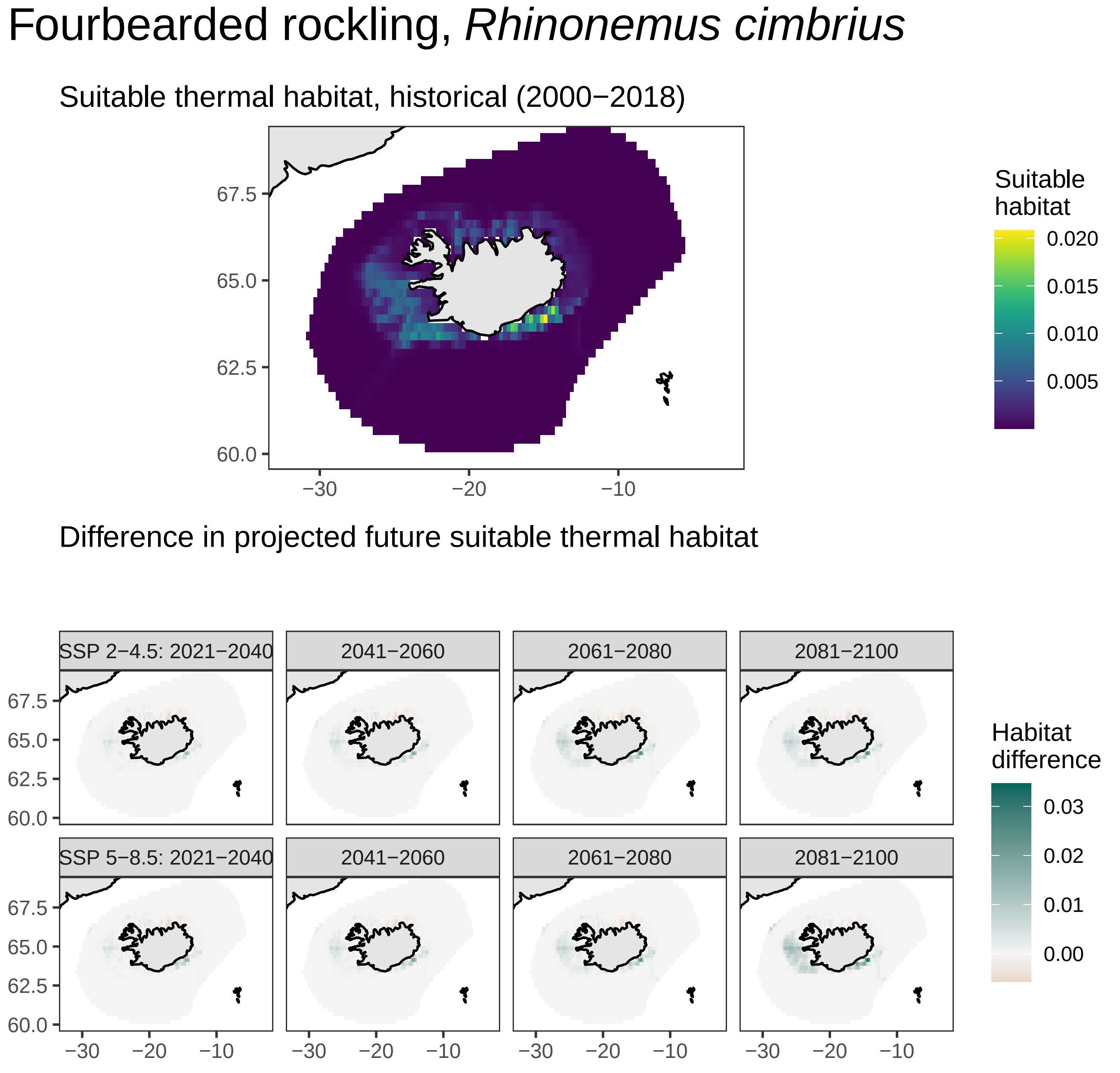
Fourbearded rockling, *Rhinonemus cimbrius*
Suitable thermal habitat, historical (2000−2018)
Suitable habitat
0.020
0.015
0.010
0.005
67.5
65.0
62.5
60.0
−30
−20
−10
Difference in projected future suitable thermal habitat
SSP 2−4.5: 2021−2040
2041−2060
2061−2080
2081−2100
SSP 5−8.5: 2021−2040
2041−2060
2061−2080
2081−2100
Habitat difference
0.03
0.02
0.01
0.00

### Redfish, *Sebastes marinus*

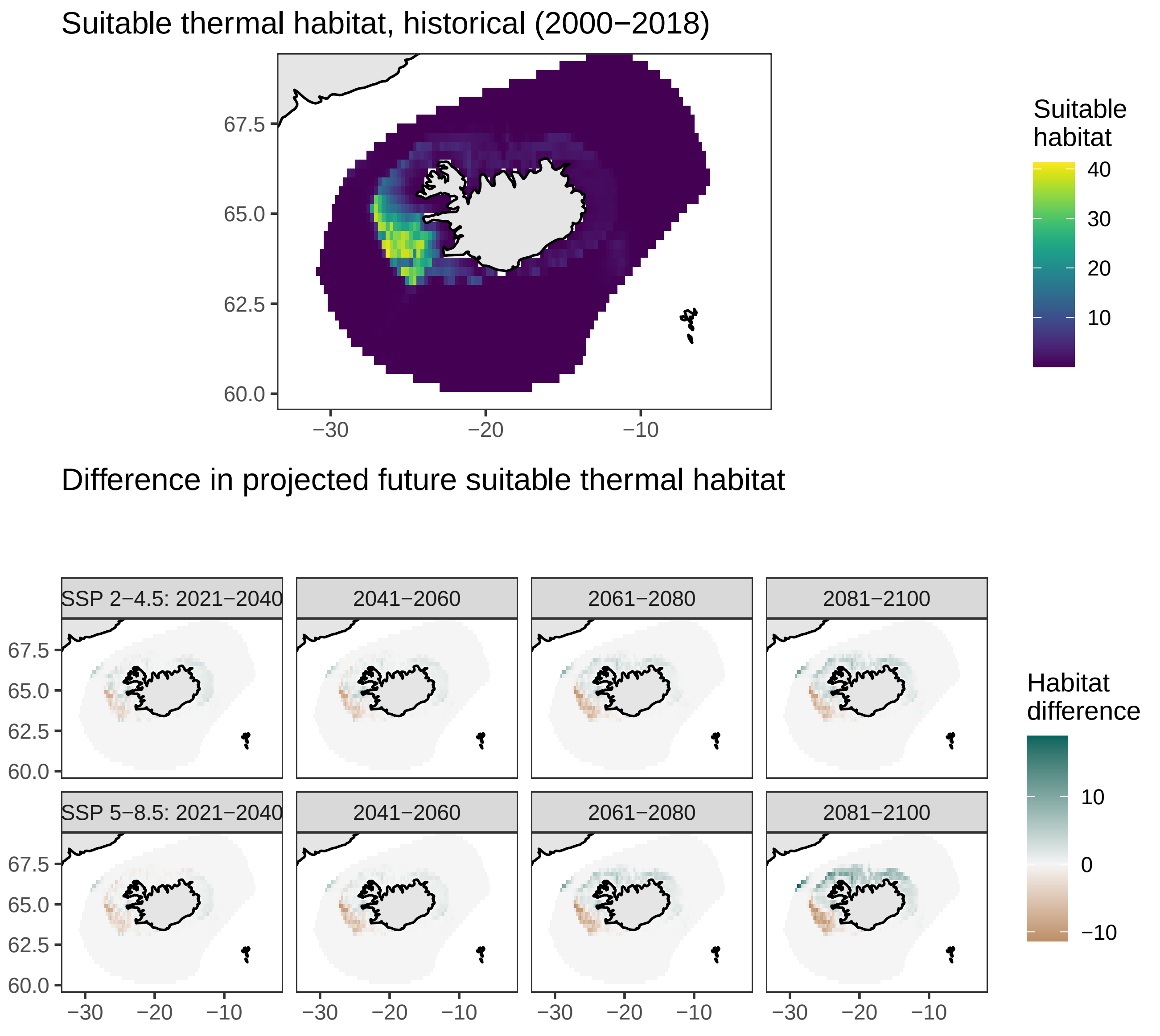

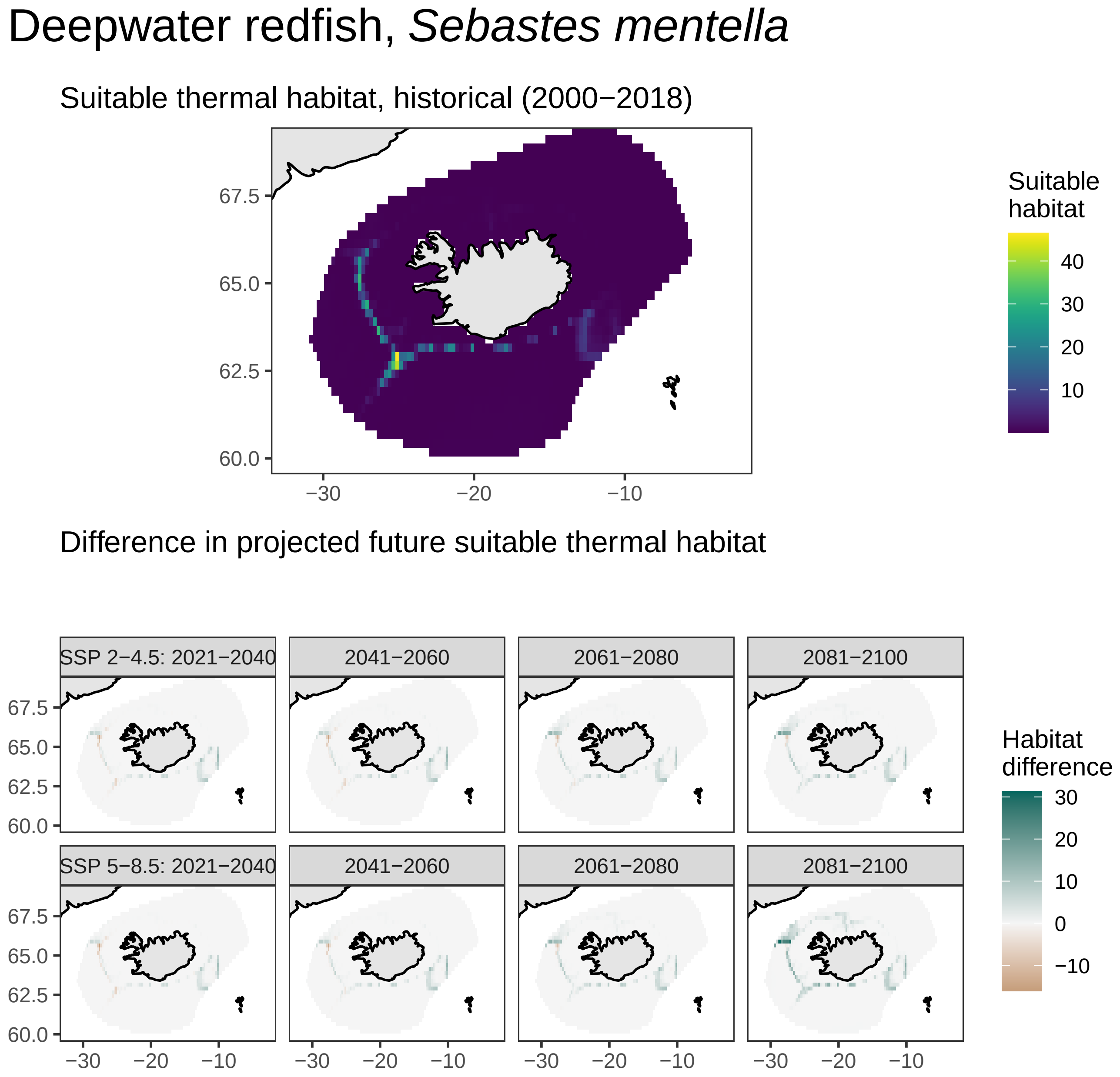
Deepwater redfish, Sebastes mentella
Suitable thermal habitat, historical (2000−2018)
Suitable habitat
40
30
20
10
67.5
65.0
62.5
60.0
−30
−20
−10
Difference in projected future suitable thermal habitat
SSP 2−4.5: 2021−2040
2041−2060
2061−2080
2081−2100
SSP 5−8.5: 2021−2040
2041−2060
2061−2080
2081−2100
Habitat difference
30
20
10
0
−10

### Norway haddock, *Sebastes viviparus*

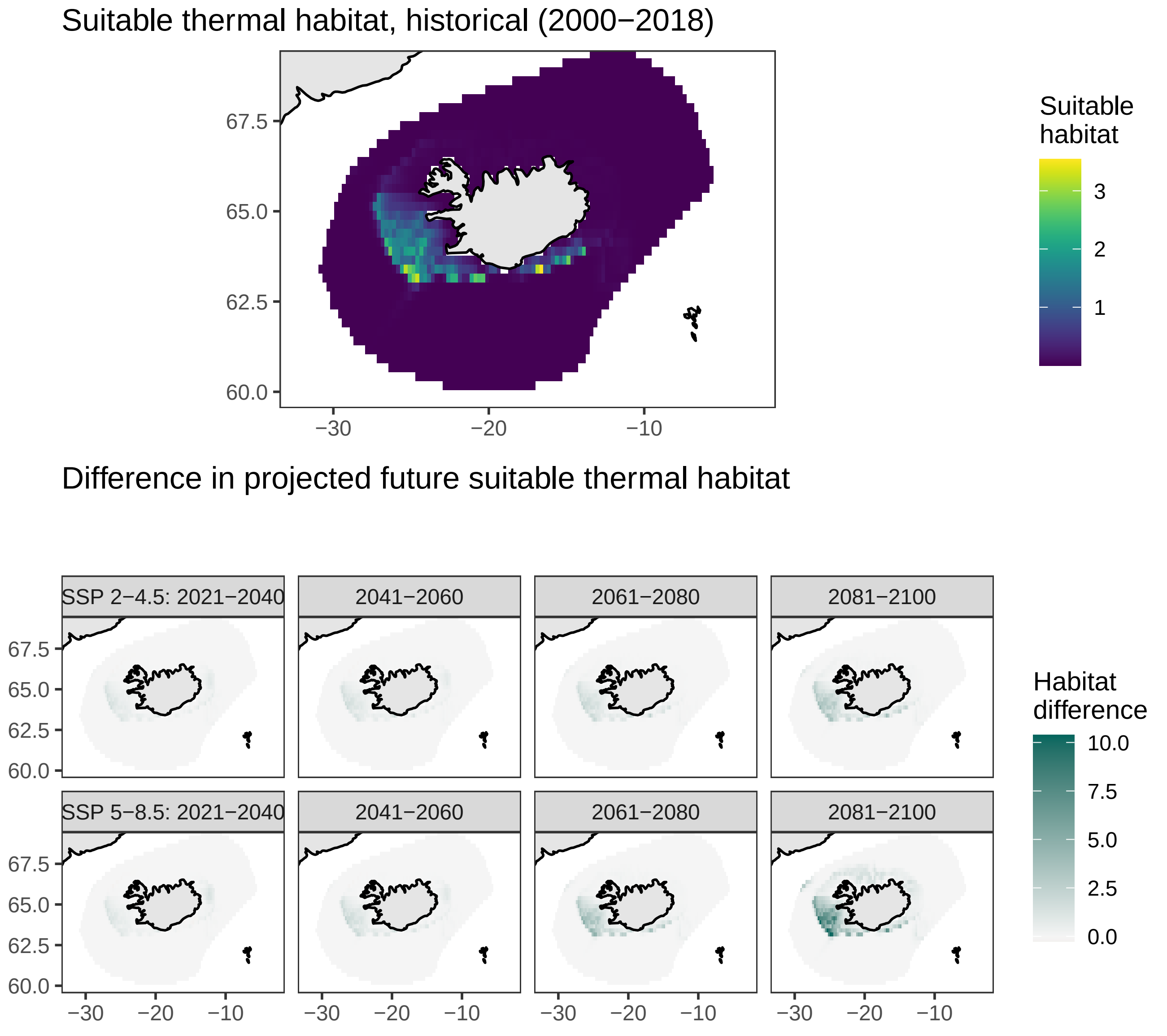

### Squid, *Squid*

Suitable thermal habitat, historical (2000−2018)

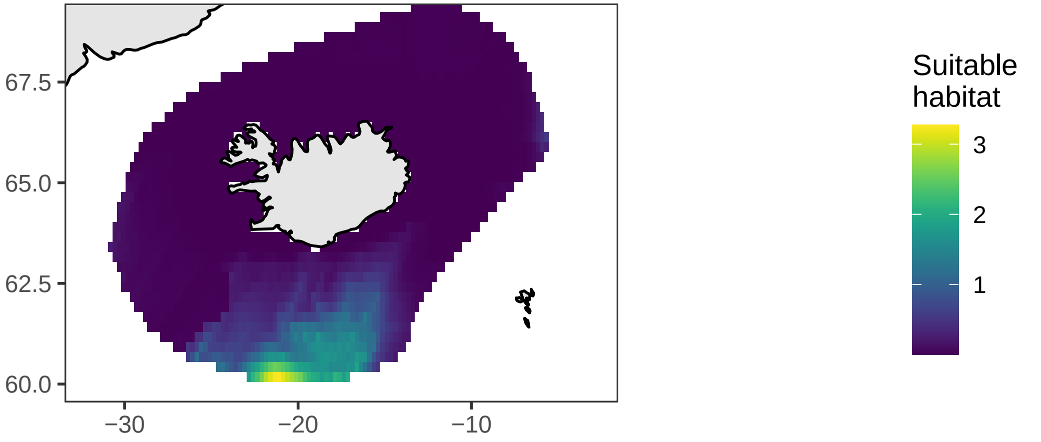


Difference in projected future suitable thermal habitat

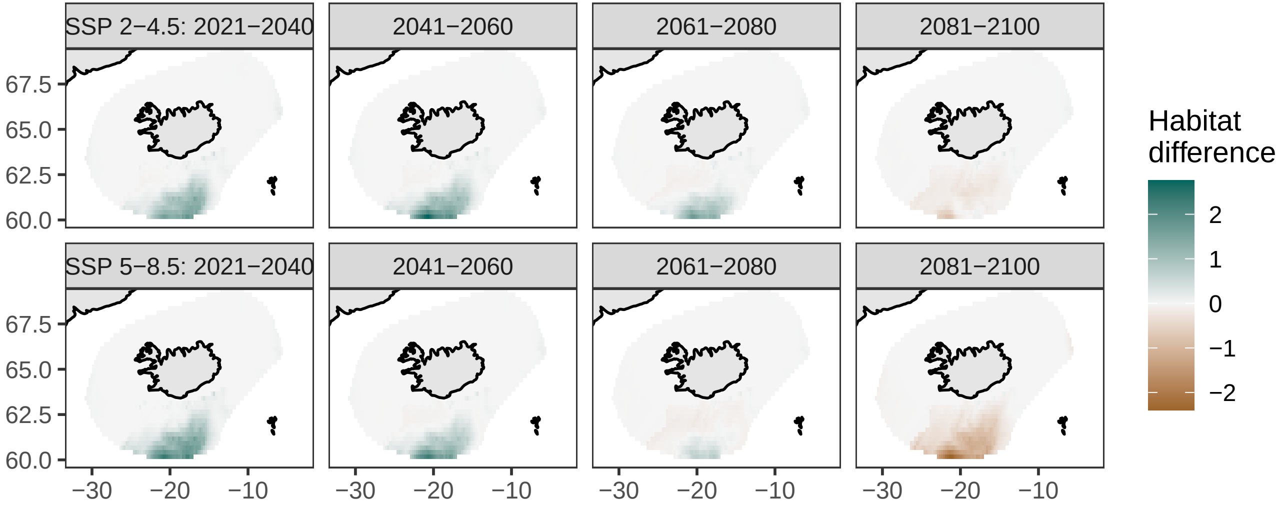

### Moustache sculpin, *Triglops murrayi*

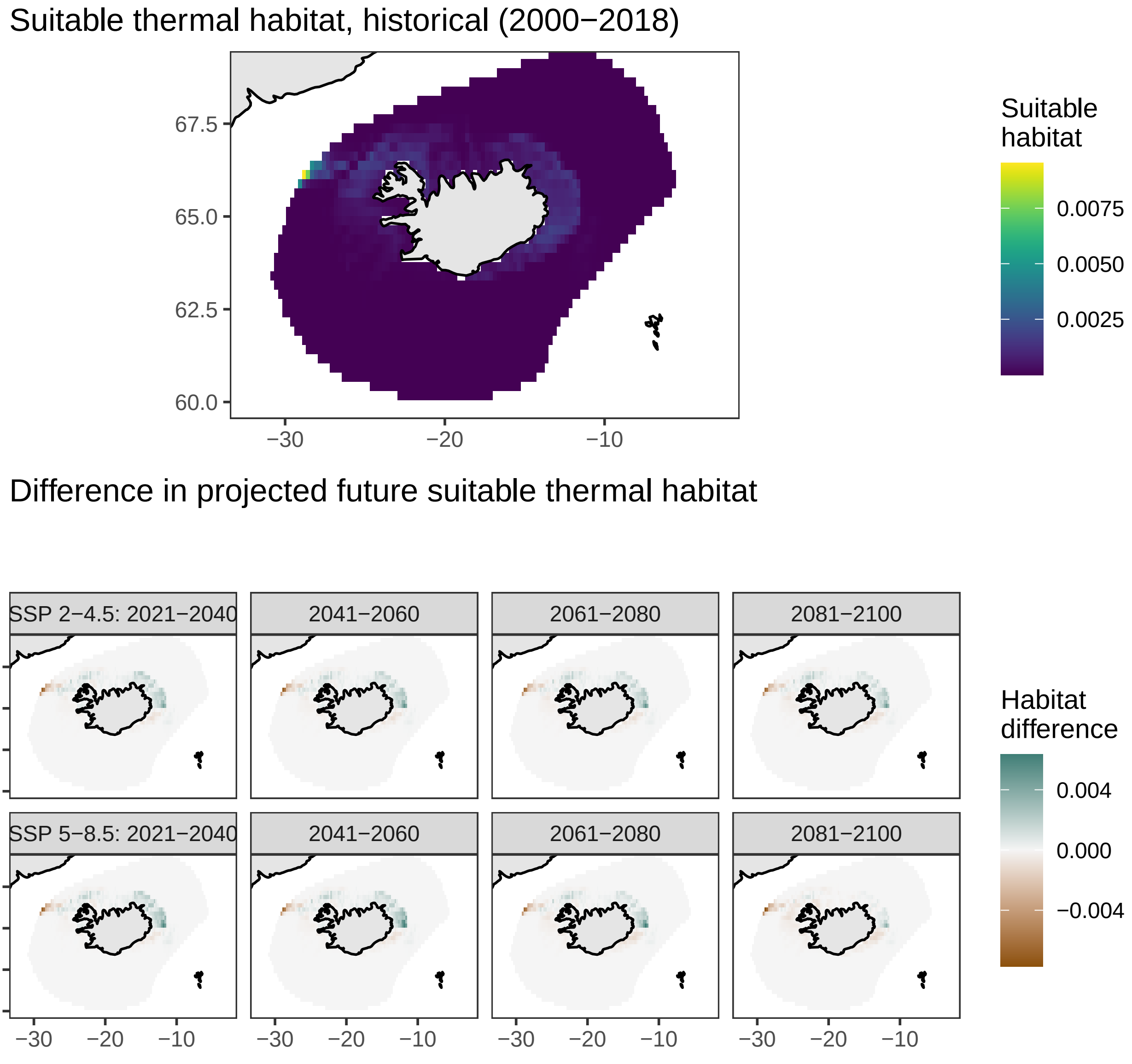

### Norway pout, *Trisopterus esmarkii*

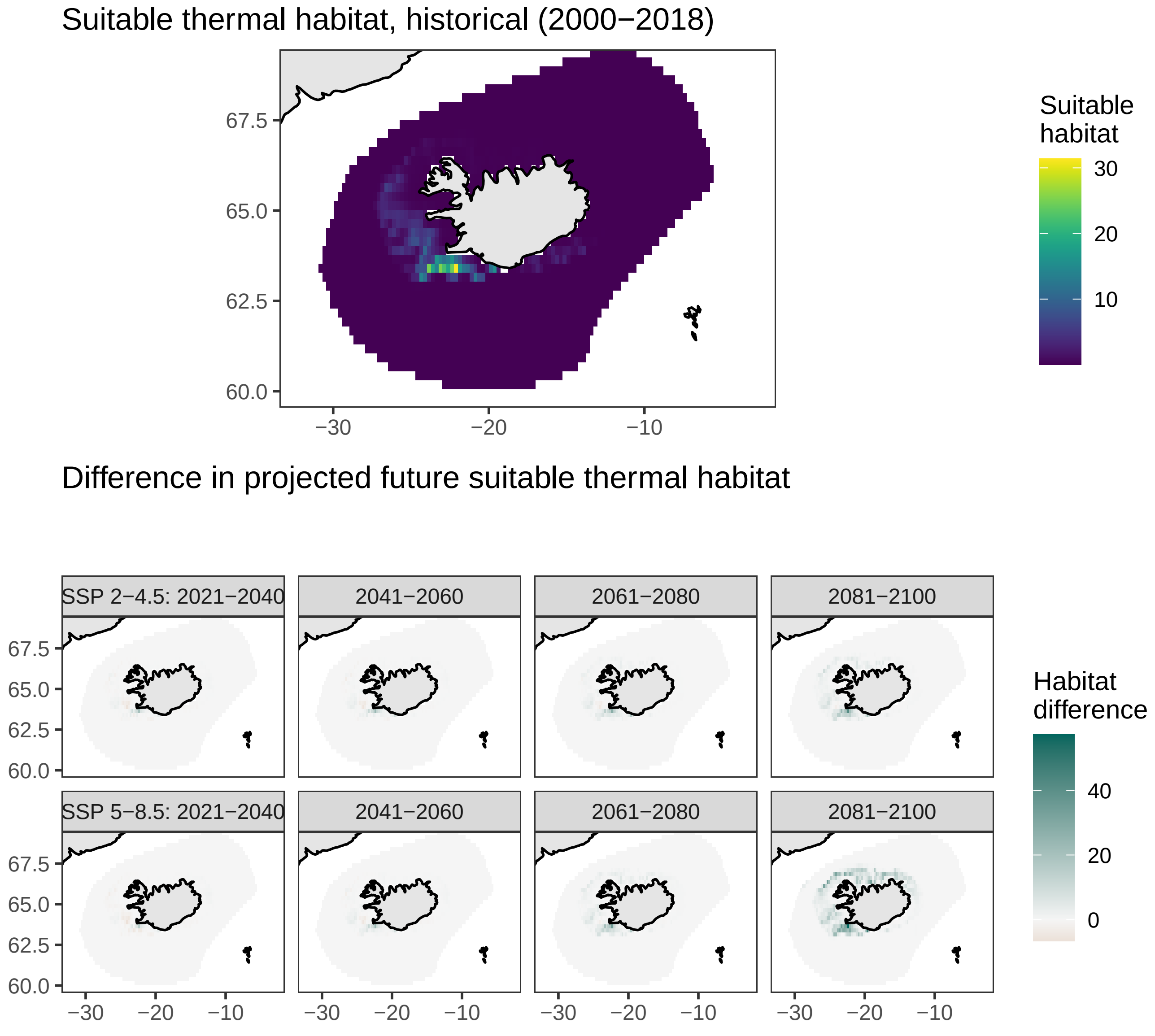
